# A dynamic, spatially periodic, micro-pattern of HES5 underlies neurogenesis in the mouse spinal cord

**DOI:** 10.1101/2020.08.03.234369

**Authors:** V Biga, J Hawley, X Soto, E Johns, D Han, H Bennett, AD Adamson, J Kursawe, P Glendinning, C.S Manning, N Papalopulu

**Affiliations:** Faculty of Biology Medicine and Health, The University of Manchester, Oxford Road, Manchester, M13 9PL, UK; Department of Mathematics, School of Natural Sciences, Faculty of Science and Engineering, The University of Manchester, Oxford Road, Manchester, M13 9PL, UK; School of Mathematics and Statistics, University of St Andrews, St Andrews, KY16 9SS, UK

**Keywords:** Neurogenesis, HES5, gene expression, oscillations, developmental patterning

## Abstract

Ultradian oscillations of HES Transcription Factors (TFs) at the single cell level, enable cell state transitions. However, the tissue level organisation of HES5 dynamics in neurogenesis is unknown. Here, we analyse the expression of HES5 ex-vivo in the developing mouse ventral spinal cord and identify microclusters of 4-6 cells with positively correlated HES5 level and ultradian dynamics. These microclusters are spatially periodic along the dorsoventral axis and temporally dynamic, alternating between high and low expression with a supra-ultradian persistence time. We show that Notch signaling is required for temporal dynamics but not the spatial periodicity of HES5. Few Neurogenin-2 cells are observed per cluster, irrespective of high or low state, suggesting that the microcluster organization of HES5 enables the stable selection of differentiating cells. Computational modelling predicts that different cell coupling strengths underlie the HES5 spatial patterns and rate of differentiation, which is consistent with comparison between the motoneuron and interneuron progenitor domains. Our work shows a previously unrecognised spatiotemporal organisation of neurogenesis, emergent at the tissue level from the synthesis of single cell dynamics.

**Synopsis:** Live imaging of HES5 expression in the ventral mouse spinal cord together with computational modelling is used to identify and analyse spatially periodic HES5 micro-patterns that emerge from the synthesis of single cell dynamics.

- HES5 is expressed in spatially periodic microclusters along the dorsal-ventral axis in spinal cord that are dynamically maintained by Notch signalling.
- Microclusters can arise, in part, from single cell oscillators that are synchronous and weakly coupled via Notch.
- Spatial patterns are different between motorneuron and interneuron progenitor domains and the probability for progenitor differentiation is regulated by the coupling strength between cells.
- NGN2 is also spatially periodic along the dorso-ventral axis and microclusters of HES5 may act to pick a single NGN2 high cell for differentiation.

## Introduction

Neurogenesis is the developmental process which generates the variety of neuronal cell types that mediate the function of the nervous system. Neurogenesis takes place over a period of days during mouse embryogenesis, thus, the transition from progenitor maintenance to differentiation needs to be balanced for development to occur normally. Neurogenesis relies on the integration of positional information with the transcriptional programme of neuronal differentiation. In the spinal cord, notable progress has been made in understanding the role and regulation of the dorso-ventral (D-V) positional system, that relies on secreted morphogens and transcriptional networks to generate the stereotyped array of different types of neurons along this axis (Briscoe & Small, 2015; Sagner & Briscoe, 2019). The transcriptional programme that mediates neurogenesis is also well understood in the spinal cord, particularly with the application of single cell sequencing (Delile *et al*, 2019; Paridaen & Huttner, 2014; Sagner & Briscoe, 2019).

Recent live imaging studies of cell fate decisions during neurogenesis have added a new dimension to this knowledge (Das & Storey, 2012, 2014; Manning *et al*, 2019a; Nelson *et al*, 2020; Soto *et al*, 2020; Vilas-Boas *et al*, 2011). They have shown the importance of understanding transcription factor (TF) expression dynamics in real time, including the key transcriptional basic helix-loop-helix repressors Hairy and enhancer of split (HES)1 and 5 (Bansod *et al*, 2017; Imayoshi & Kageyama, 2014; Ohtsuka *et al*, 1999), in regulating state transitions. We have previously shown that in spinal cord tissue, HES5 exhibits ultradian periodicity of 3-4h in about half of the progenitor population with the remaining progenitors showing aperiodic fluctuations (Manning *et al.*, 2019a). The percentage of cells that show oscillations rises in cells that enter the differentiation pathway; such cells show a transient phase of more coherent oscillations before the level of HES5 is downregulated in differentiated cells (Manning *et al.*, 2019a). Furthermore, our studies of a zebrafish paralogue Her6 showed that the transition from aperiodic to oscillatory expression is needed for neuronal differentiation, suggesting that oscillatory expression has an enabling role for cell state transitions (Soto *et al.*, 2020) as we have previously predicted computationally (Bonev *et al*, 2012; Goodfellow *et al*, 2014; Phillips *et al*, 2016).

Although these studies revealed an unappreciated dynamic behaviour at the level of HES TF protein expression, these live imaging studies are based on recording dynamics from sparsely distributed single cells in the tissue context. Therefore, little is known about how single cell dynamics are synthesised to tissue level dynamics. Do cells interact with their neighbours in order to coordinate their cell state transitions and if so, how and what is the mechanism?

Notch is of particular interest in this context because it is a highly conserved cell to cell signalling pathway that is well known for generating complex spatial patterns of cell fates in tissue development (Cohen *et al*, 2010; Corson *et al*, 2017; Henrique & Schweisguth, 2019; Hunter *et al*, 2016; Shaya & Sprinzak, 2011). Activation of Notch receptors by Notch ligands, including DLL1 and JAG1, results in downstream expression of HES1 and HES5. HES TFs can influence Notch activity on neighbouring cells by repressing Notch ligand expression either directly (de Lichtenberg *et al*, 2018; Kobayashi *et al*, 2009) or indirectly through the repression of proneural TFs such as Neurogenin1/2 (NGN1/2) (Ma *et al*, 1998). We argue that in order to understand how the balance of progenitor factor HES can be tipped in favour of a proneural factor giving rise to a decision point in neural progenitor cells, we need to address tissue-level patterns of HES expression and use computational models that can integrate the complexity of interactions at multiple scales.

The effects of Notch-Delta signaling combined with HES oscillations have been investigated during somitogenesis. Live imaging of dissociated PSM cells in vitro has shown that single cell oscillators can self-organise through Notch dependent synchronisation to generate waves in gene expression similar to those observed in vivo (Tsiairis & Aulehla, 2016). A model of mRNA and protein production and self-repression with transcriptional delay explains the emergence of autonomous oscillations of Her1 and Her7 as well as synchronization by Notch activity observed during the formation of somites (Lewis, 2003; Özbudak & Lewis, 2008; Webb *et al*, 2016). A more abstract Kuramoto-style model with time delays explains how a population of initially asynchronous and autonomous oscillators can evolve to adopt the same frequency and phase in order to periodically form somites (Morelli *et al*, 2009; Oates, 2020). The period of the oscillations determines the size of the somite and Notch abundance controls dynamic parameters such as the time to synchronisation (Herrgen *et al*, 2010). Apart from a limited number of studies suggesting an anti-phase relationship of DLL1 oscillations in neighbouring cells in neural cells (Shimojo *et al*, 2016), whether and how neural progenitor cells co-ordinate fate decisions and dynamic HES activity with their neighbours remains unknown.

In this study, we observe spatially periodic HES5 micro-patterns that can be generated by local synchronisation of low coherence single cell oscillators present in spinal cord tissue and these patterns are maintained in a dynamic way through Notch mediated cell-cell interactions. A computational model predicts that coupling strength changes spatial patterns of expression and in turn, the probability of progenitor differentiation. We confirm that between adjacent progenitor domains in the spinal cord, the rate of differentiation correlates with spatial patterns of HES5 and cell-cell coupling strength. Thus, organisation of neural progenitors in HES5 phase-synchronised and level-matched progenitors, is an exquisite spatiotemporal mechanism conferring tissue level regulation of the transition of single cells from neural progenitor to neuron.

## Results

### Positive correlations in Venus::HES5 intensity are indicative of microclusters in spinal cord tissue

Within the peak of spinal cord neurogenesis (E9.5-E11.5), HES5 is expressed in two broad domains in the dorsal and ventral embryonic mouse spinal cord (Manning *et al.*, 2019a; Sagner *et al*, 2018). Previously we have characterised the single cell dynamic behaviour of the more ventral HES5 expression domain that covers the ventral interneuron (p0/p2) and motorneuron progenitors (p2/pMN) (Manning *et al.*, 2019a). Thus, to understand how the single cell expression dynamics contributes to tissue level behaviour we have focused here on the same ventral area of HES5 expression (**Figure 1** and **Figure EV1a**). In this area, all progenitor cells (marked by SOX2) show HES5 expression (**Figure EV1b).** To characterise the spatial pattern of HES5 protein expression in this progenitor domain we made ex-vivo slices of E9.5-E11.5 Venus::HES5 knock-in mouse embryo spinal cord (Imayoshi *et al*, 2013). In snapshot images of this domain we noticed multiple local clusters of neural progenitor cells with similar levels of nuclear HES5 (**Figure 1a**) which we refer to as ‘microclusters’. These are notable after manual segmentation using a Draq5 live nuclear stain and averaging HES5 intensity across the nucleus (**Figure 1a-c**). The differences in Venus::HES5 intensity between nuclei did not correlate with the Draq5 nuclear staining indicating this was not related to global effects or effects of imaging through tissue (**Figure EV1c**). By measuring the number of nuclei in microclusters with high Venus::HES5 levels we found that they consisted of 3-4 cells wide in the apical-basal (A-B) and 2-4 cells wide in the dorso-ventral (D-V) direction (3-7 cells in total) and were similar in size between E9.5 and E11.5 (**Figure 1d**). Consistent with the presence of microclusters of cells with similar levels, nuclei showed a positive correlation in Venus::HES5 between neighbours 1 to 4 (**Figure 1e**). We took a more quantitative approach and correlated mean nuclear HES5 levels between all pairs of nuclei and found that nuclei close to each other were highly positively correlated and this correlation dropped with increasing distance, becoming negative at distances over 50 μm (**Figure 1f).** This relationship was similar across E9.5-E11.5 (**Figure EV1d)** and substantially different to the correlation coefficients calculated from randomisations of the nuclei intensities but keeping the same nuclear spatial arrangement (**Figure 1f)** which indicates the presence of a pattern in HES5 levels.

**Figure 1.**
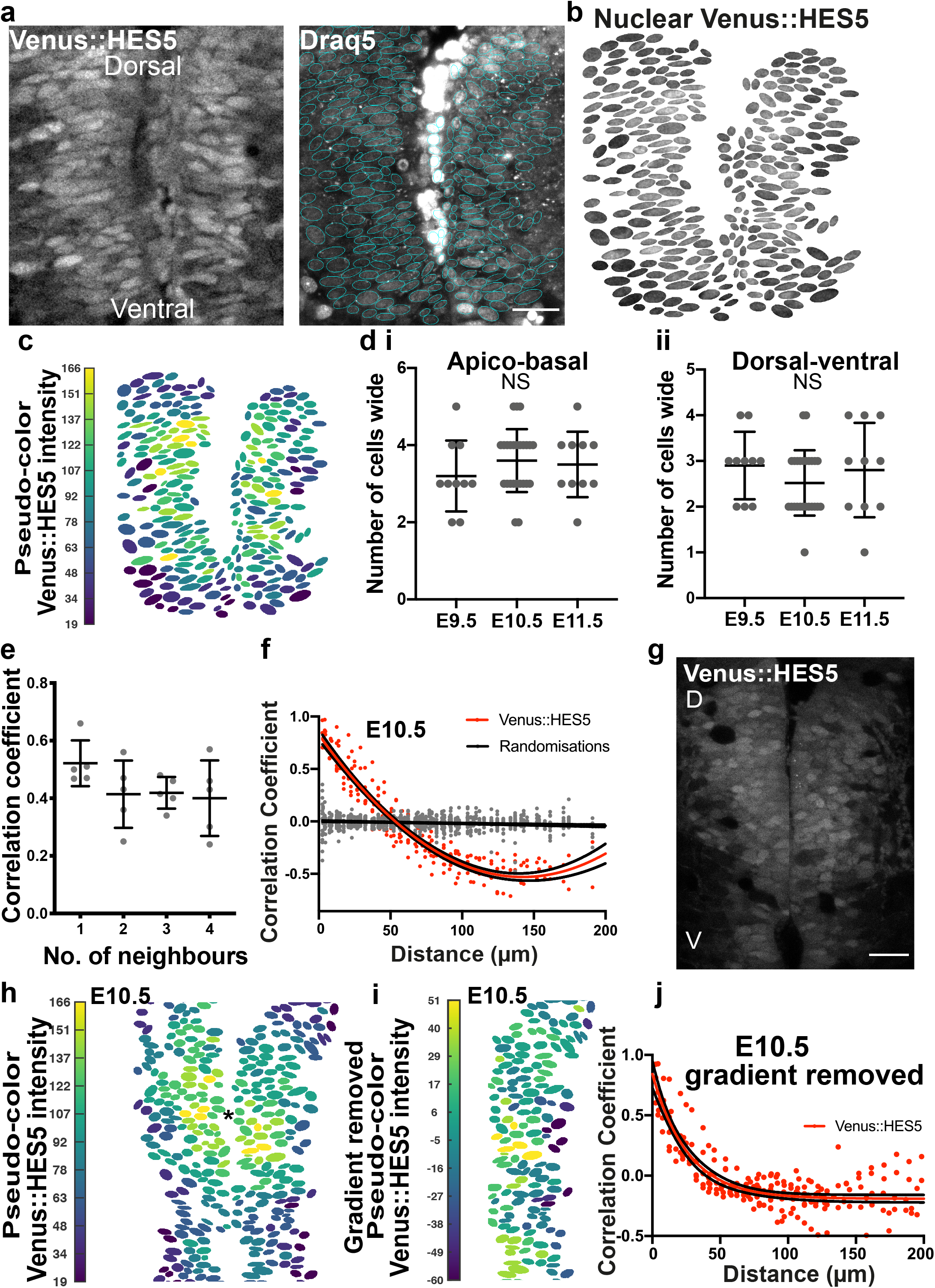
Microclusters of spinal cord neural progenitor cells have positively correlated HES5 levels. **(a)** Transverse slice of live E10.5 Venus::HES5 homozygous knock-in mouse showing the ventral HES5 domain in spinal cord ex vivo (left panel); Draq5 live nuclear stain with nuclear segmentation overlay (right panel); scale bar 30 μm. **(b)** Venus::HES5 nuclear signal corresponding to tissue in **(a)** obtained by applying nuclear segmentation onto Venus channel. **(c)** Pseudo-color look-up table applied to mean nuclear Venus::HES5 intensity (Materials and Methods) corresponding to segmented image in **(b).** Black line surrounding representative microclusters. **(d)** Dimensions of microclusters in cell numbers with high and similar levels of HES5 in i) apical-basal axes and ii) dorso-ventral at E9.5 (10 microclusters, 3 slices, 3 exps), E10.5 (10 microclusters, 9 slices, 3 exps) and E11.5 (10 microclusters, 3 slices, 3 exps). NS – no significant difference in one-way ANOVA p=0.46 (A-B), p=0.38 (D-V). **(e)** Pearson correlation coefficient observed in segmented E10.5 homozygous Venus::HES5 spinal cord ex-vivo slices showing correlation between mean nuclear Venus::HES5 intensity in any cell compared to its nearest neighbours (see Methods); dots indicate average per slice; bars indicate mean and standard deviation of 5 slices from 3 experiments (dataset is different from **(d)**. **(f)** Pearson correlation coefficient of mean nuclear Venus::HES5 intensity in relationship to distance; red dots indicate average Venus::HES5 correlation per slice of 12 slices from 3 experiments with corresponding red line indicating polynomial fit (order 2); gray dots with black line indicate correlations and polynomial fit from 5 randomizations of intensities analysed in the same way (see Methods). **(g)** Transverse slice of live E10.5 Venus::HES5 homozygous knock-in mouse showing the ventral HES5 domain in spinal cord ex vivo. Scale bar 30 μm, D-dorsal, V-ventral. **(h)** Pseudo-color look-up table applied to mean nuclear Venus::HES5 intensity of g); centre of intensity shown with *. **(i)** Pseudo-color look-up table applied to mean nuclear Venus::HES5 intensity in **h)** (only 1 side of ventricle) after radial gradient removal (see methods). **(j)** Pearsons correlation coefficient of mean nuclear Venus::HES5 intensity with distance after subtraction of radial gradient in Venus::HES5 intensity; red dots represent average in each of 12 slices from 3 experiments.

The longer-range negative correlations may arise from gradients in HES5 expression in A-B and D-V direction. Indeed the images indicate the presence of a radial gradient emanating from an area of highly expressing cells (**Figure 1g,h and Figure EV1e,f)**. Such a radial gradient could be due to the downregulation of HES5 as cells differentiate and move basally from the progenitor domain as well as to D-V differences in the level of expression (see below), and is not further investigated here. To ask whether the local positive correlations in HES5 levels are an artefact of this larger-scale domain expression pattern, we measured and subsequently removed a radial gradient across the tissue from the segmented single cell images (see Materials and Methods). However even after removing a radial gradient, mean nuclear HES5 levels at E9.5-E11.5 remained highly positively correlated at distances less than 40-50 μms (**Figure 1i** and. **Figure EV1g,h**). Therefore, a global tissue gradient of HES5 cannot fully explain the detailed spatial pattern and further factors, such as microclusters of cells with similar HES5 levels, must contribute to the formation of the HES5 spatial pattern.

### HES5 microclusters are spatially periodic along dorso-ventral axis of spinal cord

The high-resolution analysis of single-cell snapshots showed the presence of multiple microclusters in HES5 expression in the ventral domain. Next, we asked whether these microclusters have a regular spatial arrangement. To do this we drew line profiles 15 μm wide, parallel to the ventricle, in the ventral to dorsal direction **(Figure 2a** and **Figure EV2a,b)** and plotted the Venus::HES5 intensity along this line (**Figure 2b)** from lower resolution 20x images of ex-vivo slice cultures. Throughout the paper the 0 distance is the ventral most point of the HES5 domain, and distance extends dorsally (Materials and Methods). Detrending the signal removed a bell-shaped curve of expression that is a result of different HES5 levels along the D-V axis and is not further investigated here (**Figure 2b** and **Figure EV2c**). The microclusters could be detected as peaks in the detrended Venus::HES5 intensity profile (direct comparison in **Figure EV2a-c**) and we observed multiple peaks across the D-V axis (**Figure 2c**). Periodicity analysis of the detrended spatial Venus::HES5 expression profile revealed the presence of significant peaks in the power spectrum (**Figure 2d**) and multiple significant peaks in the auto-correlation function in all tissues analysed (**Figure 2e, Appendix Figure S1a** and **Appendix Table S1**). A significant peak in the auto-correlation function shows the signal has similarity to itself at the lag indicated in the x-axis. Multiple peaks in an autocorrelation function are indicative of a periodic signal with the peak-to-peak distance in the autocorrelation corresponding to the period of the signal. The spatial period in Venus::HES5 expression was 30 μm to 60 μm with a median period of 40 μm and no significant difference between E9.5 to E11.5. Periodicity measurements from auto-correlation functions and the power spectrum corresponded well (**Figure 2f** and **Appendix Figure S1b**).

**Figure 2.**
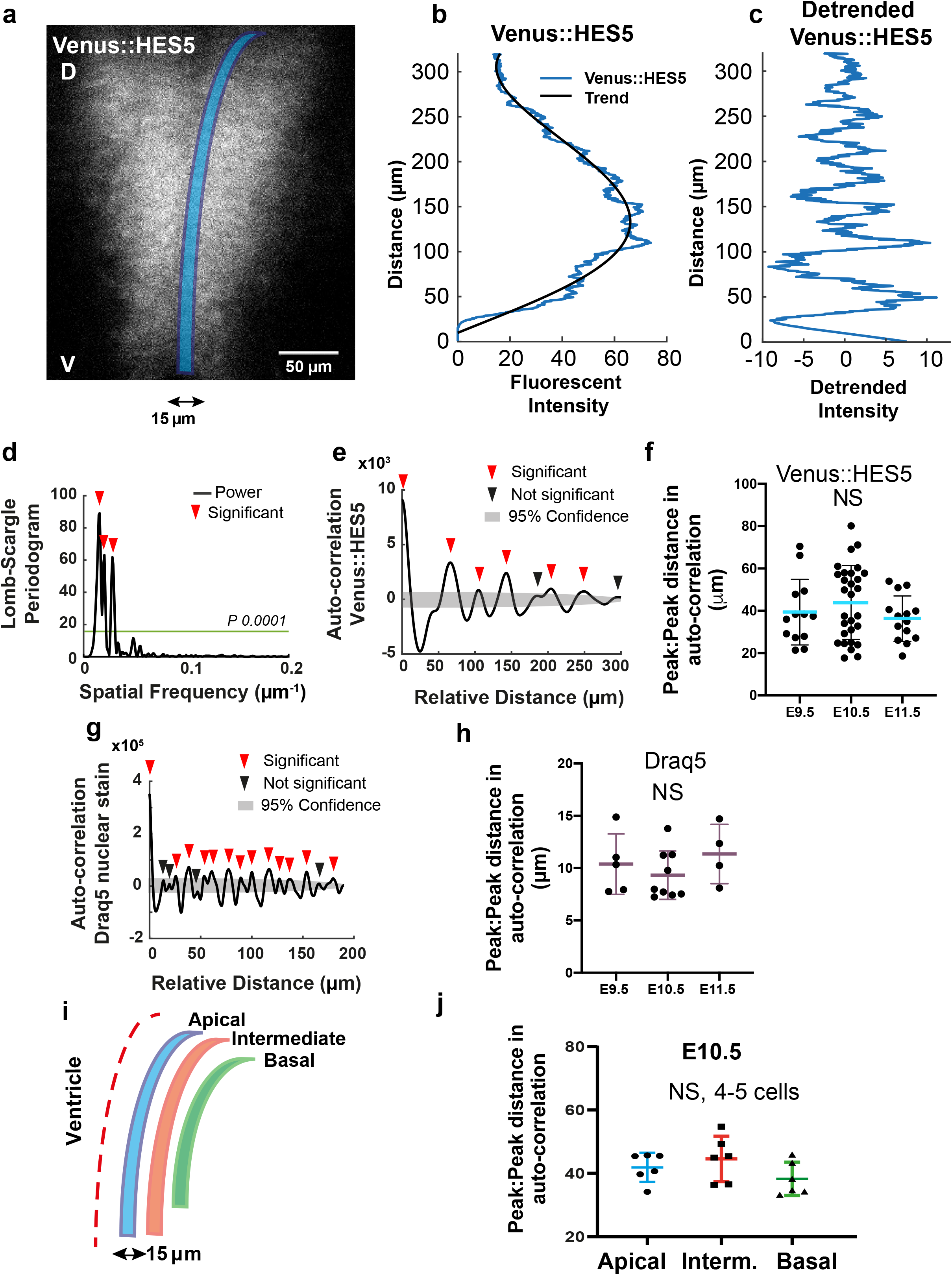
HES5 microclusters are spatially periodic along dorsal-ventral axis of spinal cord. **(a)** 20x snapshot of an ex-vivo slice culture of E10.5 spinal cord from Venus::HES5 heterozygous knock-in mouse, transverse section; delineated region (blue) correspond to data shown in (**b-c**). (**b)** Spatial profile of Venus::HES5 intensity averaged over 2.5hrs with 0 distance representing the ventral end of kymograph; black line represents the trend in Venus::HES5 data across the domain produced using an polynomial order 6 (see Methods). **(c)** Detrended spatial profile of Venus::HES5 corresponding to data shown in **(b)**. **(d)** Lomb-Scargle Periodogram analysis of detrended Venus::HES5 data in (**c**); horizontal line indicates significance level p=0.0001; red arrowhead indicate significant peaks. **(e)** Auto-correlation analysis of detrended Venus::HES5 spatial profile in **(c)** with multiple peaks indicating spatial periodicity; significant peaks (red arrowhead) lie outside gray area indicating 95% significance based on bootstrap approach (see Methods) and non-significant peaks (black arrowhead). **(f)** Peak to peak distance in auto-correlation from detrended Venus::HES5 signal collected in apical regions of spinal cord between E9.5-E11.5; bars indicate mean and SD of individual slices from 3 independent experiments; Kruskal-Wallis test not significant, p=0.44. **(g)** Representative example of auto-correlation from detrended Draq5 nuclear signal with peak to peak distances indicative of inter-nuclear distance in live tissue; gray area denotes 95% confidence area for Draq5. **(h)** Peak to peak distance in auto-correlation of detrended Draq5 spatial profile in apical regions of spinal cord between E9.5-E11.5; bars indicate mean and SD of individual slices from 3 independent experiments; Kruskal-Wallis test not significant, p=0.3. **(i)** Schematic of multiple non-overlapping regions of interest identified as Apical, Intermediate and Basal in the spinal cord tissue; width of regions in the apical-to-basal direction was 15μm. **(j)** Peak to peak distance in auto-correlation of detrended Venus::HES5 spatial profile corresponding to Apical, Intermediate and Basal regions of spinal cord at E10.5; dataset is different from **(h)**; markers indicate average distance per experiment with a minimum of 3 z-stacks per experiment and 2 repeats (left and right of ventricle) analysed per z-stack; bars indicate mean and SD of 6 independent experiments; Kruskal-Wallis test not significant, p=0.115; distances correspond to 4-5 cells considering the inter-nuclear distance in DV quantified in **(h)**.

To understand how our observed spatial periodicity relates to nuclear count in the tissue, we analysed the spatial profile for the Draq5 nuclear stain from snapshot images of Venus::HES5 ex-vivo slices. We observed peaks in Draq5 in regions of low Venus::HES5 indicating that the lower Venus::HES5 regions did not correspond to a lack of nuclei at this position of the tissue (**Figure EV2d**). As expected, the Draq5 signal observed along the D-V axis also showed multiple significant peaks in the autocorrelation that corresponded to a spatial period of 10 μm, a single-cell width and was consistent over developmental time (**Figure 2g,h, Figure EV2e** and **Appendix Figure S1c**). Using this value we estimate that the periodic occurrence of microclusters of cells with correlated levels of Venus::HES5 has a median period of 4 cells.

Since the apical region of spinal cord contains proliferative neural progenitors with high levels of HES5 that become downregulated when progenitors begin to migrate towards basal regions, we interrogated if the spatial periodicity persisted in the A-B axis. Venus::HES5 expression profiles collected from apical, intermediate and basal regions (**Figure 2i**) within the HES5 expression domain at E10.5 all showed spatial periodicity (both power spectrum and auto-correlation, **Figure EV2f**) with the period varying from approximately 4 cells in the apical side to 3 cells in the basal region (**Figure 2j, Appendix Figure S1d**). These results suggest that proliferative progenitors (localised apically) as well as differentiating progenitors (localised more basally) show local spatial correlation in Venus::HES5 levels between neighbouring cells where 3-7 neighbouring cells can be in a high or low state in synchrony with each other and that these clusters are repeated periodically in the D-V axis.

To test whether clusters extended in the anterior-posterior (A-P) axis we took longitudinal cryo-sections of the spinal cord and performed auto-correlations of the Venus::HES5 spatial profile along the A-P axis (**Figure EV2g,h)**. Peaks in the auto-correlation show spatial periodicity in A-P axis of around 30um (**Figure EV2i**). Thus, the scale of cluster size in A-P is comparable to that observed in D-V. We confirmed this in our existing kymograph data by correlating the expression of HES5 at subsequent z-positions extending in the A-P axis in the same slice (Materials and Methods). Indeed, correlations in A-P persisted at less than 30um but were lost further away (**Figure EV2j**).

The microclusters could be set up earlier on in development with fewer or single cells, and then clonally expand through cell division. However the similar microcluster size and Venus::HES5 spatial periodicity between E9.5, E10.5, and E11.5 argues against a clonal expansion mechanism. Co-ordinated cell behaviours such as nuclear-motility may also contribute. We found weak positive correlation in the movement of nuclei in apico-basal axis between cell pairs less than 30μm apart, but there was a large variation in correlations, and the correlation dropped between cells further apart (**Appendix Figure S1e**). This weak correlation in apical-basal nuclear movement may contribute weakly to maintaining microcluster pattern.

### The HES5 spatial pattern is dynamic over time

We next investigated whether the spatially periodic pattern in Venus::HES5 is also dynamic over time. To do this, we generated kymographs, single images that represent spatial intensity profiles in the same region of tissue over time, from 15 μm wide ventral-dorsal lines in movies of E10.5 Venus::HES5 spinal cord ex vivo slices **(Figure 3a,b**, **Appendix Figure S2a,b** and **Appendix Movie S1**). We noticed stripes in the kymograph, corresponding to the spatially periodic Venus::HES5 pattern (**Figure 3b**). To investigate how long high HES5 and low HES5 clusters persist over time we split the kymograph into adjacent 20μm regions (half of the 40μm spatial periodicity, chosen to capture the size of a microcluster) along the D-V axis and followed their levels over time (Methods). Hierarchical clustering of the dynamic behaviour of the kymograph regions revealed changes from low to high Venus::HES5, high to low, or re-occurring high-low-high, showing that clusters of cells can interconvert between low and high HES5 states (**Figure 3c** and additional examples **Appendix Figure S2c**). To exclude the possibility of sample drift in the DV axis being responsible for these dynamics, we used single cell tracking from the same videos as the kymographs to determine global DV drift is minimal (<20um per 12h, Appendix Figure S2) and only one in 10 tissues was excluded from temporal analysis. Thus we could proceed to analyse the persistence of a cluster in the high or low state and we found that it was on average 6-8 hours with no significant difference between persistence of high or low states in the same region (**Figure 3d,e**). We confirmed these results using a second method that detected high/low regions in the first 2 hrs of kymograph and fixing ROIs around these regions whereby we continued to observe changes in intensity over time (**Appendix Figure S2d,e**). This shows that the micro-stripes of HES5 expression are not stable but are dynamic over time.

**Figure 3.**
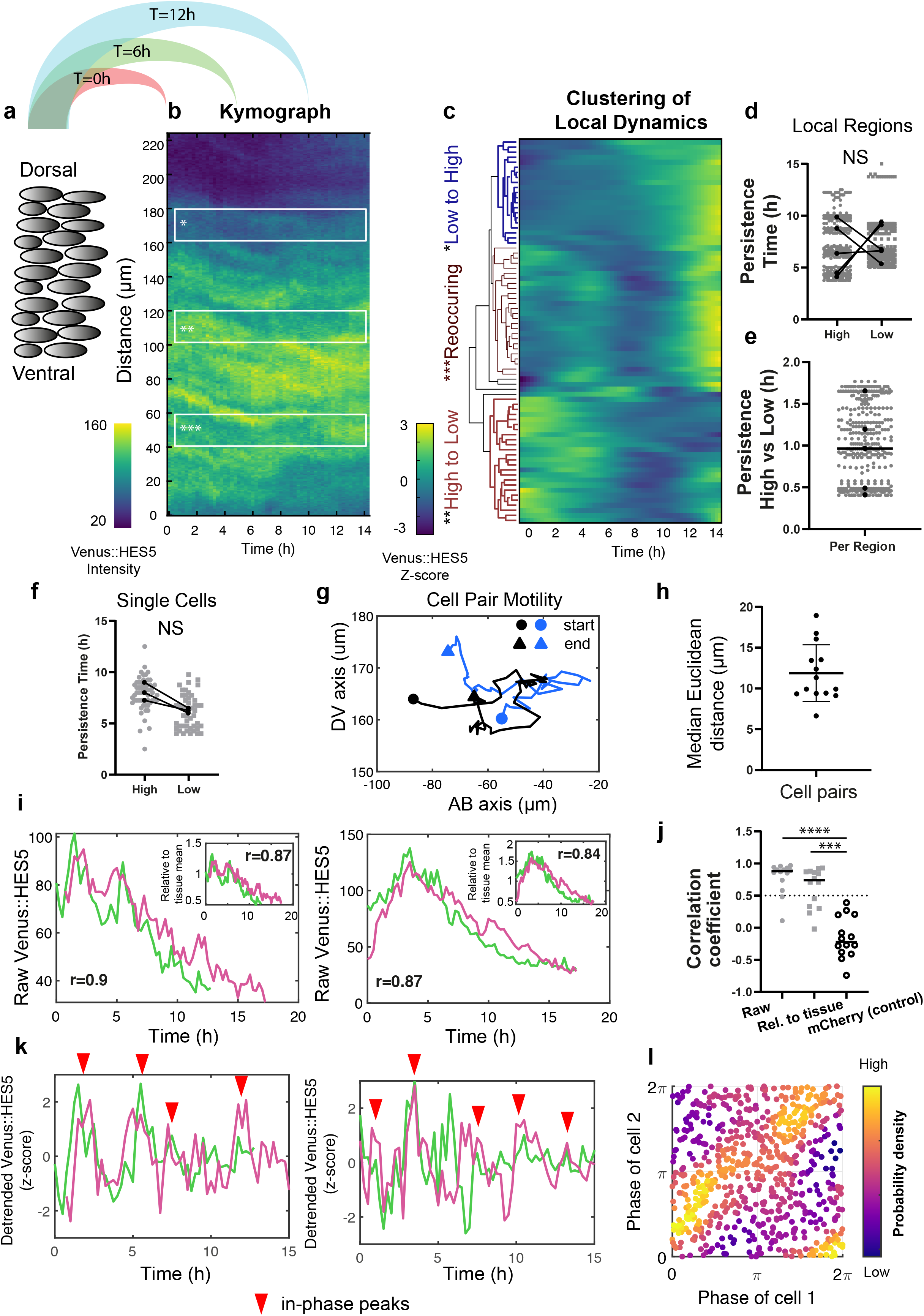
HES5 protein is expressed in a dynamical spatial periodic pattern modulated by Notch. **(a)** Schematic of extracting kymograph information from tissue data by averaging Venus::HES5 intensities observed in E10.5 heterozygous spinal cord slices to generate one intensity profile in the dorsal-ventral axis per timepoint (see Methods). **(b)** Representative kymograph data showing spatiotemporal Venus::HES5 expression profile along ventral-dorsal direction in a 15μm wide apical region and observed over 14h; local bands of 20μm width in D-V; region of interest markers indicate: *low to high, **high to low and ***re-occurring high/low activity in the same area. **(c)** Hierarchical clustering of apical Venus::HE5 expression from one representative experiment showing behaviour in the same area over time; columns represent fluctuations in Venus::HES5 intensity in small local areas (bands) obtained by dividing the spatial signal into non-overlapping 20μm regions and normalizing to the mean and standard deviation of each region over time (z-scoring); data has been subject to a Gaussian blur pre-processing step (see **Appendix Figure S2b** and Methods). **(d)** Persistence of Venus::HES5 in 20μm regions expressed as continuous time intervals when signal in the band is high or low compared to its mean (see Methods); individual datapoints (gray) indicate quantification of high and low persistence time obtained from over 300 thin bands collected from multiple tissues with 2 z-stacks per tissue and 2 repeats (left and right of ventricle) per z-stack; dots indicate paired medians of 5 independent experiments; statistical test is paired t test of median per experiment with two tail significance and p=0.7171. **(e)** Ratio measure indicating time interval at high HES5 over time interval spent in low HES5 observed in apical 20μm bands; markers indicate ratios collected from over 300 bands also shown in **(d)**; black dots indicate the median of 5 separate experiments and the black line indicates the median of all experiments. **(f)** Persistence of Venus::HES5 levels in high and low states taken from tracked single cells collected from 3 independent experiments; paired t test not significant p=0.0533. **(g)** Positional data projected in the DV and AB axis obtained from 3D tracking of a cell pair showing cells stay in close proximity over 12-15h. **(h)** Relative distance between cell pairs computed from relative 3D Euclidean distance between nuclei over 12-15h; dots indicate median distance over tracking period, horizontal lines show mean and SD of 14 cell pairs from 3 experiments. **(i)** Representative example timeseries of raw Venus::HES5 raw intensity in cells pairs identified as remaining in close proximity; insert shows Venus::HES5 intensity relative to the tissue mean intensity; r-values indicate Spearman correlation coefficients between time traces over all co-existing timepoints. **(j)** Spearman correlation coefficients computed in the same cell pairs from Venus::HES5 (Raw), Venus::HES5 relative to the tissue mean (Rel. to tissue) and H2B:mCherry (control) nuclear intensity timeseries; dots indicate correlation coefficient of pairs, lines show median from 14 cells from 3 experiments; Kruskal-Wallis test with Dunn multiple comparison with significance p<0.001*** and p<0.0001***. **(k)** Detrended Venus::HES5 fluorescent intensity timeseries (after z-scoring) corresponding to examples in **(i)**; red markers indicate in-phase peaks. **(l)** Density phase plots from instantaneous Hilbert phase reconstruction at multiple timepoints over a 12-14h period; dots indicate the phase angle in Cell 1 and Cell 2 from 14 pairs collected from 3 experiments; colormap indicates probability density showing accumulation of phase values predominantly along the (0,0) and (2π, 2π) diagonal; light colours indicate most frequent.

Since the HES5 expression is periodic along the D-V axis, it can be represented as a spatial oscillator. Therefore, we used its phase characteristics denoting the position in the spatial cycle, to analyse how the HES5 signal changes from high to low in the same region over time. We transformed the detrended spatial Venus::HES5 intensity (**Figure EV2k, Appendix Figure S3a**) along the D-V axis to phase of the spatial oscillator of Venus::HES5 using the Hilbert transform (Methods). All experiments showed a dynamic pattern with changes in phase in any area of the tissue over the 12 to 24 hr movies (**Appendix Figure S3b**). Regions could be identified that maintained a similar phase over several hours followed by a change, indicating a switch in state of the Venus::HES5 pattern (**Figure EV2k**). Phase waves could be observed in some movies, indicated by the diagonal lines of similar colours in the spatial phase map (**Appendix Fig S3biii**), however these were variable across the data and did not have a consistent direction in the D-V axis between experiments. In summary, we find microclusters of cells with correlated Venus::HES5 levels that are a maximum of 2-3 cells wide in D-V and 3-4 in A-B axes and are arranged in a spatially periodic pattern. The pattern is also temporally dynamic with a persistence in a high or low level expression of 6-8 hrs but no consistent phase wave travel in D-V.

### Correlations in Venus::HES5 levels over time and local co-ordination in phase is observed in single cells in tissue

We next addressed how the dynamic tissue pattern relates to single cell behaviour and Venus::HES5 expression. We previously tracked sparse single nuclei labelled with inducible Cre-mediated H2B::mCherry in E10.5 Venus::HES5 spinal cord ex vivo slices and reported that about half of the progenitors show oscillations of period 3-4 hrs, while others show aperiodic fluctuations (Manning *et al*, 2019b). However, the persistence of HES5 high or low microclusters in the spatial pattern is around 2x the period of Venus::HES5 temporal oscillations. This supra-ultradian behaviour that emerged from the tissue level analysis has not been reported before but we hypothesised that it may correspond to “longer-term” trends in the Venus::HES5 mean level (**Appendix Figure S4a**). These have been previously observed at the single cell level, modulating the mean level of the ultradian oscillations/fluctuations, but not analysed further. Thus, we re-analysed single cell Venus::HES5 expression data using the same methods to analyse persistence and found similar values in persistence time between microclusters and single cell long-term Venus::HES5 trends (**Figure 3f**). This suggests that changes in the levels of Venus::HES5 in microclusters are produced at single cell level through changes in the HES5 trend indicative of mean abundance levels.

We next identified single cell pairs that maintained close proximity (<20μm) over 12 hrs (**Figure 3g,h**) and we refer to these as cell pairs. We found that 13/14 cell pairs showed a high positive correlation (**Figure 3j:** raw≥0.5, median 0.88 and examples **Figure 3i**) in their mean Venus::HES5 levels which persisted in over 70% of pairs after correction to mean tissue fluorescence to account for photobleaching (**Figure 3j:** rel. to tissue≥0.5, median 0.74 and examples **Figure 3i** insert). The experimental control of nuclear H2B:mCherry in the same cells showed no strong correlation (**Figure 3i**, control <0.5, median −0.2). Furthermore, the cell pairs persistently showed in-phase Venus::HES5 peaks (examples **Figure 3k** red arrowhead) and phase visualisation maps of all pairs exhibited a large accumulation of Venus::HES5 instantaneous phases along the diagonal between (0,2π) and (2π,2π) indicating prevalence of local in-phase behaviour at single cell level (**Figure 3l)**. We also noted phase activity at (0, 2π) corresponding to anti-phase peaks, so we interrogated this further using cross-correlation analysis of cell pairs (**Appendix Figure S4b-d)** which indicated that the large majority of pairs show only minimal phase shift between them (**Appendix Figure S4d**, median 15 min coinciding with sampling rate). These findings demonstrate that ultradian activity between neighbouring cell pairs in spinal cord tissue is predominantly in-phase and we refer to this as ‘local in-phase’.

To summarise, inside a microcluster cells predominantly show synchronised ultradian oscillations of 3-4 hrs and each microcluster has a persistence time in a high or low state of about 6-8hrs coinciding with the persistence of single cell trends. A microcluster is a composite of these two activities.

### Notch inhibition extinguishes dynamic changes in Venus::HES5 microclusters between high and low states

We hypothesised that the periodic microclusters of HES5 are generated through Notch-Delta interactions that locally synchronise dynamic HES5 expression between neighbouring cells. To test this, we treated spinal cord slice cultures with the Notch inhibitor DBZ and performed kymograph analysis in the apical region of DMSO and DBZ treated slices. In Notch inhibitor conditions, the HES5 levels reduce continuously over time (**Figure 4a**) indicating that the DBZ is effective. The most noticeable difference in the spatiotemporal HES pattern was that the temporal transitions of microclusters from high to low Venus::HES5 were impaired by DBZ. We saw fewer changes in the phase of the spatial periodic Venus::HES5 pattern indicating the spatial pattern remained stable (**Figure 4b,c** and **Figure EV3a,b**). This was quantified with a phase synchronisation index (see Methods), where low values indicate the presence of phase changes at the same D-V locations **(Figure 4d)**. The phase synchronisation index was significantly higher in DBZ treated tissue indicating that in the absence of Notch signalling, HES5 microclusters were more persistent in the same region and that the dynamic changes in Venus::HES5 microclusters between high and low levels are mediated by Notch signalling. The phase detection method (Hilbert transform) is not dependent on the level of expression and so the reduction in HES5 levels in DBZ does not affect the analysis of microcluster high-to-low and low-to-high phase switches.

**Figure 4.**
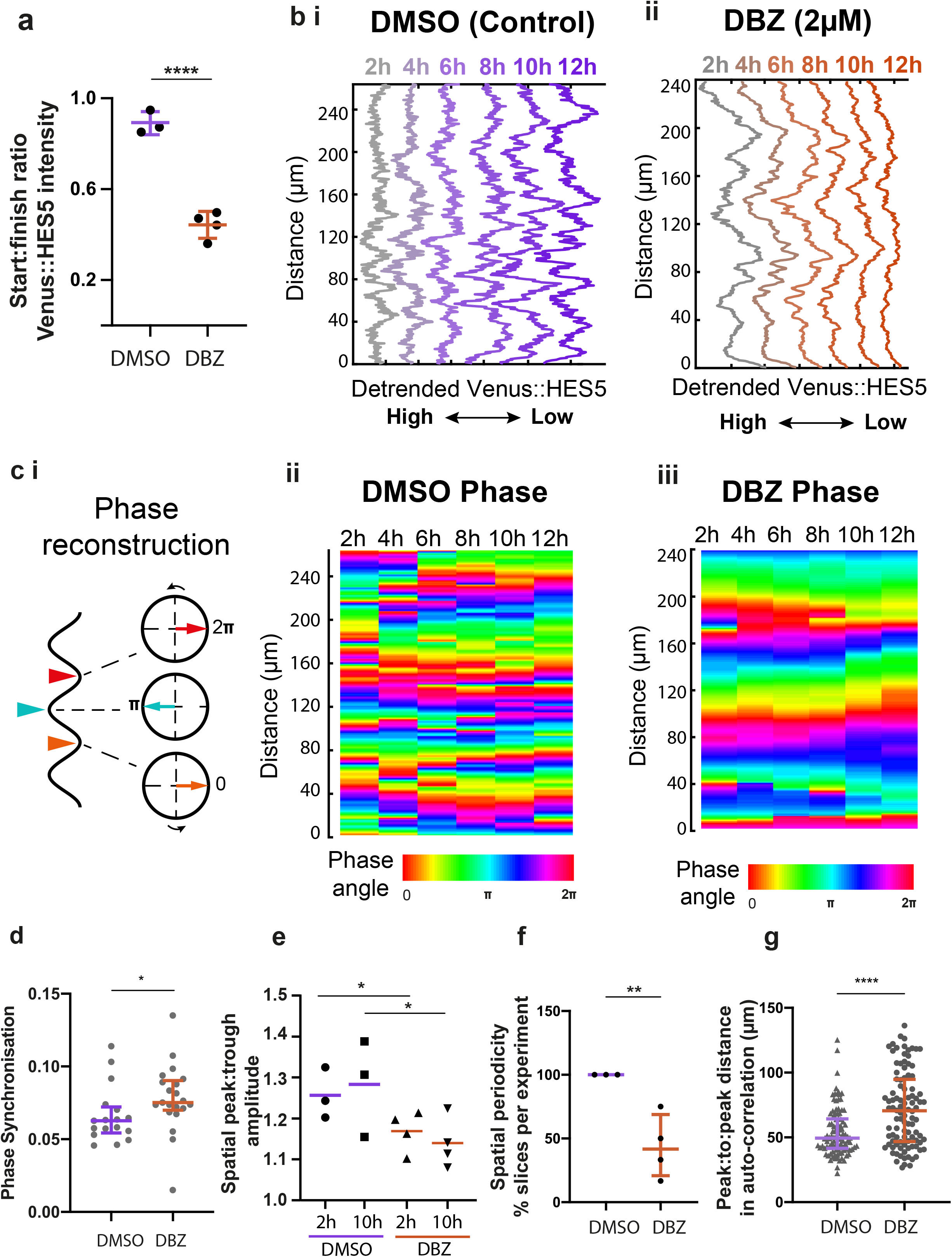
Notch inhibition increases HES5 pattern persistence. **(a)** Start:Finish Venus::HES5 intensity ratio in E10.5 Venus::HES5 heterozygous spinal cord slices treated with control (DMSO) and Notch inhibitor DBZ (2 μM) observed over 16h; bars indicate mean and standard deviation of DMS0 (n=3 experiments) and DBZ (n=4 experiments); 2-tailed t-test p=0.0001. **(b)** Representative spatiotemporal plots of the detrended Venus::HES5 pattern along ventral-dorsal direction in **(i)** DMSO control and **(ii)** DBZ conditions obtained by averaging kymographs data in the same region over 2 hr time intervals. **(c-i)** Schematic indicating the correspondence between Venus::HES5 spatial oscillator represented as detrended level and phase angle characteristics; the spatial oscillator traverses repeated cycles including start (HES5 low-orange arrowhead), middle (HES5-teal arrowhead) and end (HES5 low-red arrowhead) which in phase space corresponds to phase angles 0, π and 2π respectively. **(c-ii,iii)** Phase maps corresponding to **(ii)** DMSO and **(iii)** DBZ detrended Venus::HES5 data shown in **(b-i)** and **(b-ii)** respectively. **(d)** Phase synchronisation measure (see Methods) of the detrended Venus::HES5 spatial oscillator over time in E10.5 Venus::HES5 spinal cord slices treated in DMSO vs DBZ conditions; dots indicate DMSO (21 kymographs, n=3 experiments) and DBZ (19 kymographs, n=4 experiments); 2 tailed Mann-Whitney test p=0.0207*. (**e)** Spatial peak: trough fold change in Venus::HES5 intensity profile in the D-V axis measured at 2h and 10h in DMSO and DBZ treated E10.5 Venus::HES5 spinal cord slices; dots indicate average over 3 z-slices from DMSO (n=3) and DBZ (n=4) experiments; lines indicate median per condition; 1 tailed unpaired t test with significance p<0.05*. (**f)** Percentage of ex-vivo slices with significant spatial period detected after 10-12hrs of DMSO and DBZ conditions; significant spatial period defined as multiple significant peaks in auto-correlation detected above the 95% confidence bounds; dots indicate % per experiment; bars denote median and inter-quartile range of DMS0 (n=3) and DBZ (n=4) experiments; 1 tailed t test with significance p=0.0062**. (**d)** Peak:to:peak distance in auto-correlation plots of detrended Venus::HES5 spatial profile in DMSO and DBZ treated E10.5 Venus::HES5 spinal cord slices; gray dots represent significant mean peak-to-peak distance of DMSO (100) and DBZ (105) auto-correlation functions collected from 3 z-stacks per slice and 2 repeats (left and right of ventricle) with multiple timepoints; bars indicate median per experiments from DMSO (n=3) and DBZ (n=4) experiments; 2 tailed Mann-Whitney test p<0.0001.

We analysed the spatial periodicity of HES5 and found that the amplitude between high and low microclusters appears diminished compared to control DMSO treated conditions (**Figure 4e**). Spatial periodicity could be detected at the start of the movie, immediately after DBZ addition, however the spatial periodicity was gradually extinguished through loss of Venus::HES5 levels and spatial amplitude death (**Figure EV3c**). Approximately 45% of the DBZ treated slices did not show significant peaks in the auto-correlation of detrended spatial Venus::HES5 profile by 10-12 hrs of treatment (**Figure 4f**) whereas periodicity was maintained in all DMSO conditions. Spatial periodicity in detrended Venus::HES5 levels that could be detected in DBZ treatment at early time points frequently appeared higher in Notch inhibitor treated ex-vivo slices than in DMSO control (**Figure 4g** and **Figure EV3d**). Cell density also decreased in Notch inhibitor conditions suggesting this increase in spatial period was partially due to changes in the spatial arrangement of cells (**Figure EV3e**).

We also investigated how Notch inhibition may affect ultradian dynamics at single cell level. We had previously reported that under DBZ conditions, single neural progenitors continue to show oscillations and fluctuations in HES5 before undergoing amplitude death (Manning *et al.*, 2019b). However here we wanted to interrogate how DBZ affects the way cells co-ordinate their activity in the tissue. To do this we used the Kuramoto Order Parameter (KOP, also known as mean-field value) a population measure of synchrony (Choi *et al*, 2000). High KOP levels close to 1 are indicative of global in-phase activity whereas low KOP values close to 0 are indicative of no in-phase synchrony (see Methods). We found that KOP of single progenitors indicating weak levels of synchrony under DMSO conditions (**Appendix Figure S4e**, mean 0.36) consistent with our findings of local in-phase activity but not indicative of global synchrony. Furthermore, we observed a significant reduction of KOP values in DBZ conditions (**Appendix Figure S4e**, mean 0.15).

Taken together, these findings show that Notch signalling is responsible for certain aspects of the pattern, such as the dynamic switching between high/low HES5 microcluster states over time. However, inhibition of Notch does not seem to abolish the existence of microclusters or their spatial periodicity, as they can still be detected until amplitude death occurs and the HES5 levels are depleted. At single cell level, we observe that Notch signalling is likely to promote local in-phase ultradian co-ordination between cells within a microcluster.

### A model of Notch-Delta with HES5 auto-repression containing stochasticity and delays re-capitulates the existence of local in-phase HES5 dynamics

We used computational modelling to help us understand how positively correlated, spatially periodic, and dynamic microclusters of cells may emerge in the spinal cord. At single cell level, HES5 protein expression oscillations are due to HES5 self-repression, an intra-cellular transcriptional time delay (*τ*_*H*_) and short protein and mRNA half-lives (Jensen *et al*, 2003; Momiji & Monk, 2008; Monk, 2003). We represented the auto-repressive interactions between HES5 mRNA and protein using stochastic differential equations with time delay, as previously described in (Galla, 2009; Manning *et al.*, 2019a; Phillips *et al.*, 2016). This single cell model has been shown to faithfully recapitulate statistics of single cell HES5 expression dynamics collected from spinal cord tissue (Manning *et al.*, 2019a). We extended the single-cell mathematical description of HES5 to a coupled dynamical model by incorporating a repressive interaction in the form of a Hill function, that describes how HES5 protein in one cell represses Hes5 transcription in a neighbouring cell via Delta-Notch signalling **(Figure 5a** and **Figure EV4a)**. We introduce the following set of inter-cellular parameters (**Figure 5b** and Materials and Methods): (i) *inter-cellular time delay*, representing the time required to transfer the signal from one cell to another, that is, the time required for a change in HES5 protein in one cell to affect Hes5 transcription in a neighbouring cell through Notch-Delta; (ii) *the inter-cellular repression threshold*, representing the amount of HES5 protein required to reduce Hes5 transcription in a neighbouring cell by half; the inter-cellular repression threshold is inversely proportional to coupling strength where higher coupling strength (or low inter-cellular repression threshold) indicates that less protein is needed to repress the neighbour’s Hes5 transcription by 50%; and (iii) *inter-cellular Hill coefficient* indicating how steep the response curve of Hes5 transcription is in response to a change in HES5 protein in the neighbouring cell, with higher values corresponding to increased nonlinearity. Interactions between cells are considered in a hexagonal grid whereby each cell can interact with its immediate 6 neighbours and repression between cells is calculated through the inter-cellular Hill function by averaging HES5 protein abundance over 6 neighbours (**Figure 5b,c** and Materials and Methods). Thus, we generated a comprehensive, multiscale and stochastic model with time delays, representative of the Delta-Notch-Hes interactions in the multicellular tissue environment.

**Figure 5.**
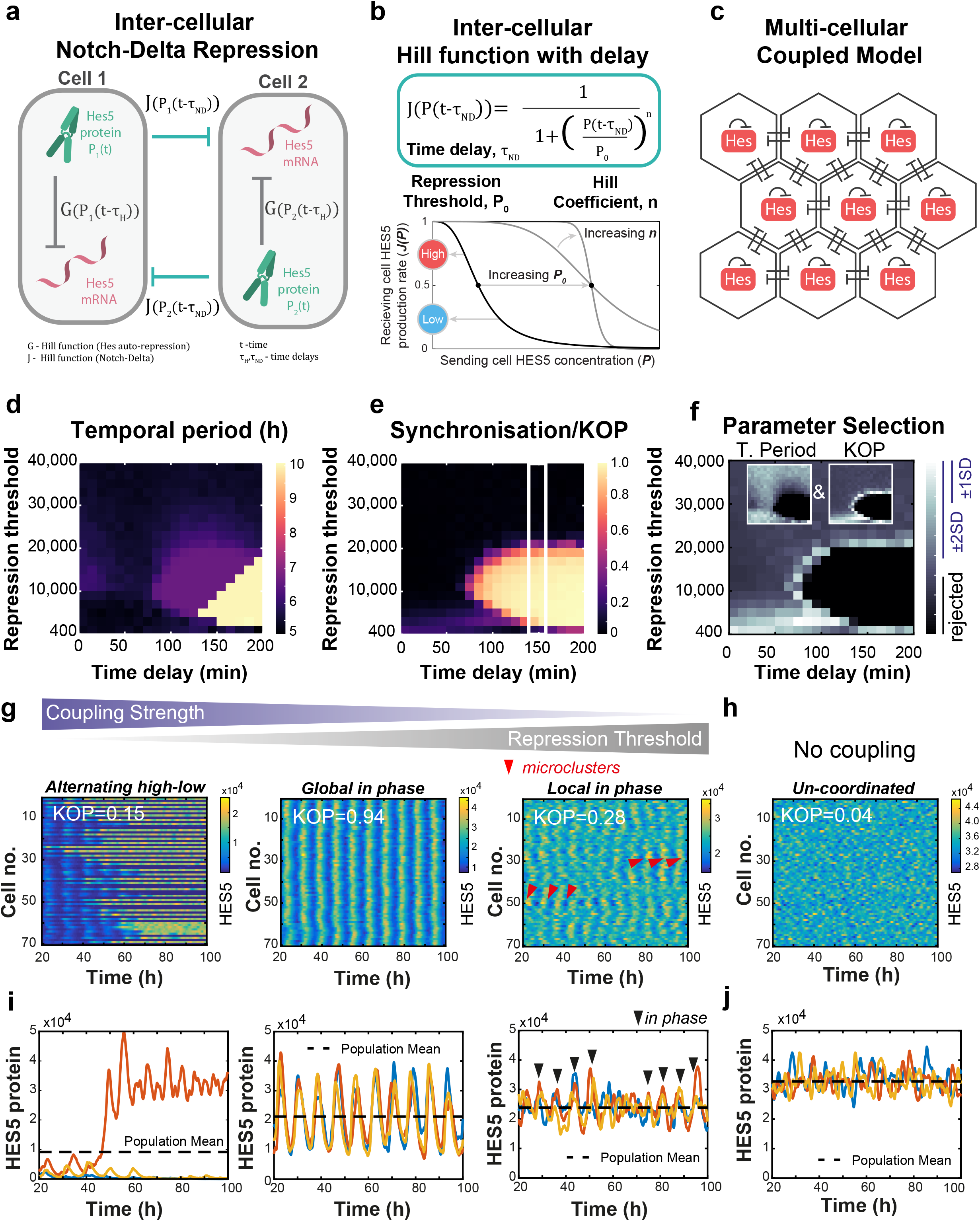
Multicellular cell-cell coupling model explains the emergence of microclusters. **(a)** Schematic of repressive interactions via Notch-Delta between neighbouring cells whereby the effects of HES5 protein in Cell 1 (marked as *P*_l_) on transcription in Cell 2 and vice-versa are represented using an inter-cellular Hill function *J*(*P*_1,2_(*t−*τ**_*ND*_)) where *t* denotes time and *τ*_*ND*_ represents the intercellular time delay, the time interval required to synthesise the intermediate molecular species (detailed in **Figure EV4a**); HES5 auto-repression is represented using an intra-cellular Hill function *G*(*P*_1,2_(*t−*τ**_*H*_)) where *τ*_*H*_ represents the inter-cellular time delay, the time interval required for protein to be produced and repress its own transcription. **(b)** Mathematical description of the inter-cellular Hill function and its parameters: time delay (*τ*_*ND*_), repression threshold (*P*_0_) and Hill coefficient (*n*); (bottom left panel) higher *P*_0_ corresponds to reduced inter-cellular repression (i.e. decreased coupling strength) and conversely lower *P*_0_ corresponds to higher coupling strength; (bottom right panel) increasing values of n correspond to increased steepness of the inter-cellular response. **(c)** Multiscale coupled mathematical model of the tissue environment consisting of a 2D hexagonal grid of cells expressing HES5 protein with corresponding auto-repression (described in **(a)**) coupled together by repressive interactions between its 6 immediate neighbours (see Methods); single cell inter-cellular repression is a Hill function (with parameters described in (**b)**) dependent on mean protein abundance in the neighbouring cells. **(d)** Parameter exploration of single cell temporal period emerging from the model at different repression threshold and time delay values. **(e)** Parameter exploration of phase synchronisation quantified using the Kuramoto Order Parameter (see Methods) where 1 indicates global in-phase activity and 0 indicates no co-ordination of phase between cells. **(f)** Parameter selection strategy combining experimentally determined temporal phase (insert left panel) and KOP (insert right panel) values in spinal cord tissue (see (Manning *et al.*, 2019a) and Methods) to indicate areas where model statistics (i.e. mean temporal period and KOP of synthetic data) resemble real tissue; values within ±1SD and 2±SD from the mean of the tissue are identified and values found outside of ±2.4SD from the mean of tissue are excluded. **(g)** Representative examples of synthetic kymograph data obtained at specific levels of repression threshold: *Alternating high-low* (*P*_0_ = 400), *Global in phase* (*P*_0_ = 15000) and *Local in phase* (*P*_0_ = 21000) and corresponding KOP values; the presence of microclusters at weak coupling is indicated with red arrowheads; time delay 150min, *n* = 4. **(h)** Kymograph data obtained in the absence of coupling between cells; phase relationships are un-coordinated resulting in a KOP≈0. **(i-j)** Synthetic data timeseries corresponding to simulations in **(g)**.

We parameterised this multiscale HES5 model with previously determined experimental measures of HES5 protein and mRNA stability and with parameter values of the single cell HES5 self-repression loop that can reproduce single neural progenitor HES5 dynamics (see Methods and **Appendix Table S3** Main), as identified through Bayesian inference in our previous work (Manning *et al.*, 2019a). We then investigated the parameter space of unknown model parameters that are characteristic of cell-to-cell interactions, namely the repression threshold (inverse of coupling strength) and time delay, to identify values that are compatible with the temporal period and phase synchronisation level of single cell Venus::HES5 expression dynamics **(Figure 5d,e)**. The mean temporal period of Venus::HES5 in ex-vivo spinal cord tissue is approx. 2-6 hrs (mean 3.3 hrs) (Manning *et al.*, 2019a), which could be reproduced by the model in a wide range of coupling strength and inter-cellular time delay values (**Figure 5d,f**). We measured the temporal phase synchronisation between single Venus::HES5 expressing cells in the apical region and we found that the KOP was between 0.15 and 0.4 (**Appendix Figure S4f**, mean 0.3) consistent with KOP in the DMSO data (**Appendix Figure S4e**, mean 0.36). This measure aided us in further reducing the parameter space of repression threshold and inter-cellular time delay that could fit the observed data (**Figure 5e,f**). The accepted parameter values for inter-cellular time delay were consistent with a Delta to HES5 transmission time of 128 min measured experimentally (Isomura *et al*, 2017). A Hill coefficient value larger than 2 was required for notable synchrony to emerge (KOP>0) and only minor differences in terms of parameter selection were observed for values between 3 and 6 (**Appendix Figure S5a**).

This parameter exploration allowed us to optimise the search for spatial patterns that emerge at different coupling strengths using kymograph analysis (**Figure 5g,h**). We set the inter-cellular time delay to 150 min and Hill Coefficient to 4 (Methods) and then compared the synthetic HES5 spatiotemporal characteristics at specific coupling strength levels (parameter space indicated by the white box in **Figure 5e**).

Our comparison showed that strong coupling (i.e. high coupling strength or low inter-cellular repression threshold) induces *Alternating high-low* dynamics whereby single neighbouring cells adopt either high oscillatory HES5 or stable low HES5 in an alternating spatial pattern that does not evolve over time (**Figure 5g**, *Alternating high-low*, **Appendix Movie S2 first panel**). Meanwhile at mid-level coupling, the multiscale model induces globally synchronised oscillations in all cells (**Figure 5g**, *Global in phase* and **Appendix Movie S2 second panel**). At weak coupling strength, the spatial patterns show areas of local synchronisation emerging between neighbouring cells (**Figure 5g**, *Local in phase* and **Appendix Movie S2 third panel**) resembling activity observed in tracked single cell pairs in experimental data (**Figure 3k**). Under no coupling conditions we observed autonomous non-synchronised stochastic oscillations and fluctuations across the tissue (**Figure 5h** and **Appendix Movie S2 fourth panel**). These observed changes in synchronisation are indicated by population KOP values (**Figure 5g**) and we further confirmed that the KOPs correspond to changes in synchrony in terms of single cell expression dynamics between neighbouring cells (**Figure 5i**). As expected, in the uncoupled cells we observed no synchrony (KOP≈0) and activity in neighbouring cells was un-coordinated over time (**Figure 5h,j**). Therefore the model can recapitulate the local in-phase behaviour in Venus::HES5 observed between single cell pairs in a microcluster.

Our explorations of synthetic data show that at weak coupling strength microclusters consisting of in-phase cells can be generated in the model with a diameter of 2-6 cells (**Appendix Figure S5b and Methods**), consistent with cluster size in spinal cord tissue. However, the occurrence rate of microclusters was low, as these were observed around 20-30% of the time, although still higher than in the uncoupled situation (**Appendix Figure S5c** and **Appendix Figure S6a**). Thus, weak coupling conditions generate microclusters by promoting in-phase activity between neighbouring cells, however these appear transiently and with low probability. In addition, the microclusters of locally synchronised cells were not spatially periodic (**Appendix Figure 5d and Appendix Figure 6b**). As expected, at high coupling (low repression threshold) we detected an alternating pattern of HES5 with a spatial periodicity of 2 cells, which is a characteristic of the classic lateral inhibition alternating high-low pattern (**Appendix Figure S5d** and **Appendix Figure S6b**).

In conclusion, our multicellular coupled model shows that spinal cord progenitors can locally synchronise at weak coupling strength to generate microclusters of 2 to 6 cells in diameter, a similar size to those seen in tissue, (**Figure 1d**) with single cell Venus::HES5 expression dynamics consistent with previous reports (Manning *et al.*, 2019a). However, the model cannot recapitulate the repeated spatial co-ordination and continuous presence of dynamic microclusters, suggesting that additional mechanisms may act in the tissue environment to stabilise their presence and promote spatially periodic emergence.

### The model predicts that probability of differentiation is regulated by the coupling strength between cells

To understand how the spatial pattern of HES5 and dynamic micropatterns in particular may affect properties of neurogenesis, we made the assumption that when HES5 is low, there is increased probability that the cell would differentiate consistent with findings that differentiation is accompanied by switching off of HES5, a repressor of neurogenesis (Bansod *et al.*, 2017; Manning *et al.*, 2019a; Sagner *et al.*, 2018). We introduced a “differentiation threshold”, which was set at the level of the HES5 population mean for each simulation (**Figure 5i**, *Population Mean*) and we reasoned that if expression level in a cell dropped below this threshold there was an increasing probability to switch off HES5 and differentiate (**Figure 6a**). We found that at high coupling strength (*Alternating high-low* conditions) the probability to differentiate is the highest, whereas medium and weak coupling strength (corresponding to *Global* and *Local in phase* synchronisation, respectively) had progressively lower probability of differentiation (**Figure 6b)**.

**Figure 6.**
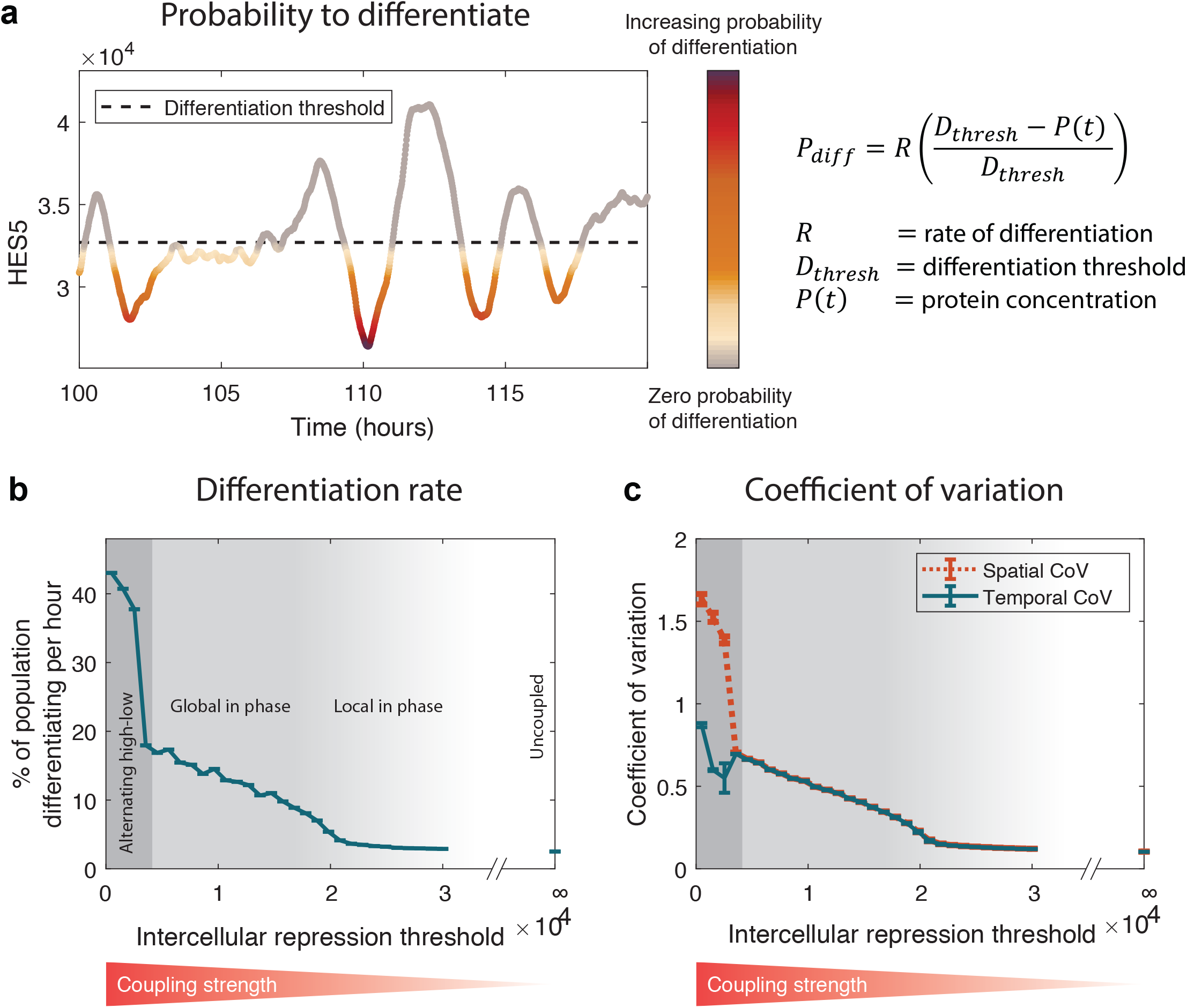
Cell-cell coupling strength can regulate probability of differentiation in a multicellular environment. **(a)** Representative synthetic timeseries example and mathematical description of probability of differentiation (*P*_*diff*_) in relation to population mean HES5 protein levels (referred to as ‘differentiation threshold’, *D*_*thresh*_) whereby HES5 protein abundance (*P(t)*) dropping below the threshold increases the rate at which cells differentiate. **(b)** Differentiation rates estimated from the multicellular coupled model (detailed in **Figure 5)** over a wide range of repression threshold values corresponding to decreasing coupling strength; three dynamic regimes are labelled as *Alternating high-low*, *Global in phase* and *Local in phase* mirroring examples shown in **Figure 5g,i**. **(c)** Analysis of temporal CoV and spatial CoV from synthetic data corresponding to differentiation rates shown in **(b)**; these statistics indicate that spatial variability correlates better with differentiation rates meanwhile temporal variability shows only a moderate quasi-linear increase in *Alternating high-low* conditions compared to the rest. Single cell parameters used to generate **(b)** and **(c)** are shown in **Appendix Table S3**-Main, and the multicellular parameters used were *n*_*ND*_ = 4, *τ*_*ND*_ = 150 minutes. Each value plotted in **(b)** and **(c)** shows the mean and SD from 10 simulations at each repression threshold value.

To understand why this is happening, we looked at the Coefficient of Variation (CoV, **Figure 6c**), a measure of variability denoting standard deviation over the mean. We investigated both the temporal (Temporal CoV) and spatial variation (Spatial CoV) in simulated HES5 expression. Indeed, both temporal (indicative of single cell amplitude) and spatial CoV (indicative of variation between HES5 high and low regions in space) appear highest in *Alternating high-low* conditions and lowest for *Local in phase* micropatterns (**Figure 6c**). However, we found that changes in spatial CoV correlated better with changes in rate of differentiation, especially at low repression threshold/high coupling strength (**Figure 6c** versus **Figure 6b***Alternating high-low*). Thus, our model predicts that the strength of cell:cell coupling may increase the probability of differentiation through amplifying cell:cell differences in abundance which in turn affects how far the cells dip below the threshold of differentiation.

### In tissue, HES5 spatial pattern varies predictably with the rate of differentiation

To test the computational prediction that the spatial pattern of HES5 (determined by the coupling strength) regulates the probability of differentiation, we compared the pattern in motorneuron and interneuron progenitor domains. We chose this comparison because at E10.5 the motorneuron domain is known to have a higher differentiation rate than the interneuron domain (Kicheva *et al*, 2014), therefore one would expect a different HES5 spatial pattern. We stained for the motorneuron progenitor marker OLIG2 (Figure 7aand Figure EV4b,c) and analysed expression levels of Venus::HES5 and Neurogenin 2 (NGN2) in the two domains. The motorneuron domain had lower HES5 levels and higher NGN2 levels than the interneuron domain (Figure 7b) consistent with the opposing activity of these genes on cell differentiation (Imayoshi & Kageyama, 2014).

**Figure 7.**
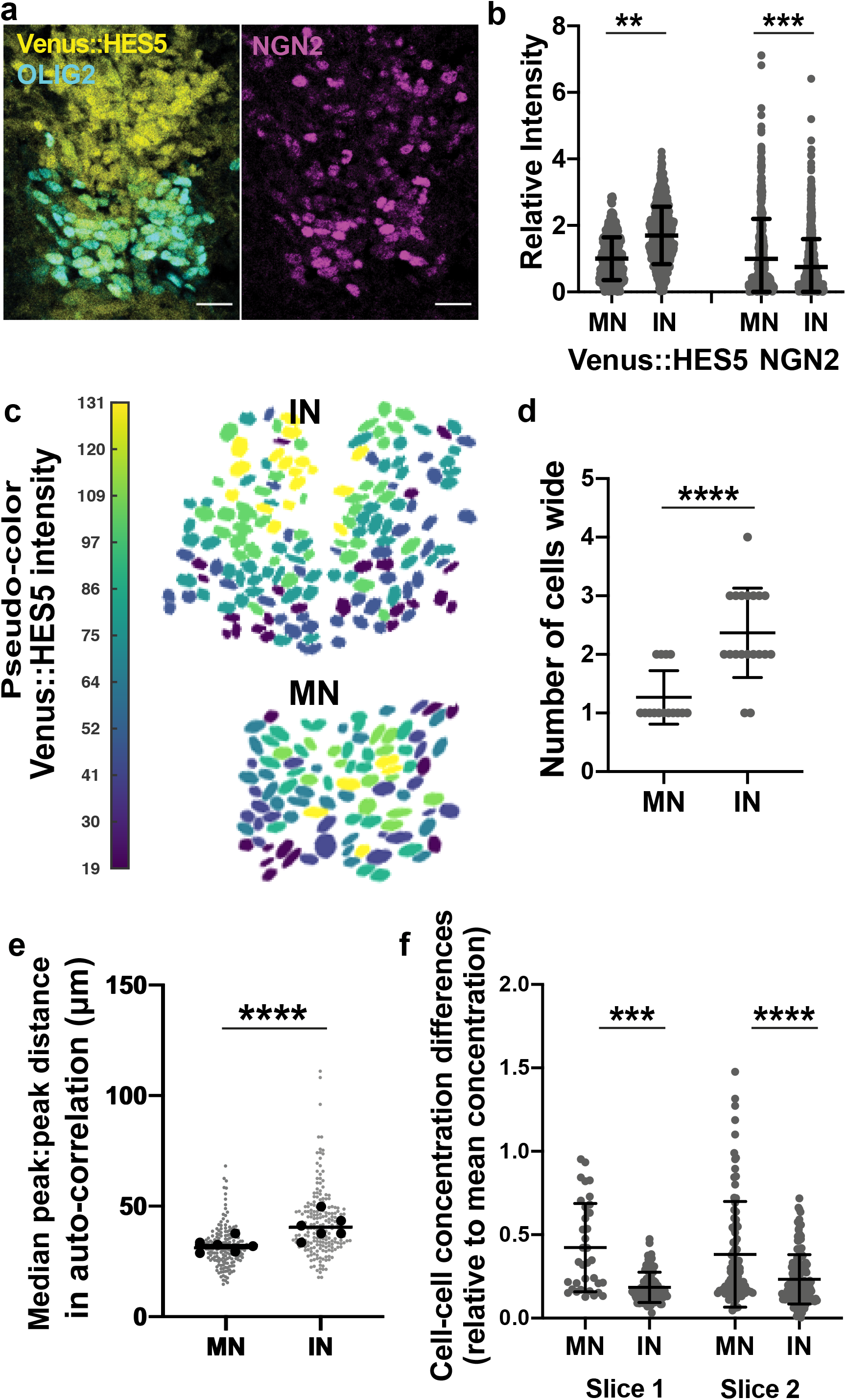
Type of HES5 spatial pattern and coupling strength correlates with rate of differentiation in motorneuron and ventral interneuron domains. **(a)** Transverse cryosection of E10.5 Venus::HES5 spinal cord. Venus::HES5 endogenous signal, OLIG2 – motorneuron progenitor marker, NGN2 – early marker of neuronal commitment; scale bar 20 μm. **(b)** Relative nuclear intensities of Venus::HES5 and NGN2 in motorneuron and interneuron progenitors; bars show mean and SD of at least 494 cells per domain from 5 slices in 2 experiments; Kruskal-Wallis with Dunn’s multiple comparison test adjusted p-values **p=0.0032, ***p<0.001. **(c)** Pseudo-color look-up table applied to mean nuclear Venus::HES5 intensity within motorneuron (MN) and interneuron (IN) domains, corresponding to segmented image in **(a). (d)** Dimension of microclusters in DV axis for MN and IN domains; microclusters counted contained cells with high and similar levels of HES5 (Methods); data consists of 34 microclusters measured from 5 sections and 3 independent experiments; 2-tailed Mann-Whitney test. Exact p-values **** p<0.0001. **(e)** Peak to peak distance in auto-correlation plots of de-trended Venus::HES5 spatial profile in MN and IN domains; this is a measure of spatial period in Venus::HES5 profile along dorsal-ventral axis of spinal cord; grey data points represent mean peak to peak distance of at least 3 slices with left and right ventricle analysed separately in 6 experiments; black dots show median per experiment and line shows overall median; 2-tailed Mann-Whitney test p-values ****p<0.00001. **(f)** Cell-cell concentration differences in HES5 between neighbours, normalised to mean concentration of HES5 in that domain; grey data points represent normalised concentration difference between a pair of neighbours, bars shows mean and SD; 2 independent experiments; 2-tailed Mann-Whitney test with p-values ***p = 0.003, ****p<0.00001.

We then used nuclear segmentation and pseudo-color analysis of mean Venus::HES5 intensity per nucleus (**Figure 7c**) and found that the interneuron domain shows the presence of microclusters mainly consisting of 2-3 cells wide in the dorsal to ventral axis whereas in the motoneuron domain high Venus::HES5 cells were mainly found as single cells, alternating with cells expressing lower Venus::HES5 (**Figure 7d**). We validated this finding further by investigating spatial periodicity by domain in live tissue slices. The domain border between motorneuron and interneuron progenitors was 35 μm ventral to the peak of HES5 expression (**Figure EV4d**) allowing us to correctly identify the two domains without the need for an OLIG2 reporter in the same tissue. We found that spatial periodicity was reduced in the motorneuron compared to the interneuron domain when analysed using both peak to peak distance in auto-correlation (**Figure 7e**, MN mean 31μm vs IN mean 41 μm and **Figure EV4e,f**) and dominant spatial period by Lomb-Scargle periodogram (**Figure EV4g**, MN mean 25 μm vs IN mean 40 μm). Thus, both nuclear segmentation analysis and spatial periodicity indicated that, in the interneuron domain, microclusters of 2-3 cells are found in a spatially periodic pattern repeated every 4 cells. Meanwhile, the motorneuron domain shows alternating high and low HES5 levels between neighbouring cells and a significant reduction in spatial periodicity, both of which are pointing to the motorneuron domain more closely resembling *Alternating high-low* conditions.

The model predicts that the coupling strength regulates the type of spatial micro-patterning hence, we hypothesised that the interneuron and motorneuron domains have different coupling strength. The model indicates that weak coupling, likely to be characteristic of the interneuron domain, would generate smaller cell-cell concentration differences compared to strong coupling (**Appendix Figure S7a**). This is because weakly coupled cells have less ability to repress the transcription of their neighbours and so are more similar in levels. This relationship should persist even after correcting for mean level in each condition. We have previously used fluorescence correlation spectroscopy (FCS) to generate a spatial map of nuclear Venus::HES5 concentration in the E10.5 spinal cord (Manning *et al.*, 2019b). Using this data, we calculated the difference in Venus::HES5 concentration between neighbouring cell-pairs relative to the mean by domain and indeed found that it is lower in the interneuron domain compared to the motorneuron domain (**Figure 7f**). The correction by mean was important as variability in expression is expected to scale with the mean. This finding was confirmed by measuring the spatial amplitude of Venus::HES5, which was also higher in the motorneuron domain (**Figure EV4h**). These findings are consistent with the notion that the coupling strength in the IN domain is lower than in MN one. Taken together, these results show that interneuron progenitors are more likely to be found in a locally synchronised state through weak coupling which correlates with a lower rate of differentiation. By comparison, progenitors in the motorneuron domain are mostly found in alternating high-low pattern and show a higher rate of differentiation, as predicted computationally by a higher coupling strength.

### NGN2 expression is spatially periodic and coordinates with the HES5 pattern

Given that the spatial pattern of HES5 is relevant to the rate of neurogenesis we investigated the wider applicability of our findings by characterising the spatial patterns of other genes in the Notch-Delta gene network. Chromogenic in-situ hybridisation of *Dll1* and *Jag1* mRNA shows that *Dll1* has a broad expression domain that covers the motor neuron domain and the ventral-most part of the interneuron domain (**Figure EV5a)** (Marklund *et al*, 2010). Alternate stripes of *Jag1* and *Dll1* are observed in the intermediate spinal cord, which covers the remaining part of the interneuron domain (**Figure EV5a)** (Marklund *et al.*, 2010). We have performed smiFISH for *Dll1* to get a high-resolution understanding of *Dll1* expression pattern in the interneuron domain where HES5 is expressed in microclusters. We found that *Dll1* expression is non-uniform and appeared in microstripes of a few cells (**Figure EV5b,c**), suggesting that other genes show similarities in local spatial patterning.

We next analysed the spatial expression pattern of the pro-neural factor NGN2. Using both NGN2 antibody staining and a NGN2:mScarlet fusion reporter mouse, we found that NGN2 also has a spatially periodic expression pattern, with around half the spatial period of Venus::HES5 (**Figure 8a-c)**. The spatial period of NGN2 is smaller in the motorneuron domain with a mean period of 21 μm supporting the conclusion that NGN2 spatial expression patterns are different between motorneuron and interneuron domains (**Figure 8d**). To understand how the NGN2 and Venus::HES5 periodic patterns map on to each other we used the cross-correlation function of the NGN2 and Venus::HES5 spatial profile from the same tissue (**Figure 8e,f**). The cross-correlation analysis showed the presence multiple peaks indicating coordination between the two signals that was not reflected in the brightfield control (**Figure 8f**). As expected for signals of different periodicities, we observed primary peaks indicative of positively correlated activity (**Figure 8f**, red arrowheads) as well as secondary peaks indicative of negatively correlated activity (**Figure 8g**, black arrowheads). The cross-correlation is also indicative of whether peaks of activity are present in the same area. To ascertain this we performed a phase-shift analysis by measuring phase shift as the absolute lag corresponding to the primary cross-correlation peak closest to lag 0. In **Figure 8f** a dominant peak falls close to lag 0 thus indicating that NGN2 and Venus::HES5 patterns coordinate in the same region. Thus we concluded that NGN2 shows a spatial periodic pattern of half the period of Venus::HES5 resulting in half of the NGN2 high cells occurring in HES5 high microclusters and half in HES5 low.

**Figure 8.**
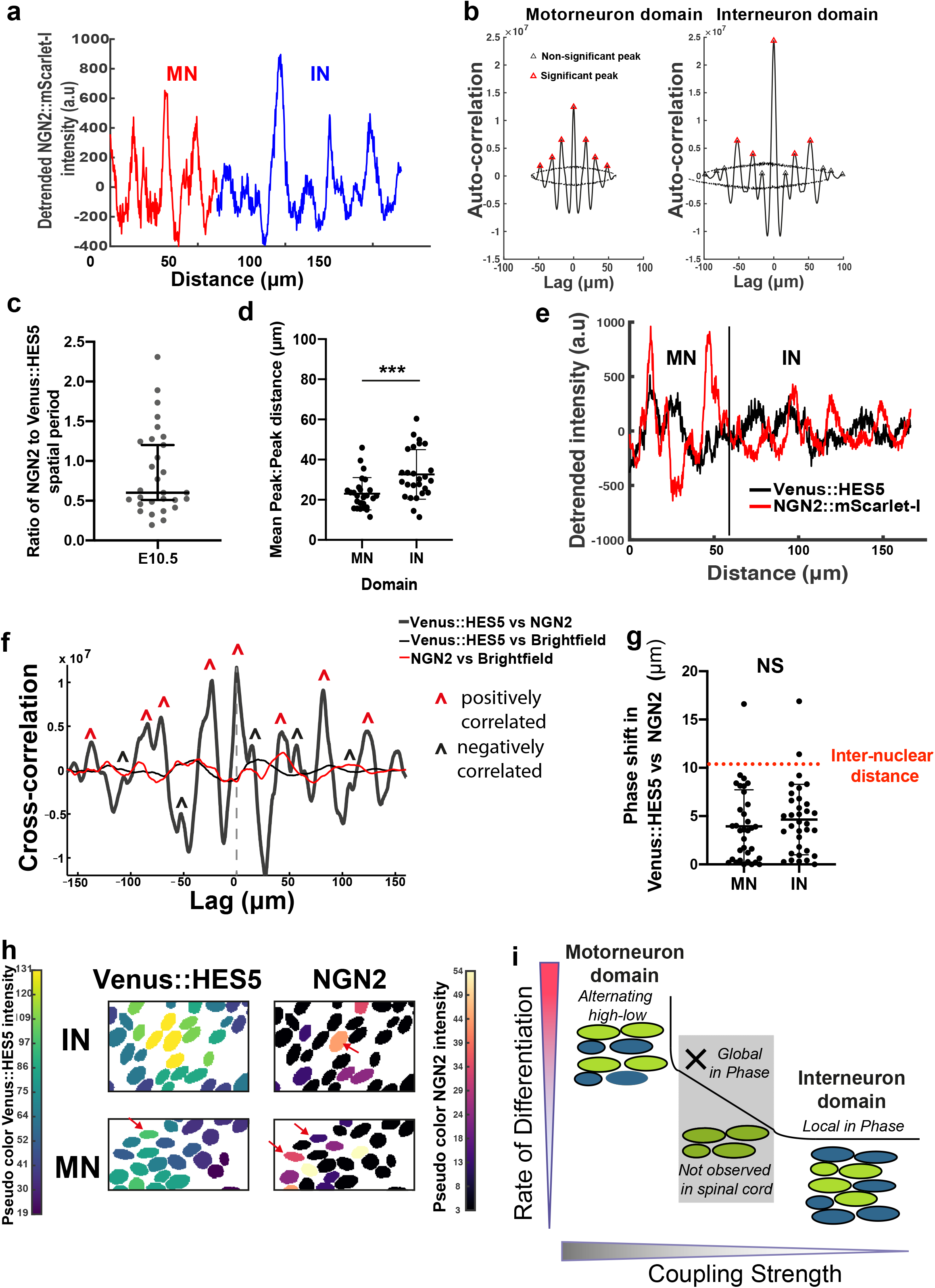
NGN2 expression is spatially periodic and positively correlates with the HES5 pattern. **(a)** Detrended spatial profile of NGN2::mScarlet-I intensity from transverse slice of E10.5 spinal cord from heterozygous knock-in mouse in ventral-dorsal direction;red indicates motorneuron (MN) domain, blue interneuron domain (IN). **(b)** Auto-correlation analysis of detrended NGN2::mScarlet-I intensity spatial profiles from motorneuron and interneuron domains; multiple peaks indicating spatial periodicity; significant peaks (red triangle) lie outside black dotted lines indicating 95% significance based on bootstrap approach (see Methods) and non-significant peaks (black triangle). **(c)** Peak to peak distance in auto-correlation plots of de-trended NGN2::mScarlet-I spatial profile in motorneuron (MN) and interneuron (IN) domains as a measure of spatial period in NGN2 expression along dorsal-ventral axis of spinal cord; Grey data points represent mean peak to peak distance in a single slice, n=33, left and right ventricle analysed separately in 4 experiments; black line shows overall mean, error bars show SD; 2-tailed Mann-Whitney test with exact p-value *** p=0.0003. (**d)** Ratio of NGN2:HES5 spatial period in the same tissue; grey dots show ratio for single image from 4 experiments; line shows overall median and error bars 95% confidence limits. (**e)** Detrended spatial profile of Venus::HES5 (black) and NGN2::mScarlet-I (red) intensity from the same transverse slice of E10.5 spinal cord in ventral-dorsal direction. (**f)** Example cross-correlation function of Venus::HES5 with NGN2::mScarlet-I (blue), Venus::HES5 with brightfield signal (black), and NGN2::mScarlet-I with brightfield signal (red) from the same transverse slice of E10.5 spinal cord; markers indicate the presence of two types of co-ordination namely in-phase (red arrowhead) and out-of-phase (black arrowhead). (**g)** Phase shift showing absolute lag distance corresponding to in-phase peak in Venus::HES5 vs NGN2::mScarlet-I cross-correlation function of spatial intensity profiles from the same slice; red line indicates average inter-nuclear distance in D-V; 2-tailed Mann-Whitney test not significant, p=0.32. **(h)** Pseudo-color look-up tables applied to mean nuclear Venus::HES5 and NGN2 staining intensity in motorneuron (MN) and interneuron (IN) domains. Venus::HES5 microcluster and single NGN2 high cell (red arrow) in IN domain; Alternating high-low expression of Venus::HES5 in MN, red arrows show high cells. **(i)** Graphical summary: Through a combination of experimental and computational work we characterised the HES5 dynamic expression in the mouse E10.5 ventral spinal cord. We found evidence that progenitors located in two domains (motorneuron, MN and interneuron, IN) give rise to distinct spatiotemporal characteristics that are indicative of differences in coupling strength and can explain increased differentiation rates observed in MN.

Furthermore when we performed phase-shift analysis in multiple cross-correlation examples (Methods), the shift was minimal and consistently less than a single-cell width (**Figure 8g)**. This strongly pointed to coordination not only in the same region but also in the same cells. We subsequently investigated this by using single nuclear segmentation of high resolution images to visualise the NGN2-HES5 spatial relationship. Indeed we found that within a HES5 microcluster in the interneuron domain, only 1-2 cells (and in the MN domain only one cell per cluster) show high NGN2 expression levels (**Figure 8h**). As high NGN2 is an early marker of differentiation, this suggests that similar to the mathematical model (**Appendix Figure S7b**) cells in a cluster do not differentiate in unison; instead microclusters may act to select a cell for differentiation, hence regulating spatial and temporal aspects of neurogenesis.

## Discussion

In this paper, we have addressed how cells co-ordinate their decisions with that of their neighbours so that neurogenesis takes place at a pace appropriate for the anatomical location. We have investigated the fine-grained pattern of neurogenesis in the spinal cord by monitoring the spatiotemporal patterning of key progenitor TF HES5 using live imaging analysis that is optimised towards revealing coordinated tissue level behaviour that would not otherwise be evident. In combination with computational modelling it enabled a multi-scale synthesis of the data with predictive power. We have uncovered an unexpected 3-tiered spatial and temporal organisation, which we discuss below in an ascending order of complexity.

First, within the ventral HES5 expression domain, which encompasses distinct MN and IN domains, we have discovered clusters of cells with positively corelated HES5 expression levels. These clusters, described for the first time here, are 2-3 cells wide in D-V and 3-4 cells wide in A-B axes, hence termed microclusters. To detect microclusters, we removed longer range spatial trends such as overall gradients of intensity in HES5 expression (which have not been dealt further here) allowing us to concentrate on local correlations of expression. By following Venus::HES5 in pairs of single cells in proximity we find that microclusters are a composite of positive correlations in levels of Venus::HES5 and locally synchronised (in-phase) HES5 dynamics. We also found that the microcluster organisation extends to DLL1 although we have not been able to study it with live imaging in this work. The clustering organisation was surprising because previous studies have suggested that in neurogenesis oscillators are in anti-phase in neighbouring cells (Kageyama *et al*, 2008; Shimojo *et al.*, 2016; Shimojo & Kageyama, 2016). DLL1 oscillations were observed with live imaging in tissue but only 2 neighbouring cells were monitored in close vicinity and a single example shown for anti-phase oscillations, which may explain the contradictions with our data. Thus, the discrepancy could be to a difference in scale of analysis or perhaps to the different molecules studied.

Second, we have found that HES5 microclusters are arrayed in a spatially periodic pattern along the D-V axis of the ventral HES5 domain, meaning that high and low HES5 expression clusters alternate regularly in space. We also found that NGN2 is expressed periodically along the D-V axis with half the periodicity of HES5 such that NGN2 high cells are found both within HES5 high and low microclusters. SmiFISH showed *Dll1* expressed in microstripes but the images of single Dll1 mRNA molecules were not amenable to auto-correlation thus, it is not known whether they occur on the same spatial scale as HES5. Multiple stripes of *Dll1* and *Jag1* and *Lfng* have been observed, but at the larger progenitor domain scale (Marklund *et al.*, 2010; Ramos *et al*, 2010). Such spatial periodicity at a fine level within the ventral HES5 domain contrasts with the large-scale organisation of HES5 in 2 separate broad domains along the D-V axis (Sagner *et al.*, 2018).

Thirdly, the HES5 spatial pattern of microclusters was not static but appeared dynamic over time; High HES5 expressing microclusters persisted for 6-8 hrs and then switched to low expression, while low expressing microclusters showed the opposite behaviour. In other words, high and low expressing microclusters alternated and sometimes created phase waves that travelled through the tissue over time. These waves are somewhat reminiscent of phase waves of LFNG and AXIN2 expression that are observed in somitogenesis (Baron & Galla, 2019; Sonnen *et al*, 2018; Tsiairis & Aulehla, 2016) however, in the spinal cord such phase waves were weak. This complex spatial and temporal dynamic pattern of HES5 in spinal cord generated two important questions: how might it be generated and what might it mean for neurogenesis? Knowing that Hes genes and HES5 in particular, are activated by Notch signalling, we treated ex-vivo spinal cord tissue with DBZ to disrupt Notch signalling. We observed that the Notch inhibitor treatment extinguished spatial periodicity gradually and slowly, over a period of 12hrs, concurrent with HES5 level downregulation. This is consistent with the amplitude death that we observed in single cell data under the same treatment (Manning *et al.*, 2019a). The effect of Notch inhibition was far more pronounced in the temporal nature of the pattern; in the absence of Notch signalling, the HES5 spatially periodic pattern of low and high expressing microclusters became “frozen” in time. These findings suggest that Notch signalling plays a part in making the pattern dynamic over time but cannot account for the entire spatiotemporal complexity of HES5 expression that we see ex vivo.

Computational modelling helped us to explore further the role of Notch in generating the spatiotemporal pattern of HES5 expression. We have used a simplified multiscale stochastic model of HES5 self-repression and inter-cellular coupling with delay, parameterised on our own experimental data, namely the single cell HES5 temporal period and extent of HES5 expression synchronicity between cells using the KOP. With this model we were able to explore the influence of the coupling strength between cells in producing spatiotemporal HES5 expression patterns. We found multiple spatiotemporal patterns, namely; an alternating high and low pattern (at high coupling strength), global tissue synchronisation (at mid coupling strength), and un-coordinated pattern (at no coupling), see **Appendix. Movie S2**. Importantly, at weak coupling strength and intercellular time delay that is consistent with previous reports, we observed the emergence of dynamic microclusters that matched our experimental observations. The emergence of dynamic patterns that do not resolve into steady HES “on” or “off” static patterns has been previously observed in a stochastic multicellular tissue model combining Notch-Delta and Hes auto-repression but not confirmed in tissue (Tiedemann *et al*, 2017). However, the dynamic microclusters in our model occurred infrequently (with a probability of 20-30%) even though the model takes into consideration stochasticity and time delays; two features that represent the tissue context well. The low frequency of clusters did not improve after detailed optimised exploration of parameter space, which led us to conclude that a Notch based cell-to-cell signalling with the assumptions we have made, recapitulates only part of the observed pattern in vivo. Extension of the model to include i) longer range cell-cell interaction via cytonemes, or due to the elongated shape of the progenitor cells, ii) increased complexity of the gene network such as cis-inhibition between Delta-Notch or differences in signalling between different Notch ligands, may be able to increase the fidelity of microcluster emergence. Indeed, it has been shown that such modifications increase the range of spatial patterns that can be obtained (Boareto *et al*, 2015; Cohen *et al.*, 2010; De Joussineau *et al*, 2003; Hadjivasiliou *et al*, 2019; Petrovic *et al*, 2014; Sprinzak *et al*, 2010). Other ways in which the model can be extended is to incorporate the influence of morphogen signalling gradients along the D-V axis or differentiation gradients along the A-B axis, as these are known to exist in the tissue.

Nevertheless, the computational model we developed, allowed us to explore the advantages that organisation in dynamic microclusters may offer as a developmental strategy for neurogenesis in the embryonic spinal cord. Overall, we found that the spatiotemporal HES5 pattern was affected by the coupling strength between cells and in turn, affected the rate of differentiation. Based on our findings we propose that a classic lateral inhibition alternating high-low HES5 pattern (achieved at high coupling strength) shows the highest rate of differentiation because it generates two HES5 states (“on” and “off”) in a spatially-alternating pattern and this is likely to result in tipping of more cells towards differentiation. Global synchronisation (medium coupling strength) shows a medium rate of differentiation; however this regime is not observed in spinal cord data perhaps because the synchronous differentiation in “blocks” of cells found close by in tissue, although an appropriate developmental strategy for somitogenesis, may be incompatible with the structural integrity of the neural tissue or the finer diversification of neuronal fates within each domain. The un-coordinated pattern (no coupling between cells) has similar rates of differentiation as weak coupling, however, weak coupling strength is advantageous because it allows local in phase synchronisation, which by analogy to global synchronisation (**Figure 6c**, *Global vs Local in phase)*, appears to transiently increase the amplitude of temporal oscillations in HES5 expression (**Figure 5i***, panel 3-Local in phase*). This is important because a transient amplitude increase (due to the presence of microclusters at *Local in phase* conditions) could faciliate the progression to differentiation. Indeed, we have previously shown that HES5 oscillations in proliferating spinal cord progenitors have low amplitude and show mainly aperiodic fluctuations (noisy dynamics) but the propensity to oscillate as well as the peak-to-trough amplitude increases as cells enter the differentiation pathway (Manning *et al.*, 2019a). We have also shown that when the transition from noisy dynamic expression to oscillatory expression does not take place, progenitor cells are unable to downregulate HES levels and differentiate (Soto *et al.*, 2020). We speculate that microclusters may act to reliably select one or two cells that go on to express NGN2 and differentiate and that the spatial periodicity of microclusters may space out differentiating cells to maintain tissue organisation.

We tested the model hypothesis that by changing the HES5 spatiotemporal pattern through tuning the coupling strength, the tissue is able to fine tune the rate of neurogenesis. We compared the motorneuron and interneuron progenitor domains as these two neighbouring domains in the D-V axis are known to have different rates of differentiation (Kicheva *et al.*, 2014; Kuzmicz-Kowalska & Kicheva, 2020). Indeed, we find that that in the MN domain where the rate of differentiation is highest at E10.5, the HES5 and NGN2 pattern most closely matches the alternating high-low pattern **(Figure 8i, MN)**. In the ventral interneuron domain, we propose that the local in phase synchronisation pattern (predicted to occur at weak coupling strength) is the closest match to the ex-vivo situation **(Figure 8i, IN)**. We propose it represents a strategy to balance prolonged neurogenesis, with a reasonable rate of differentiation and a transient increase in oscillation amplitude that is suitable for decoding by downstream genes. There may be additional molecular differences between the motorneuron and interneuron domains that regulate the rate of differentiation. Indeed, the transcription factor OLIG2 is expressed in the motorneuron domain and has been shown to promote differentiation by directly inhibiting HES5 (Sagner *et al.*, 2018). We speculate that this mechanism could interplay or directly affect the cell-cell coupling strength by changing HES5 levels or binding partners.

In conclusion, our findings show HES5 spatially periodic micro-patterns exist in the developing spinal cord, they underlie the rate of neurogenesis and are an emergent property of the multi-scale synthesis of dynamical gene expression and Notch coupling. This temporally dynamic expression is a testament to the power of live tissue imaging in providing mechanistic insights of complex phenomena as they unfold in real time.

## Materials and Methods

### Animals

Animal experiments were performed by personal license holders under UK Home Office project license PPL70/8858 and within the conditions of the Animal (Scientific Procedures) Act 1986. Venus::HES5 knock-in mice (ICR.Cg-Hes5<tm1(venus)Imayo>) were obtained from Riken Biological Resource Centre, Japan and maintained as a homozygous line. In these mice the mVenus fluorescent protein is fused to the N-terminus of endogenous HES5. Sox1Cre:ERT2 mice (Sox1tm3(cre/ERT2)Vep were obtained from James Briscoe with the permission of Robin Lovell-Badge. R26R-H2B::mCherry mice were obtained as frozen embryos from Riken Centre for Life Science Technologies, Japan and C57Bl6 mice were used as surrogates. NGN2::mScarlet-I mouse was generated by the University of Manchester Genome Editing Unit (see Appendix methods). The mScarlet-I fluorescent protein is fused to the C-terminus of endogenous NGN2.

### Embryo slicing and live imaging

E0.5 was considered as midday on the day a plug was detected. For matings with R26R-H2B::mCherry Sox1Cre:ERT2, intra-peritoneal injection of pregnant females with 2.5 mg Tamoxifen (Sigma) was performed 18 hours prior to embryo dissection. This enables single cell tracking through mosaic labelling of nuclei with H2B::mCherry. Whole embryos were screened for H2B::mCherry expression using Fluar 10x/0.5 objective on a Zeiss LSM880 confocal microscope. After decapitation, embryo bodies were embedded in 4% low-gelling temperature agarose (Sigma) containing 5mg/ml glucose (Sigma). 200μm transverse slices of the trunk containing the spinal cord around the forelimb region were obtained with the Leica VT1000S vibratome and released from the agarose. Embryo and slice manipulation were performed in phenol-red free L-15 media (ThermoFisher Scientific) on ice and the vibratome slicing was performed in chilled 1xPBS (ThermoFisher Scientific).

For snapshot imaging of live E10.5 spinal cord, slices were stained with 50μM Draq5 (Abcam – ab108410) in 1xPBS (ThermoFisher Scientific) for 1.5hrs on ice if required, and then placed directly on to a 35mm glass-bottomed dish (Greiner BioOne). Images were acquired with a Zeiss LSM880 microscope and C-Apochromat 40x 1.2 NA water objective. E10.5 spinal cord slices for live timelapse microscopy were placed on a 12mm Millicell cell culture insert (MerckMillipore) in a 35mm glass-bottomed dish (Greiner BioOne) incubated at 37°C and 5%CO_2_. The legs of the cell culture insert were sanded down to decrease the distance from the glass to the tissue. 1.5mls of DMEM F-12 (ThermoFisher Scientific) media containing 4.5mg/ml glucose, 1x MEM non-essential amino acids (ThermoFisher Scientific), 120ug/ml Bovine Album Fraction V (ThermoFisher Scientific), 55μM 2-mercaptoethanol, 1x GlutaMAX (ThermoFisher Scientific), 0.5x B27 and 0.5x N2 was added. Movies were acquired using Zeiss LSM880 microscope and GaAsP detectors. A Plan-Apochromat 20x 0.8 NA objective with a pinhole of 5AU was used. 10 z-sections with 7.5 μm interval were acquired every 15 mins for 18-24 hrs. DMSO (Sigma) or 2μM DBZ (Tocris) was added to media immediately before imaging.

### Single cell tracking over time

Single neural progenitor cells in E10.5 spinal cord slices were tracked in Imaris on the H2BmCherry channel using the ‘Spots’ function with background subtraction and the Brownian motion algorithm. All tracks were manually curated to ensure accurate single-cell tracking. Background fluorescence was measured via an ROI drawn on a non-Venus::HES5 expressing region on the tissue and subtracted from spot intensity. To account for any photobleaching and allow comparison of intensities between movies the mean intensity of mCherry and Venus in each spot was normalised to the mean intensity of mCherry or Venus in the whole tissue. The whole tissue volume was tracked using the ‘Surfaces’ and ‘Track over time’ function. For the calculation of Kuramoto order parameter from timeseries see **Temporal phase synchronisation**.

### Immunofluorescent staining

Trunks of E10.5 embryos for cryo-sectioning were fixed in 4% PFA for 1 hour at 4°C, followed by 3 quick washes with 1xPBS and 1 longer wash for 1 hour at 4°C. Embryos were equilibrated overnight in 30% sucrose (Sigma) at 4°C before mounting in Tissue-Tek OCT (Sakura) in cryomoulds and freezing at −80°C. 12μm sections were cut on Leica CM3050S cryostat. E10.5 spinal cord slices cultured on Millicell inserts were fixed in 4% PFA for 4 hours. For staining, tissue and sections were washed in PBS followed by permeabilisation in PBS 0.2% Triton X-100 (Sigma) and blocking with PBS 0.05% Tween20 (Sigma) + 5% BSA (Sigma). Primary and secondary antibodies were diluted in PBS 0.05% Tween20 + 5% BSA. Tissue was incubated with primary antibodies overnight at 4°C, then washed three times for 5– 10 minutes in PBS 0.05% Tween20, incubated with secondary antibodies and DAPI (Sigma) for 6 hours at room temperature, and washed again three times in PBS-T. Sections were mounted using mowiol 4-88 (Sigma). Primary antibodies used were rabbit anti-SOX2 (ab97959, 1:200), rabbit anti-OLIG2 (EMD Millipore AB9610, 1:200) and goat anti-NGN2 (Santa Cruz Biotechnology sc-19233, 1:200).

### smiFISH probe design and synthesis

The smFISH probes were designed using the probe design tool at http://www.biosearchtech.com/stellarisdesigner/. Depending on the GC content of the input sequence, the software can return varied size of probes, 18 and 22nt, hence giving the largest number of probes at the maximum masking level. It also uses genome information for the given organism to avoid probeswith potential off‐target binding sites. Using the respective gene mature mRNA sequence, we designed 36 probes for Hes5 and 48 probes for Dll1 (Tables X and Y) and added a FLAP sequence (5’-CCTCCTAAGTTTCGAGCTGGACTCAGTG-3’) to the 5’ of each gene-specific sequence (IDT). The designed set of probes were labelled with Quasar 670 (Biosearch Technologies) for Hes5 and CalFluor 610 (Biosearch Technologies) for Dll1 following the protocol from Marra et al. 2019

### smiFISH on mouse sections

smiFISH protocol for mouse section embryos was developed by adapting smiFISH protocol from Marra et al (2019) and Lyubimova et al (2013) (REF). 50μm‐thick sections of E10.5 spinal cord were collected and transferred onto superfrost glass slides (VWR 631‐0448) and kept at −80°C. Sections were left at room temperature to dry for 5-10 min and then fixed in 4% formaldehyde in 1× PBS followed by two quick washes in 1XPBS. 1:2000 dilution of proteinase K (20mg mL-1stock) in 1X PBS was pipetted onto each slide and left for 5-10 min followed by two washes in 2X SCC. Sections were then incubated at 37°C twice in wash buffer (5 ml of 20× SSC, 5 ml of formamide and 45 ml of deionized, nuclease-free water). 250uL of hybridization buffer (1 g dextran sulfate, 1 mL 20X SSC, 1 mL deionized formamide, 7.5 mL nuclease-free water) with 100-240nM the fluorescent smiFISH probes was pipetted onto each slide and incubated overnight at 37°C in a humid container shielded from light. Samples were then washed as follows: twice in wash buffer at 37°C for 3 min, twice in wash buffer at 37°C for 30 min, one wash in 1X PBS at room temperature for 5 min. After smiFISH staining sections were washed for 2min in PBS and mounted using Prolong Diamond Antifade Mountant with DAPI (Thermo Fisher P36962).

### smiFISH microscopy and deconvolution

smiFISH images were collected with Leica TCS SP8‐inverted confocal microscope using objective HC PL APO CS2 40x/1.30oil. We acquired three‐dimensional stacks 2048×1024pixels and z size 0.4μm. The voxel size was 0.19×0.19×0.4μm. Quasar 670 and CalFluor 610 were imaged with pinhole 1 Airy Unit. Channels were sequentially imaged. Deconvolution of confocal images was performed using Huygens Professional Software. As pre‐processing steps, the images were adjusted for the “microscopic parameters” and for additional restoration such as “object stabilizer”; the latter was used to adjust for any drift during imaging. Following, we used the deconvolution Wizard tool, the two main factors to adjust during deconvolution were the background values and the signal‐to‐noise ratio. Background was manually measured for every image and channel, while the optimal signal‐to‐noise ratio identified for the images was value 3. After deconvolution, the images were generated with Imaris 9.3

### Microcluster quantification and correlation of nuclear Venus::HES5 intensity with distance and neighbours

Individual Draq5+ nuclei were manually segmented as ellipses using ImageJ, converted to a mask and subsequently eroded using the ImageJ function ‘erode’ to ensure no overlap between nuclei. Small nuclei with very high Draq5 intensity were removed to avoid dead or dying cells. The mean intensity of each nuclear ROI was plotted using the ‘viridis’ (Venus::HES5) or ‘magma’ (NGN2) LUT. Microcluster size was calculated by quantile normalising the distribution of Venus::HES5 intensities across experiments, then counting the maximum number of adjacent cells in the highest colour band of the viridis LUT. The centroids of the manually segmented cells were used to measure distance and hence rank between neighbours and a correlation of the distance and mean nuclear Venus::HES5 intensity was calculated using the ‘corr’ function in MATLAB. Mean nuclear Venus::HES5 intensity was also randomised between nuclei and repeated 100 times before undergoing the same distance vs mean intensity correlation.

### Centre of intensity detection and radial gradient analysis

The centre of intensity (COI) was calculated using a centre of mass approach. The intensity of each nuclei was multiplied by their position. These were then summed and divided by the sum of all nuclear intensities. The COI was used to sort cells in to 5 equally spaced radial zones with increasing distance from the COI. The mean Venus::HES5 intensity of nuclei in these zones was calculated and then subtracted from each nucleus in that zone to remove the radial gradient. A simulated radial gradient from a single focal point in the image was generated using

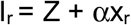

Where **I**_**r**_ is the new intensity of the cells, Z is simulated intensities with the mean and variance similar to that of real data, α is the gradient strength parameter and x_r_ is a function of the distance from the centre of intensity. As α increases the radial gradient is less affected by random deviation in HES5 expression.

### Detection and periodicity analysis of spatial expression patterns

To remove the possibility that changes in cell positions lead to shifts in the kymograph stripes and artifacts in the dynamic analysis, movies underwent image registration to account for global tissue drift and were subject to strict quality controls for local tissue deformation. The average motility of tracked single cells over time in the D-V axis was compared to patterns/waves of Venus::HES5 intensity in the kymograph and included a maximum threshold of the averaged single cell displacement (see **Appendix Table S1**). Bleach correction was then performed using a ratiometric method in ImageJ.

Spatial expression profiles and kymographs were generated in Zen Blue (Carl Zeiss Microscopy) by drawing a spline 15 μm wide starting ventrally and extending parallel to the ventricle in the dorsal direction, then using the ‘Line profile’ or ‘Kymograph’ function. A 15 μm width was chosen as this covered more than a cell width and the average distance a cell travels in 2 hrs, the value used for time averaging. 0 distance corresponded to the ventral most end of the spline. Apical, medium and basal expression profiles and kymographs were generated from splines around 10, 30 and 60 μm from the ventricle respectively, and analysing each side of the ventricle separately. 2-3 non-overlapping z-sections were used to generate kymographs per movie. Expression profile data for Draq5 and NGN2 from single snapshot images of live slices were generated in ImageJ using a rectangular ROI of width 15μm and the “Plot profile” function.

Kymographs were analysed using custom scripts in MATLAB. Kymographs were averaged along the time axis in 2 hr windows. Subsequently spatial Venus::HES5 intensity in the ventral-dorsal direction was de-trended by fitting a polynomial (order 4-6) and subtracting this from the raw data. This removed the larger trend due to the profile of the HES5 expression domain.

Auto-correlation and Lomb-Scargle periodograms were used to analyse periodicity of the de-trended spatial intensity plots. Lomb-Scargle periodograms were generated with the MATLAB ‘plomb’ function and plotting power level thresholds as specified in figure legends. Auto-correlation was performed with the MATLAB ‘xcorr’ function. Auto-correlation functions were smoothed using Savitzky-Golay filter and then peaks identified using the ‘find peaks’ function. Significant peaks were identified using a bootstrap method with 100 randomisations. Auto-correlations were randomised and then re-subjected to auto-correlation. 2 standard deviations of the auto-correlations of randomised data was used as a threshold and peaks were designated as significant if they exceeded this threshold. The mean distance between significant peaks was calculated per kymograph timepoint. Fold-changes of spatial intensities were calculated between significant peaks and troughs in the signal identified using ‘find peaks’ on the negative signal.

Splitting Venus::HES5 kymographs in to motorneuron and interneuron domains was based on staining of cryo-sectioned E10.5 spinal cord with motorneuron progenitor domain marker OLIG2. The peak of the trend in Venus::HES5 was found to occur on average at 35 μm dorsally from the edge of the OLIG2+ domain. This criteria was used to split kymographs from movies of Venus::HES5 spinal cords that had not been immuno-stained.

### Correlation coefficient analysis in the anterior to posterior (A-P) axis

We produced kymographs from multiple non-overlapping stacks extending in the AP direction using the same region of interest (ROI) which meant that Venus:HES5 intensity was comparable at the same position in DV. We used detrended Venus::HES5 averaged over 2h per z and compared the detrended coefficients pairwise across subsequent z-stacks. Using the confocal magnification in the AP axis per experiment we reconstructed the absolute distance between subsequent z-stacks. Data from untreated and tissue treated with DMSO was analysed in the same way.

### Hierarchical clustering of local HES5 expression and microcluster persistence time

Kymographs of HES5 expression were split into adjacent 20μm regions along the D-V axis and the HES5 intensity averaged in these regions to give a timeseries per region. To account for any single cell movement in DV, we applied a 2um Gaussian blur filter onto the kymograph data using the Matlab routine *imgaussfilt.m* prior to extracting timeseries per region.These timeseries were normalized to the mean and standard deviation of each region over time (z-scoring) and subject to hierarchical clustering using the *clustergram,m* routine in Matlab with Euclidean distance and average linkage. The persistence time was calculated as continuous time when the signal in the region was above (high) or below (low) its mean level. The persistence ratio was calculated as the time interval spent in a high state divided by the time interval spent in a low state within the same 20μm region. Where only high or low persistence time intervals were detected in a region these observations were excluded from the ratio. We also used an alternative method to compute persistence time relying on zero-crossing of the detrended Venus::HES5 signal averaged over 0 to 2h timepoints; in this approach we identified specific areas containing a microcluster with high expression (above the mean) and low expression (below the mean) and repeated the persistence time calculation as described above. For the calculation of phase synchronisation index see **Spatial phase synchronisation**.

### Phase Mapping and Phase Shift Analysis in Cell Pairs

We analysed Venus::HES5 ultradian dynamics using the approach in Manning et *al.* 2019, Phillips et *al.* 2017. Specifically, we used a Gaussian Processes pipeline to fit the single cell trend of Venus::HES5 expression (examples shown in **Appendix Figure S2d**). We performed detrending of Venus::HES5, followed by z-scoring and estimated a periodic Ornstein-Uhlenbeck covariance model. This procedure produces a smooth detrended curve (examples shown in **Appendix Figure S4b**). Using the detrended smoothed curves, we extracted the phase shift using cross-correlation analysis of pairs of timeseries. The pairing was done using Euclidean distance in the same tissue. The phase shift corresponded to the lag time interval closest to 0 at which the cross-correlation function shows a peak. From detrended smooth curves, we then performed Hilbert reconstruction of instantaneous phase using the *hilbert.m* Matlab routine. We used the phase angles corresponding to neighbouring cell pairs at multiple timepoints to produce a phase mapping. We plotted the density of the phase map using the *dscatter.m* routine with 24×24 binning of phase values (Eilers and Goeman, Bioinformatics 2004).

### Stochastic Multicellular HES5 Model with Time Delay

The core unit of the multicellular model is a single-cell unit that explicitly models Hes5 protein and mRNA abundance and is adapted from the work done in Manning et *al.*, 2019. The single cell model makes use of a Langevin approach to include stochastic fluctuations in both protein and mRNA as well as the inclusion of a time delay associated with the inhibitory Hill function used to describe the repressive action of Hes protein on its own mRNA production. This implementation, along with the parameter inference (Manning et al., 2019) results in a single cell model capable of reproducing stochastic oscillations closely matched with the single-cell dynamics observed in the developing neural tube. The multicellular approach extends the single cell model by introducing an inhibitory Hill function to couple nearest-neighbour cells (in a fixed, no cell movement, hexagonal geometry) whereby high Hes5 protein in one cell is able to repress Hes5 mRNA production in a neighbouring cell. This inhibitory Hill function (the coupling function) is representative of the overall behaviour of the Notch Delta pathway and its interaction with Hes5, allowing for the bidirectional interaction of Hes5 dynamics between neighbouring cells. Three parameters are associated with this Hill function that make it flexible enough to explore different possible coupling realisations of the Notch-Delta pathway, the effects of which are illustrated in **Figure 5b**. The main parameter modulated for the analysis in this paper is the repression threshold which defines the number of protein molecules that is required to repress mRNA in a neighbouring cell.

### Cell to cell HES5 differences by domain and by coupling strength

We used raw Venus::HES5 data, absolute HES5 quantitation by Fluorescence Correlation Spectroscopy (FCS) and manually segmented nuclear maps made available in (Manning et al., 2019). We obtained average HES5 concentration per nuclei by quantile-quantile matching the Venus distribution to the reference FCS distribution of HES5 levels across the tissue. Using nuclear centroid location, we produced absolute cell to cell concentration differences between every cell and its closest neighbor. We performed a by domain analysis by dividing the cell to cell concentration differences by the average HES5 concentration by domain. In the synthetic examples, HES5 molecular abundance data obtained from the multicellular model was used to produce absolute cell to cell abundance differences over a range of coupling strength values. We also produced synthetic cell to cell abundance differences relative to the mean HES5 abundance per simulation over a range of coupling strength values.

### Phase reconstruction and Kuramoto order value as a measure of synchrony

To determine the synchronisation of real signals both in the model and experimental data, the phase of each oscillator was first reconstructed in complex space. This reconstruction was achieved by using the Hilbert transform, which shifts the phase of each frequency component in a signal by 90 degrees (Benedetto, 1996). The Hilbert transform of a function *u*(*t*) is defined as

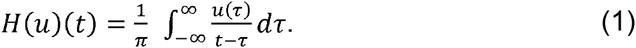

To obtain a rotating vector that contains both the amplitude and phase information of the signal at a given time *t*, the original signal and the 90 degrees shifted Hilbert transform can be combined in complex space to give

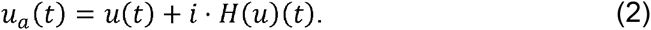

By comparing *u*_*a*_(*t*) of two or more cells, a measure of how synchronised a population of cells is can be determined by first calculating what is known as the complex order parameter

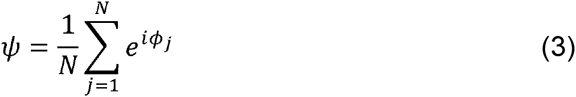

where N is the number of oscillators and *ϕ*_*j*_is the phase of oscillator *j*. From this, the Kuramoto order parameter is defined as the absolute value of the complex order parameter *ψ*, which is the magnitude of the vector and has a value between 0 and 1 (Choi *et al.*, 2000). A value of 1 indicates perfect synchrony and matching phase, meaning that in complex space the phases of each oscillator would be at the same angle and would rotate at the same frequency. A value of 0 indicates no synchronisation, and in complex space would appear as a distribution of phases that average to a point at the origin.

### Phase synchronisation index

In addition to calculating KOPs, we also used the Hilbert transform to extract phase from spatial data to determine how dynamic the positions of peak and trough were over time. This involved extracting and plotting the phase from time-averaged spatial signals. The phase synchronisation index for DMSO and DBZ conditions (**Figure 4d)** was obtained by calculating KOP per position in D-V axis and averaging per z-slice (with left and right of the ventricle analysed separately).

### Statistical testing

Statistical tests were performed in GraphPad Prism 8. Data was tested for normality with D’Agostino-Pearson test. The relevant parametric or non-parametric test was then performed. Bar plots and discrete scatter plots show mean mean±SD where multiple independent experiments are analysed. Statistical significance between 2 datasets was tested with either t-test (parametric) or Mann-Whitney test (non-parametric). Statistical significance (p<0.05) for 2+ datasets was tested by Kruskall-Wallis with Dunn’s multiple comparison correction. All tests were 2-sided. Multiple comparison testing involved comparing all pairs of data columns. Correlations were analysed using Pearson correlation coefficient. Sample sizes, experiment numbers, p values<0.05 and correlation coefficients are reported in each figure legend.

## Acknowledgements

We are grateful to members of the Papalopulu lab, Andrew Hazel, James Briscoe and Andy Oates for advice and discussions. The authors would also like to thank Robert Lea, the Biological Services Facility and the Bioimaging Facilities of the University of Manchester for technical support. CM was supported by a Sir Henry Wellcome Fellowship (103986/Z/14/Z) and University of Manchester Presidential Fellowship. VB was supported by a Wellcome Trust Senior Research Fellowship to NP (090868/Z/09/Z). JH, EJ and DH (Wellcome Trust Grant No. 215189/Z/19/Z) were supported by Wellcome Trust PhD studentships. JK was supported by Wellcome Trust Senior Research Fellowship to NP and a University of St Andrews Lectureship. The funders had no role in study design, data collection and analysis, decision to publish, or preparation of the manuscript.

## Author Contributions

CM and NP conceived and designed the experimental study.

CM performed wet lab experiments, supervised and performed data analysis, interpreted data and wrote the paper.

VB supervised and performed development of method to analyse spatial micropatterns of HES5 expression both in data and from the model, developed method for Hilbert phase persistence analysis, performed data analysis, interpreted data and wrote the paper.

JH designed and implemented stochastic coupled HES5 model, explored parameterization, analysed and interpreted the model.

XS performed smiFISH and imaging.

EJ collected data and developed method to analyse correlations of HES5 nuclear intensity.

DH contributed to the method to analyse periodic spatial micropatterns of HES5 expression.

HB designed and generated Neurog2::mScarlet-I knock-in mouse ADA designed and generated Neurog2::mScarlet-I knock-in mouse

JK supervised and assisted design, analysis and interpretation of the mathematical model.

PG supervised and assisted analysis and interpretation of the mathematical model. NP supervised and directed the work, interpreted data and co-wrote the paper with CM, VB and JH with input from JK, EJ, DH and PG.

## Conflict of interest

None.

## Code and Data availability

All code written in MatLab and is available on GitHub: https://github.com/Papalopulu-Lab/Biga2020

## Expanded View Figure Legends

**Figure EV1. Venus::HES5 expression in single progenitor cells located in the ventral domain of spinal cord. Related to Figure 1. (a)** Transverse slice of live Venus::HES5 homozygous knock-in mouse spinal cord E10.5 ex vivo, Draq5 live nuclear stain and brightfield image. Scale bar 50μm. (**b)** Immunofluorescence of E10.5 Venus::HES5 transverse slice of spinal cord ex-vivo. SOX2+ progenitors and endogenous Venus::HES5 signal. Scale bar 40 μm. **(c)** Viridis look-up table applied to mean nuclear Draq5 intensity corresponding to transverse slice in **Figure 1c**. **(d)** Pearson correlation coefficient of mean nuclear Venus::HES5 intensity in relationship to distance for **(i)** E9.5 and **(ii)** E11.5 Venus::HES5 spinal cord ex-vivo slices; red dots indicate average Venus::HES5 correlation per slice of 3 slices from 3 experiments with corresponding red line indicating one phase decay fit. Black line denotes 95% confidence levels. Gray dots indicate correlations from randomizations of intensities analysed in the same way. (**e**) Transverse slice of live **(i)** Venus::HES5 homozygous knock-in mouse spinal cord E10.5 ex vivo showing **(ii)** segmentation of Draq5 and **(iii)** mask applied to Venus::HES5 channel. Images correspond to slice shown in **Figure 1g**. **(f)** Representative example of mean Venus::HES5 intensity as a function of radial distance with respect to center of intensity (see Methods). **(g)** and **(h)** Left panels - Viridis look-up table applied to mean nuclear Venus::HES5 intensity in E9.5 and E11.5 slices respectively after radial gradient removal. Right panels - Pearson correlation coefficient of mean nuclear Venus::HES5 intensity in relationship to distance for E9.5 and E11.5 Venus::HES5 spinal cord ex-vivo slices respectively. Red dots indicate average Venus::HES5 correlation per slice of 3 slices from 3 experiments with corresponding red line indicating one phase decay fit. Black line denotes 95% confidence levels. Gray dots indicate correlations from randomizations of intensities analysed in the same way.

**Figure EV2. Draq5 and Venus::HES5 spatial periodicity in live spinal cord tissue slices. Related to Figure 2. (a)** Transverse slice of live Venus::HES5 homozygous knock-in mouse spinal cord E10.5 ex vivo, Draq5 live nuclear stain; rectangle shows region for spatial profile. **(b)** Viridis look-up table applied to mean nuclear Venus::HES5 intensity in the same slice shown in **(a)** and **(b)**, rectangle shows the same region of interest for spatial profile. **(c)** Venus::HES5 intensity spatial profile (black) from the yellow box in **(a)**, and trend line in blue (fitted polynomial order 6). **(d)** Detrended spatial profile of Venus::HES5 (grey) and Draq5 (red) nuclear stain from region delineated in **(a)**; arrows show regions of Venus::HES5 low and Draq5 high indicating low Venus::HES5 areas are not nuclei free. **(e)** Distribution of peak-to-peak distance in auto-correlation plots of Draq5 spatial profile; this is a measure of inter-nuclear distance in Draq5 profile along dorsal-ventral axis of spinal cord; data points represent all peak to peak distances from 9 slices, 6 experiments; mean 9.3 ± 0.42 μm (95% confidence limits). **(f)** Spatial periodicity detected by Lomb-Scargle periodogram in apical, medium and basal regions (10, 30 and 60 μm from ventricle respectively); dots indicate mean periodicity from at least 3 z-sections and both left and right sides of ventricle analysed in 6 experiments; lines indicate mean and SD per experiment; Kruskal-Wallis test not significant, p=0.3137. **(g)** Longitudinal cryo-section of neural tube in E10.5 Venus::HES5 embryos. A-anterior, P-posterior. Scale bar 60 μm. **(h)** Representative detrended spatial profile of Venus::HES5 from neural tube in anterior-posterior (A-P) direction. **(i)** Representative auto-correlation of Venus::HES5 spatial profile in A-P direction. Multiple significant peaks indicate spatial periodicity in A-P direction. Significant peaks (red triangle) lie outside black dotted lines indicating 95% significance based on bootstrap approach (see Methods) and non-significant peaks (black triangles). **(j)** Pearson correlation coefficient of detrended Venus::HES5 spatial profile between subsequent z-sections of transverse E10.5 spinal cord tissue slices at known distances (i.e correlations in A-P direction); untreated slices – dots show 18 pairs of z-sections from 6 experiment; DMSO treated slices – dots show 9 pairs from 3 experiments. **(k)** (Left panel) Representative spatiotemporal plot of the detrended Venus::HES5 pattern along ventral-dorsal direction (0 to 200μm) obtained by averaging kymograph data in the same region over 2 hr time intervals; (Right panel) Representative phase map of spatially periodic Venus::HES5 intensity obtained using the Hilbert transform (see Methods) from data shown in the left panel; markers indicate areas underdoing a high-to-low * and low-to-high** transition.

**Figure EV3. Changes in Venus::HES5 spatiotemporal expression pattern in live slice cultures treated with Notch inhibitor DBZ. Related to Figure 4. (a)** Representative examples of the detrended Venus::HES5 signal observed along ventral-dorsal direction in DMSO and DBZ conditions; Venus::HES5 intensity data obtained by averaging kymographs over 2 hrs intervals; panels represent individual slice cultures (2 per condition) in addition to examples in **Figure 4b**. (**b)** Spatial phase maps obtained from detrended Venus::HES5 signal in DMSO and DBZ conditions; panels correspond to Venus::HES5 intensity traces shown in **(a). (c)** Auto-correlation analysis of detrended apical Venus::HES5 spatial profile in DMSO (control) and DBZ (2 μM) treated E10.5 Venus::HES5 spinal cord slices; panels show auto-correlation of detrended Venus::HES5 signal averaged from 0-2h of timelapse video (top panels) and corresponding auto-correlation functions **i** n the same slices averaged in 2hr windows for 10h; we observed a decrease in amplitude of auto-correlation peaks over time in DBZ-treated slices. **(d)** Peak to peak distance in auto-correlation from spatial data shown in **Figure 4g** presented as median per experiment from DMSO (n=3 experiments) and DBZ (n=4 experiments); 1 tailed t test p=0.0526. **(c)** Nuclear density represented by the % area covered by nuclei in DMSO and DBZ treated ex-vivo E10.5 spinal cord slice cultures dots indicate multiple z-stacks from DBZ (5 slices) and DMSO (4 slices); bars indicate mean and SD per condition; 2 tailed t test p=0.0004***.

**Figure EV4. HES5 spatiotemporal dynamics correlate with rate of differentiation. (a)** Diagram of Notch-Delta inter-cellular communication and HES5 interactions. Our model takes into account that HES5 negatively regulates its own mRNA production (gray highlighted area), downstream proneural genes and Delta; the direct and/or indirect (via proneural genes) repression of Delta by HES5 is expressed mathematically through the means of an inter-cellular Hill function (see Methods). **(b)** Transverse cryosection of E10.5 Venus::HES5 spinal cord. Venus::HES5 endogenous signal, OLIG2: motorneuron progenitor marker, NGN2: early marker of neuronal commitment, DAPI; scale bar 25 μm. **(c)** Spatial expression profile of Venus::HES5, NGN2 and OLIG2 from the same tissue to help delineate motorneuron (OLIG2+) vs interneuron (OLIG2-) domains. **(d)** Spatial profile of Venus::HES5 intensity (black) generated by averaging 2.5hrs of kymograph data; 0 distance represents ventral end of kymograph; blue dotted line is trend in Venus::HES5 data across the domain determined by polynomial fit order 6; domain boundary between motorneuron progenitors (MN) and interneuron progenitors (IN) marked with red dashed line. **(e)** Detrended spatial profile of Venus::HES5 corresponding to **(c)** in motorneuron progenitors (red:MN) and interneuron progenitors (blue:IN). **(f)** Auto-correlation plot of detrended Venus::HES5 spatial profile in MN and IN progenitors; black lines show confidence limits for peak significance based on bootstrap approach on de-trended Venus::HES5 intensity profile (see Methods); red triangle - significant peak, black triangle – non-significant peak; multiple significant peaks in auto-correlation shows periodicity in spatial profile of Venus::HES5 intensity. **(g)** Spatial periodicity in motorneuron vs interneuron domain measured with the Lomb-Scargle periodogram; top 2 significant peaks were used to calculate spatial period from 2 to 3 z-sections per experiment, left and right side of ventricle analysed separately and 6 experiments; bars indicate mean with SD; Mann-Whitney test with two-tail significance for p<0.0001****. **(h)** Fold-change in Venus::HES5 spatial pattern between hi-low regions in IN domain relative to MN domain. Data points represent mean per experiment. Lines shows mean and SD of 6 experiments. 2-tailed Mann-Whitney test p=0.0022**.

**Figure EV5. Exploration of HES5 spatiotemporal dynamics in a multicellular environment. (a)** Schematic of *Hes5*, *Dll1* and J*ag1* expression in progenitor domains along D-V axis of E10.5 mouse spinal cord compiled based on data from (Manning *et al.*, 2019b) and (Marklund *et al.*, 2010). **(b)** DAPI nuclear stain of interneuron region in E10.5 mouse; lines indicate spatial localisation in tissue. **(c)** Single-molecule FISH of *Dll1* and *Hes5* expression in spinal cord region corresponding to **(b)** (Materials and Methods); panels 1 and 2 indicate *Dll1* and *Hes5* respectively; panels 3 and 4 show false-color merged *Dll1/Hes5* and *Dll1/Hes5/DAPI* respectively; scale bar 20 μm.

## Notes

### Competing Interest Statement

The authors have declared no competing interest.

### Summary of Updates

New figures and experimental data and analysis added.

## References

Bansod S, Kageyama R, Ohtsuka T (2017) Hes5 regulates the transition timing of neurogenesis and gliogenesis in mammalian neocortical development. Development (Cambridge) 144: 3156–3167

Baron JW, Galla T (2019) Intrinsic noise, Delta-Notch signalling and delayed reactions promote sustained, coherent, synchronized oscillations in the presomitic mesoderm. J R Soc Interface 16: 20190436

Benedetto JJ (1996) Harmonic analysis and applications. CRC Press

Boareto M, Jolly MK, Lu M, Onuchic JN, Clementi C, Ben-Jacob E (2015) Jagged-delta asymmetry in Notch signaling can give rise to a sender/receiver hybrid phenotype. Proceedings of the National Academy of Sciences of the United States of America 112: E402–E409

Bonev B, Stanley P, Papalopulu N (2012) MicroRNA-9 Modulates Hes1 ultradian oscillations by forming a double-negative feedback loop. Cell Rep 2: 10–18

Briscoe J, Small S, 2015. Morphogen rules: Design principles of gradient-mediated embryo patterning. Company of Biologists Ltd, pp. 3996–4009.

Choi MY, Kim HJ, Kim D, Hong H (2000) Synchronization in a system of globally coupled oscillators with time delay. Phys Rev E Stat Phys Plasmas Fluids Relat Interdiscip Topics 61: 371–381

Cohen M, Georgiou M, Stevenson NL, Miodownik M, Baum B (2010) Dynamic Filopodia Transmit Intermittent Delta-Notch Signaling to Drive Pattern Refinement during Lateral Inhibition. Developmental Cell

Corson F, Couturier L, Rouault H, Mazouni K, Schweisguth F (2017) Self-organized Notch dynamics generate stereotyped sensory organ patterns in Drosophila. Science (New York, NY) 356: eaai7407–eaai7407

Das RM, Storey KG (2012) Mitotic spindle orientation can direct cell fate and bias Notch activity in chick neural tube. EMBO Rep 13: 448–454

Das RM, Storey KG (2014) Apical abscission alters cell polarity and dismantles the primary cilium during neurogenesis. Science 343: 200–204

De Joussineau C, Soulé J, Martin M, Anguille C, Montcourrier P, Alexandre D (2003) Delta-promoted filopodia mediate long-range lateral inhibition in Drosophila. Nature 426: 555–559

de Lichtenberg KH, Funa N, Nakic N, Ferrer J, Zhu Z, Huangfu D, Serup P (2018) Genome-Wide Identification of HES1 Target Genes Uncover Novel Roles for HES1 in Pancreatic Development. bioRxiv: 335869

Delile J, Rayon T, Melchionda M, Edwards A, Briscoe J, Sagner A (2019) Single cell transcriptomics reveals spatial and temporal dynamics of gene expression in the developing mouse spinal cord. Development: dev.173807–dev.173807

Galla T (2009) Intrinsic fluctuations in stochastic delay systems: Theoretical description and application to a simple model of gene regulation. Physical Review E 80: 021909–021909

Goodfellow M, Phillips NE, Manning C, Galla T, Papalopulu N (2014) microRNA input into a neural ultradian oscillator controls emergence and timing of alternative cell states. Nat Commun 5: 3399

Hadjivasiliou Z, Moore RE, McIntosh R, Galea GL, Clarke JDW, Alexandre P (2019) Basal Protrusions Mediate Spatiotemporal Patterns of Spinal Neuron Differentiation. Developmental Cell 49: 907–919.e910

Henrique D, Schweisguth F (2019) Mechanisms of Notch signaling: a simple logic deployed in time and space. Development 146

Herrgen L, Ares S, Morelli LG, Schröter C, Jülicher F, Oates AC (2010) Intercellular coupling regulates the period of the segmentation clock. Current Biology 20: 1244–1253

Hunter GL, Hadjivasiliou Z, Bonin H, He L, Perrimon N, Charras G, Baum B (2016) Coordinated control of Notch/Delta signalling and cell cycle progression drives lateral inhibition-mediated tissue patterning. Development (Cambridge, England) 143

Imayoshi I, Isomura A, Harima Y, Kawaguchi K, Kori H, Miyachi H, Fujiwara T, Ishidate F, Kageyama R (2013) Oscillatory control of factors determining multipotency and fate in mouse neural progenitors. Science 342: 1203–1208

Imayoshi I, Kageyama R (2014) bHLH factors in self-renewal, multipotency, and fate choice of neural progenitor cells. Neuron 82: 9–23

Isomura A, Ogushi F, Kori H, Kageyama R (2017) Optogenetic perturbation and bioluminescence imaging to analyze cell-to-cell transfer of oscillatory information. Genes and Development 31: 524–535

Jensen MH, Sneppen K, Tiana G (2003) Sustained oscillations and time delays in gene expression of protein Hes1. FEBS Letters 541: 176–177

Kageyama R, Ohtsuka T, Shimojo H, Imayoshi I (2008) Dynamic Notch signaling in neural progenitor cells and a revised view of lateral inhibition. Nature Neuroscience 11: 1247–1251

Kicheva A, Bollenbach T, Ribeiro A, Valle HP, Lovell-Badge R, Episkopou V, Briscoe J (2014) Coordination of progenitor specification and growth in mouse and chick spinal cord. Science (New York, NY) 345: 1254927–1254927

Kobayashi T, Mizuno H, Imayoshi I, Furusawa C, Shirahige K, Kageyama R (2009) The cyclic gene Hes1 contributes to diverse differentiation responses of embryonic stem cells. Genes Dev 23: 1870–1875

Kuzmicz-Kowalska K, Kicheva A (2020) Regulation of size and scale in vertebrate spinal cord development. Wiley Interdiscip Rev Dev Biol: e383

Lewis J (2003) Autoinhibition with transcriptional delay. Current Biology 13: 1398–1408

Ma Q, Chen Z, del Barco Barrantes I, de la Pompa JL, Anderson DJ (1998) neurogenin1 is essential for the determination of neuronal precursors for proximal cranial sensory ganglia. Neuron 20: 469–482

Manning CS, Biga V, Boyd J, Kursawe J, Ymisson B, Spiller DG, Sanderson CM, Galla T, Rattray M, Papalopulu N (2019a) Quantitative single-cell live imaging links HES5 dynamics with cell-state and fate in murine neurogenesis. Nature communications

Manning CS, Biga V, Boyd J, Kursawe J, Ymisson B, Spiller DG, Sanderson CM, Galla T, Rattray M, Papalopulu N (2019b) Quantitative single-cell live imaging links HES5 dynamics with cell-state and fate in murine neurogenesis. Nat Commun 10: 2835

Marklund U, Hansson EM, Sundström E, de Angelis MH, Przemeck GK, Lendahl U, Muhr J, Ericson J (2010) Domain-specific control of neurogenesis achieved through patterned regulation of Notch ligand expression. Development 137: 437–445

Momiji H, Monk NAM (2008) Dissecting the dynamics of the Hes1 genetic oscillator. Journal of Theoretical Biology 254: 784–798

Monk NAM (2003) Oscillatory Expression of Hes1, p53, and NF-κB driven by transcriptional time delays. Current Biology 13: 1409–1413

Morelli LG, Ares S, Herrgen L, Schröter C, Jülicher F, Oates AC (2009) Delayed coupling theory of vertebrate segmentation. HFSP Journal 3: 55–66

Nelson BR, Hodge RD, Daza RA, Tripathi PP, Arnold SJ, Millen KJ, Hevner RF (2020) Intermediate progenitors support migration of neural stem cells into dentate gyrus outer neurogenic niches. Elife 9

Oates AC (2020) Waiting on the Fringe: cell autonomy and signaling delays in segmentation clocks. Curr Opin Genet Dev 63: 61–70

Ohtsuka T, Ishibashi M, Gradwohl G, Nakanishi S, Guillemot F, Kageyama R (1999) Hes1 and Hes5 as notch effectors in mammalian neuronal differentiation. The EMBO journal 18: 2196–2207

Özbudak EM, Lewis J (2008) Notch signalling synchronizes the zebrafish segmentation clock but is not needed to create somite boundaries. PLoS Genetics 4

Paridaen JT, Huttner WB (2014) Neurogenesis during development of the vertebrate central nervous system. EMBO Rep 15: 351–364

Petrovic J, Formosa-Jordan P, Luna-Escalante JC, Abello G, Iban M, Es, Neves J, Giraldez F, Abelló G, Ibañes M et al (2014) Ligand-dependent Notch signaling strength orchestrates lateral induction and lateral inhibition in the developing inner ear. Development (Cambridge) 141: 2313–2324

Phillips NE, Manning CS, Pettini T, Biga V, Marinopoulou E, Stanley P, Boyd J, Bagnall J, Paszek P, Spiller DG et al (2016) Stochasticity in the miR-9/Hes1 oscillatory network can account for clonal heterogeneity in the timing of differentiation. Elife 5

Ramos C, Rocha S, Gaspar C, Henrique D (2010) Two Notch ligands, Dll1 and Jag1, are differently restricted in their range of action to control neurogenesis in the mammalian spinal cord. PLoS One 5: e15515

Sagner A, Briscoe J, 2019. Establishing neuronal diversity in the spinal cord: A time and a place. Company of Biologists Ltd.

Sagner A, Gaber ZB, Delile J, Kong JH, Rousso DL, Pearson CA, Weicksel SE, Melchionda M, Mousavy Gharavy SN, Briscoe J et al (2018) Olig2 and Hes regulatory dynamics during motor neuron differentiation revealed by single cell transcriptomics. PLOS Biology 16: e2003127–e2003127

Shaya O, Sprinzak D, 2011. From Notch signaling to fine-grained patterning: Modeling meets experiments. pp. 732–739.

Shimojo H, Isomura A, Ohtsuka T, Kori H, Miyachi H, Kageyama R (2016) Oscillatory control of Delta-like1 in cell interactions regulates dynamic gene expression and tissue morphogenesis. Genes & development 30: 102–116

Shimojo H, Kageyama R, 2016. Oscillatory control of Delta-like1 in somitogenesis and neurogenesis: A unified model for different oscillatory dynamics. Academic Press, pp. 76–82.

Sonnen KF, Lauschke VM, Uraji J, Falk HJ, Petersen Y, Funk MC, Beaupeux M, François P, Merten CA, Aulehla A (2018) Modulation of Phase Shift between Wnt and Notch Signaling Oscillations Controls Mesoderm Segmentation. Cell 172: 1079–1090.e1012

Soto X, Biga V, Kursawe J, Lea R, Doostdar P, Thomas R, Papalopulu N (2020) Dynamic properties of noise and Her6 levels are optimized by miR‐9, allowing the decoding of the Her6 oscillator. The EMBO Journal

Sprinzak D, Lakhanpal A, Lebon L, Santat LA, Fontes ME, Anderson GA, Garcia-Ojalvo J, Elowitz MB (2010) Cis-interactions between Notch and Delta generate mutually exclusive signalling states. Nature 465: 86–90

Tiedemann HB, Schneltzer E, Beckers J, Przemeck GKH, Hrabě de Angelis M (2017) Modeling coexistence of oscillation and Delta/Notch-mediated lateral inhibition in pancreas development and neurogenesis. Journal of Theoretical Biology 430: 32–44

Tsiairis CD, Aulehla A (2016) Self-Organization of Embryonic Genetic Oscillators into Spatiotemporal Wave Patterns. Cell 164: 656–667

Vilas-Boas F, Fior R, Swedlow JR, Storey KG, Henrique D (2011) A novel reporter of notch signalling indicates regulated and random Notch activation during vertebrate neurogenesis. BMC Biol 9: 58

Webb AB, Lengyel IM, Jörg DJ, Valentin G, Jülicher F, Morelli LG, Oates AC (2016) Persistence, period and precision of autonomous cellular oscillators from the zebrafish segmentation clock. eLife 5

