## Supplementary figures and images for "A dynamic, spatially periodic, micro-pattern of HES5 underlies neurogenesis in the mouse spinal cord"

### Figure EV1

#### EV1. Related to Fig.1

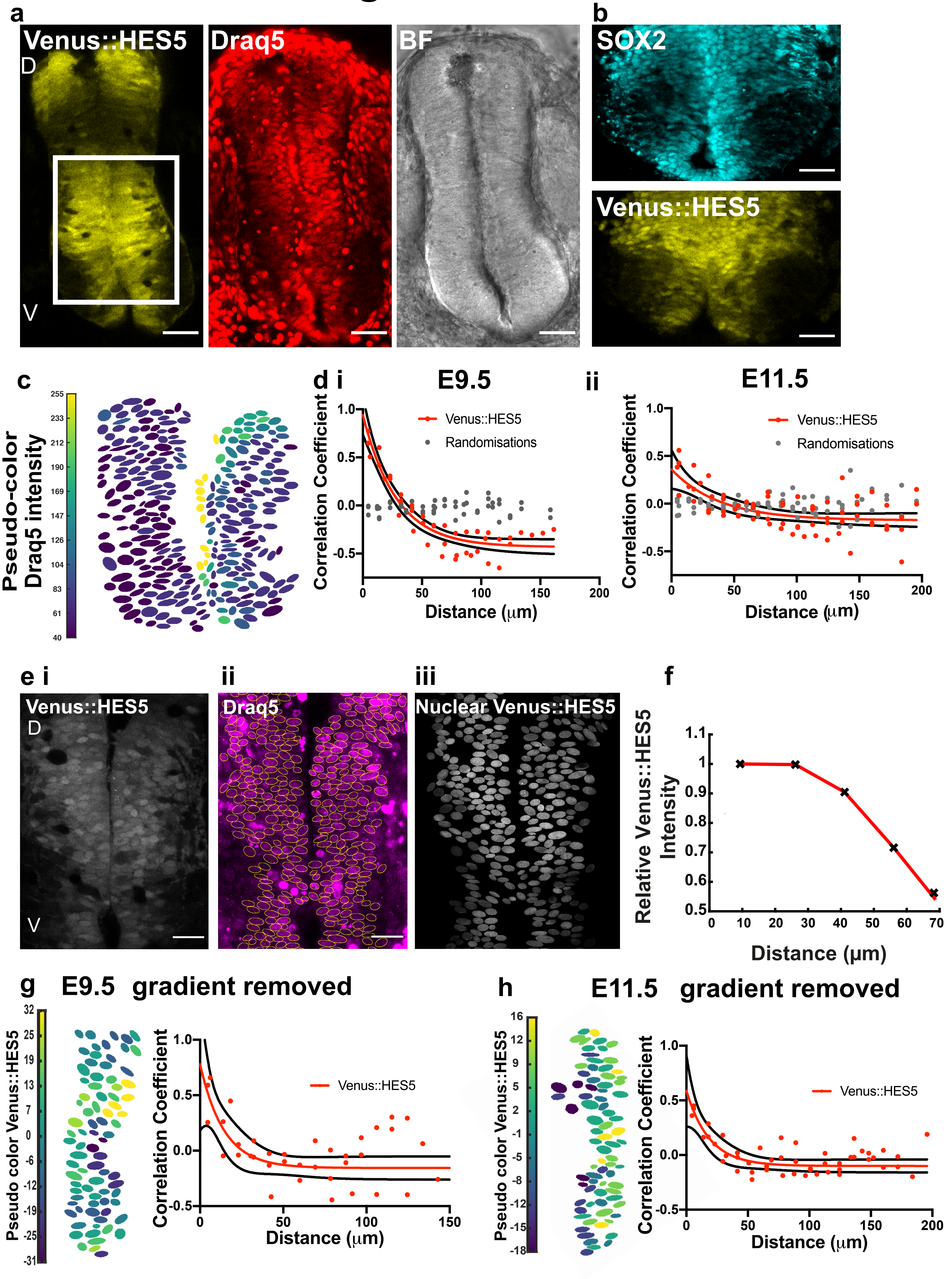

### Figure EV2

#### EV2. Related to Fig.2

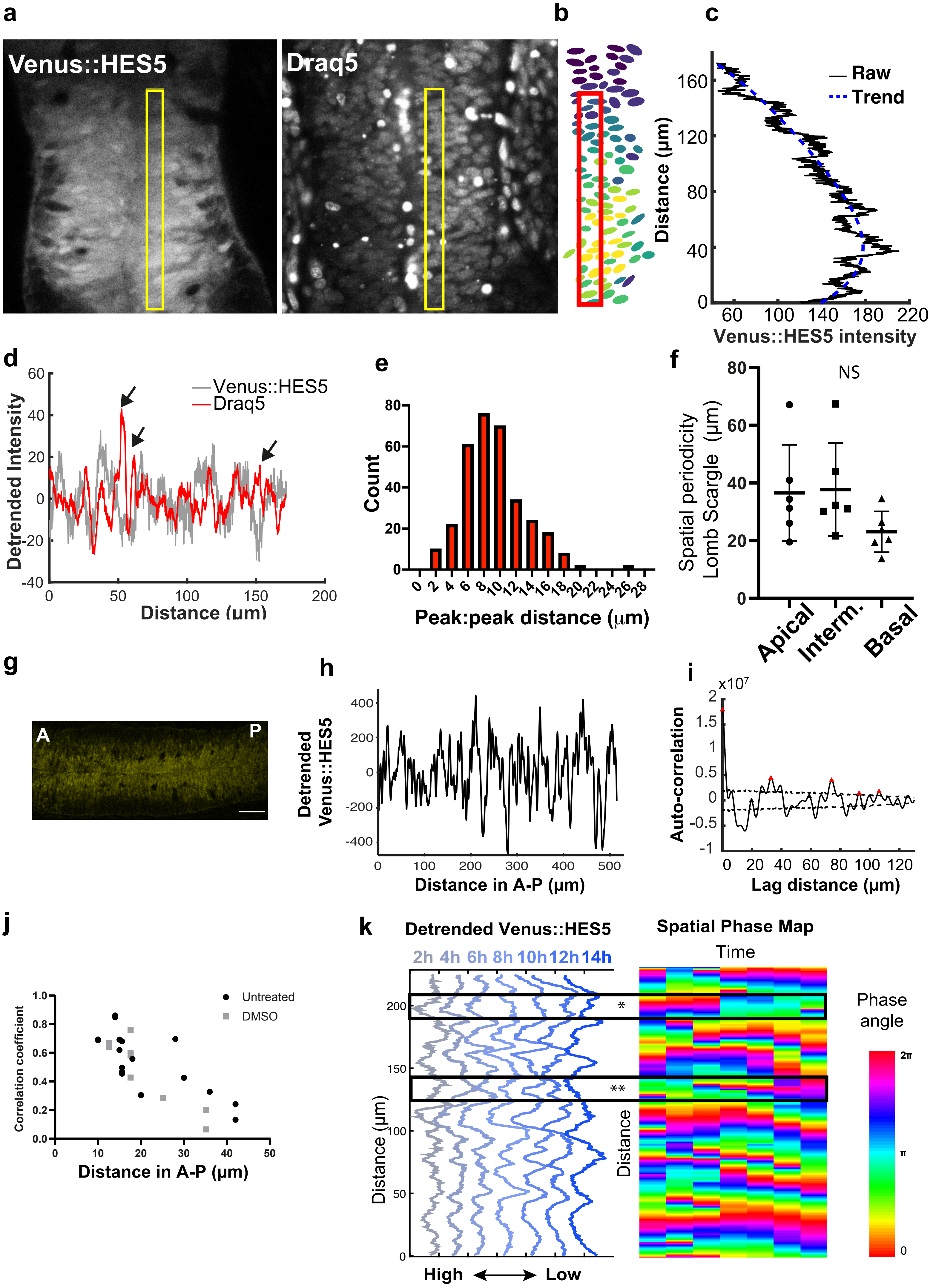

### Figure EV3

#### Figure EV3. Related to Figure 3.

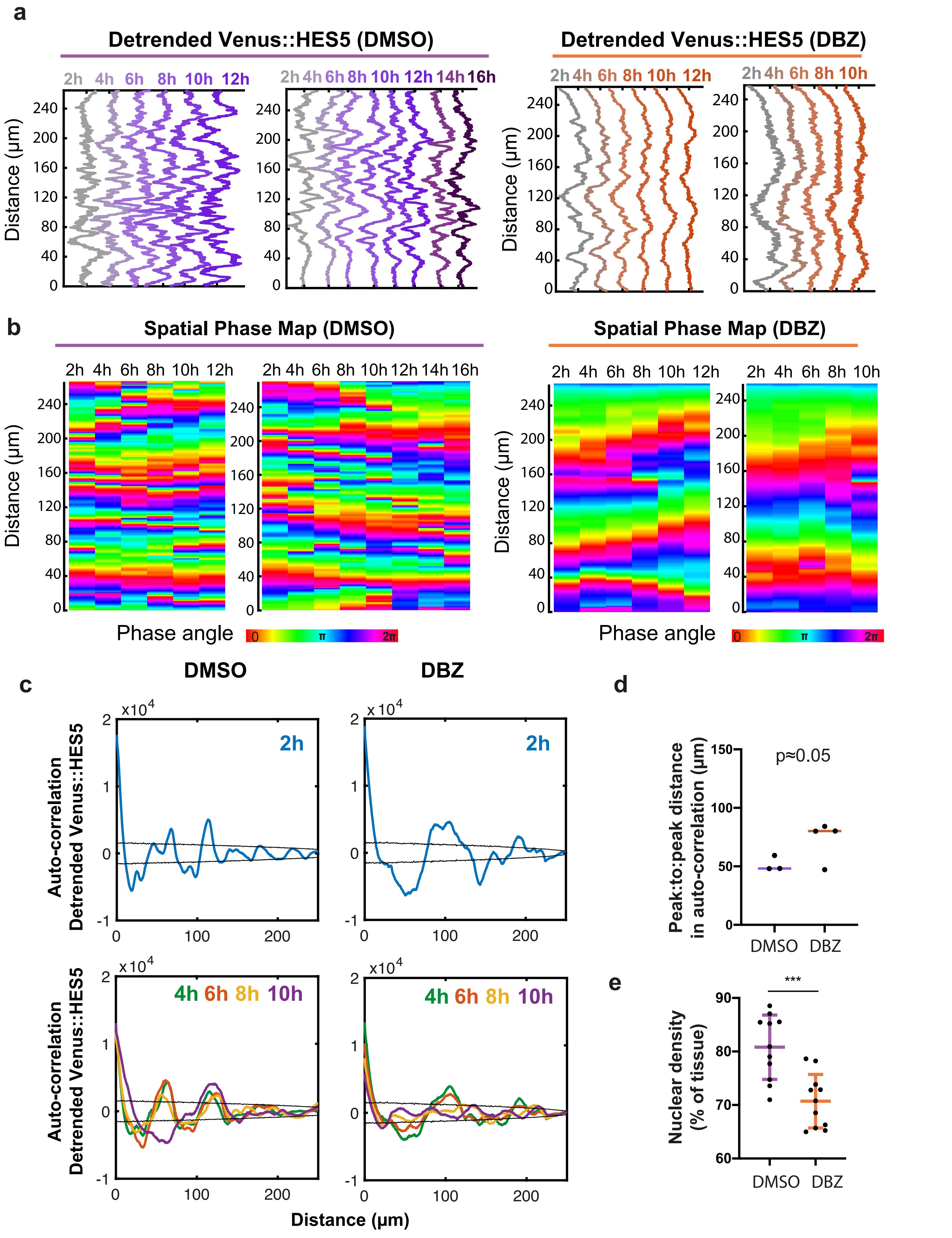

### Figure EV4

#### EV4. Related to Figures 5 and 7

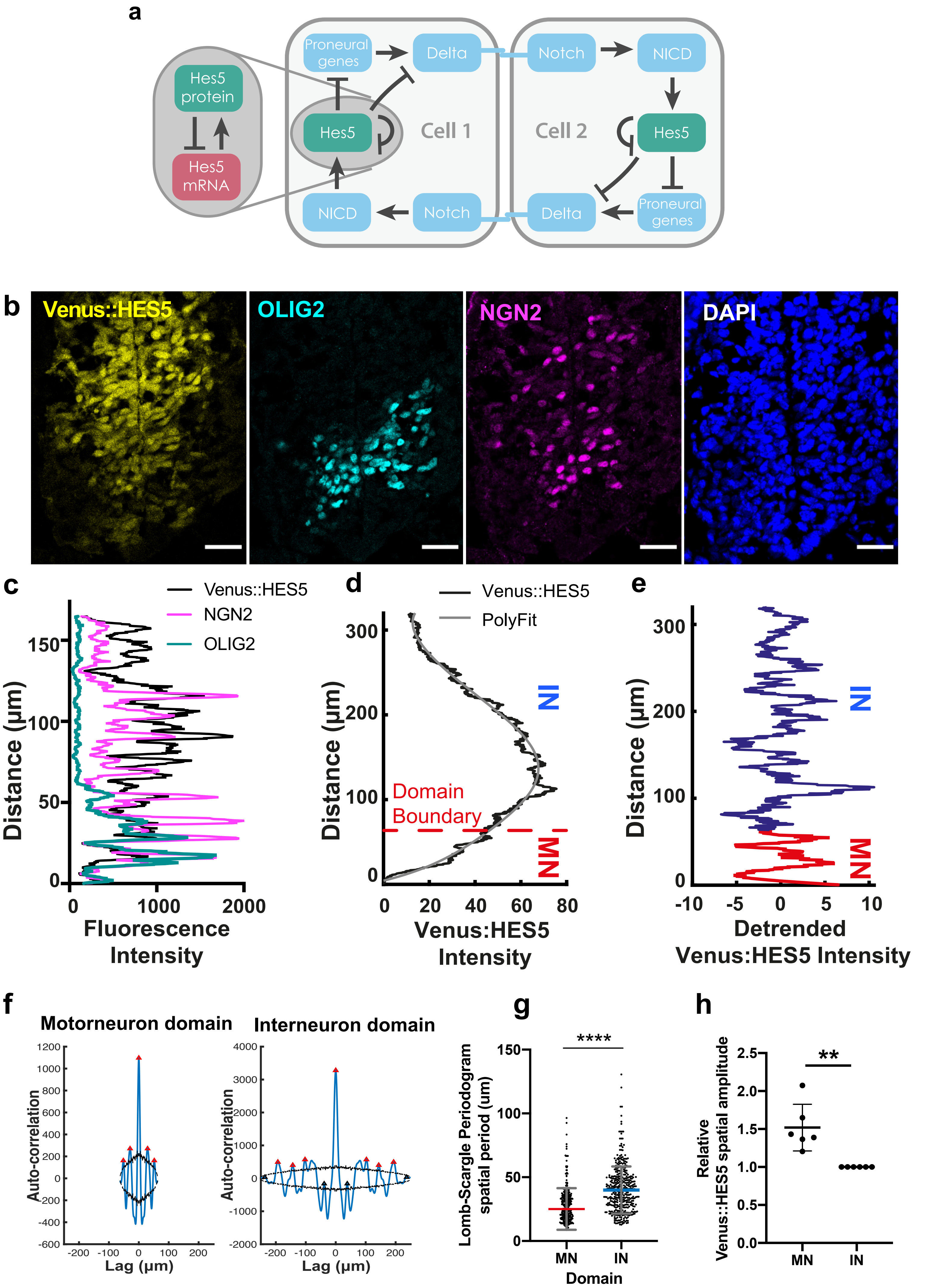

### Figure EV5

#### EV5. Related to Fig. 8

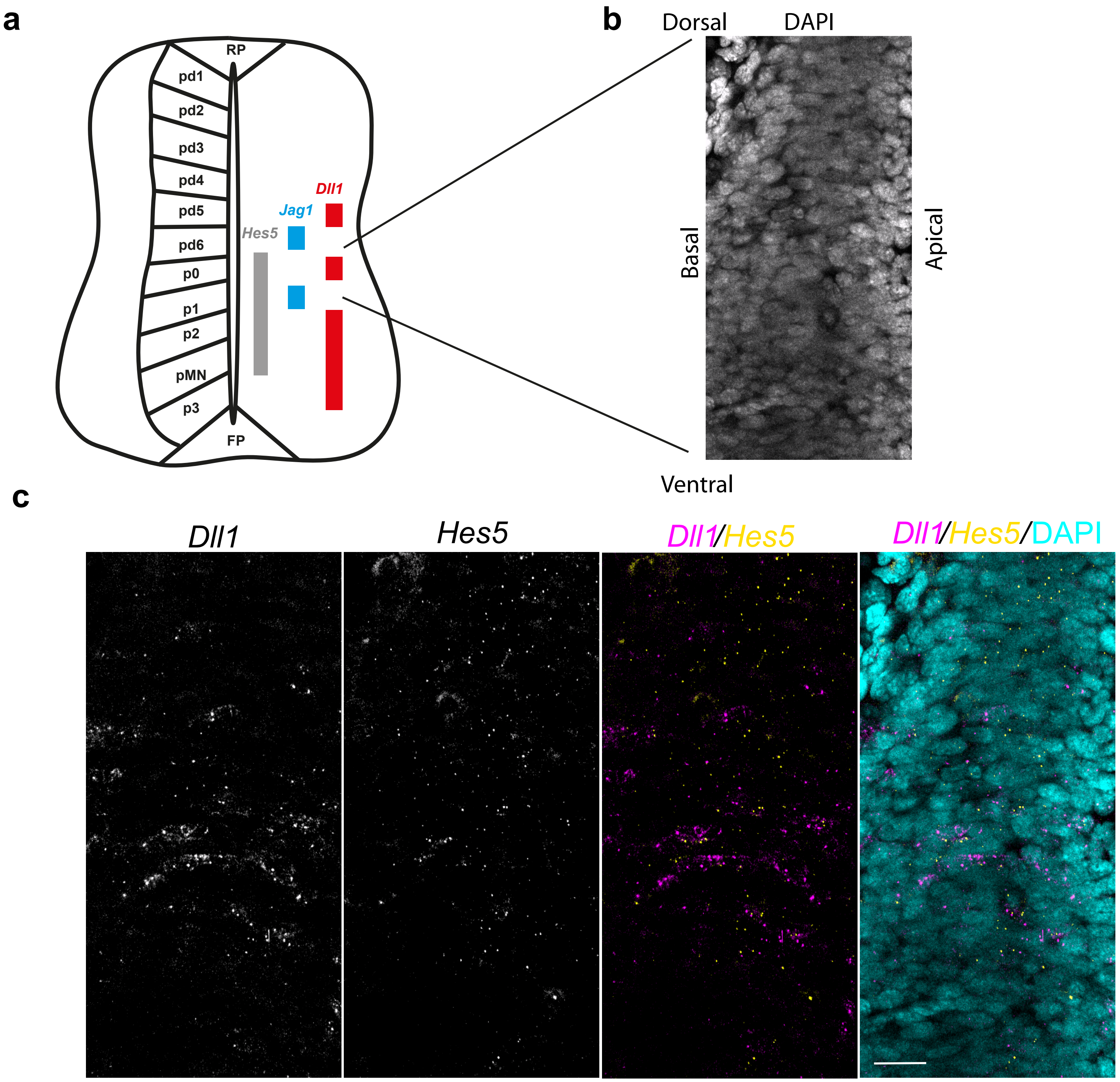
