## Appendix for "A dynamic, spatially periodic, micro-pattern of HES5 underlies neurogenesis in the mouse spinal cord"

#### Appendix Figures

### Appendix Figure S1. Related to Figure 2.

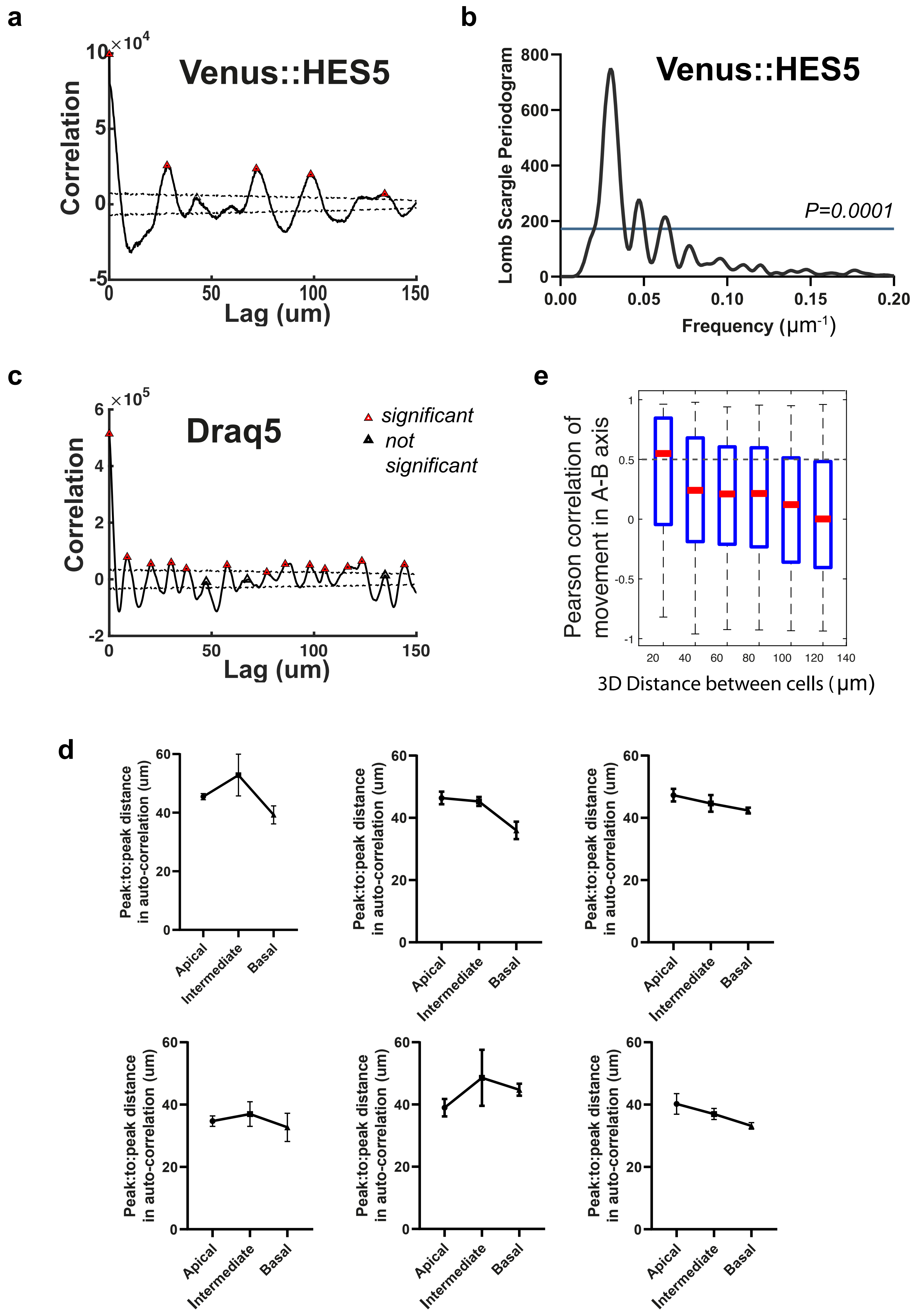

### Appendix Figure S2. Related to Figure 3.

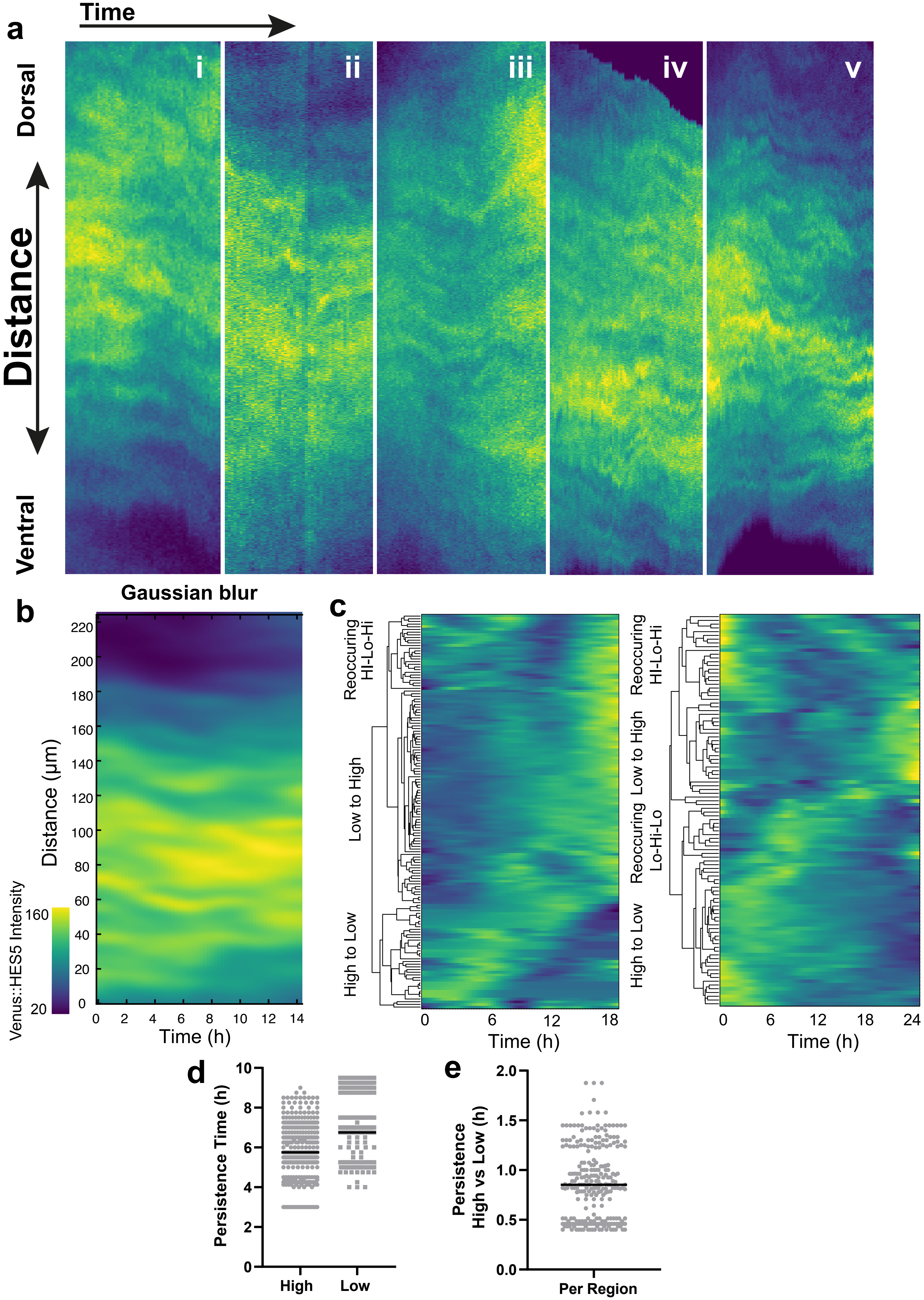

### Appendix Figure S3. Related to Fig. 3

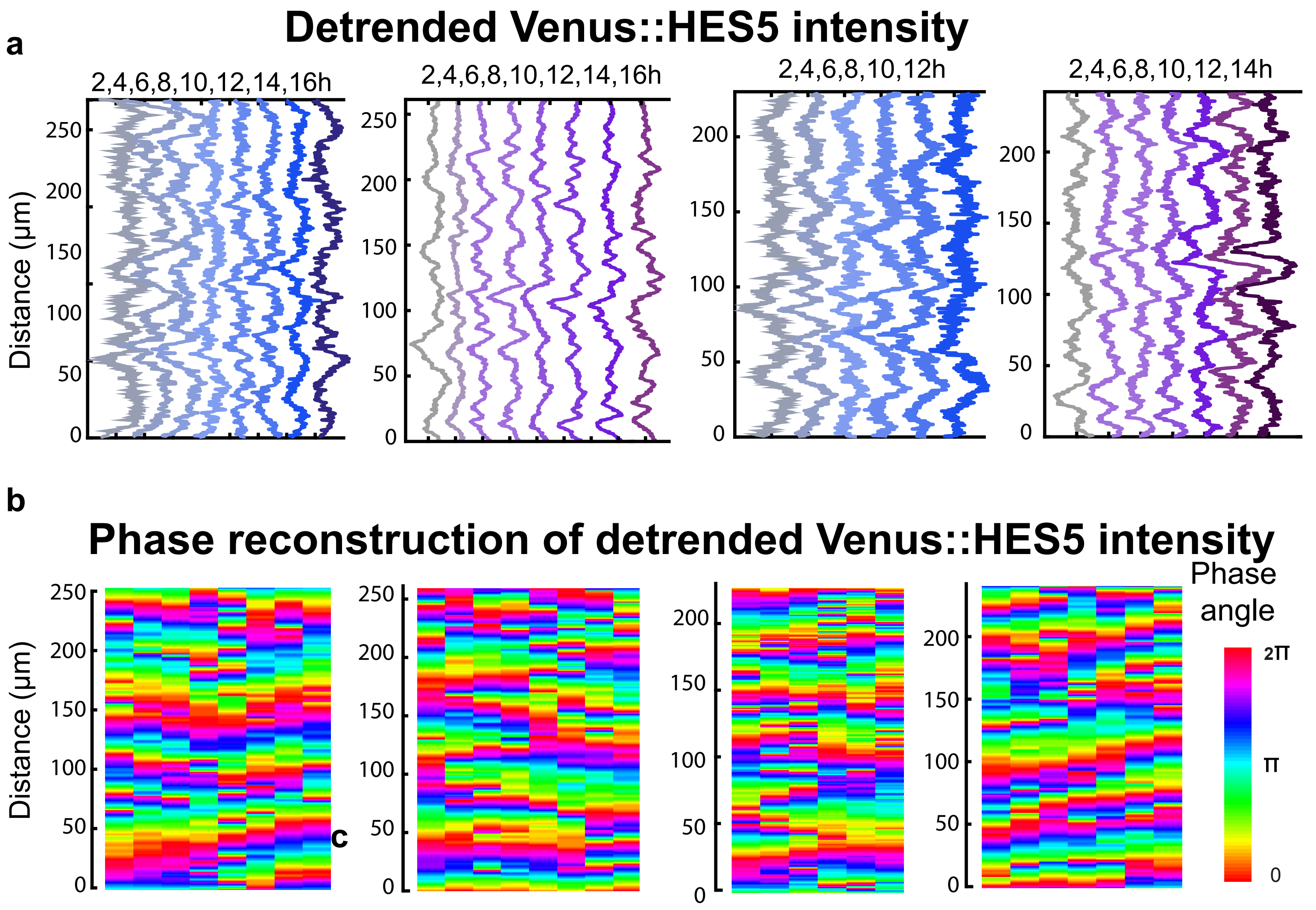

### Appendix Figure S4. Related to Figs 3 and 5.

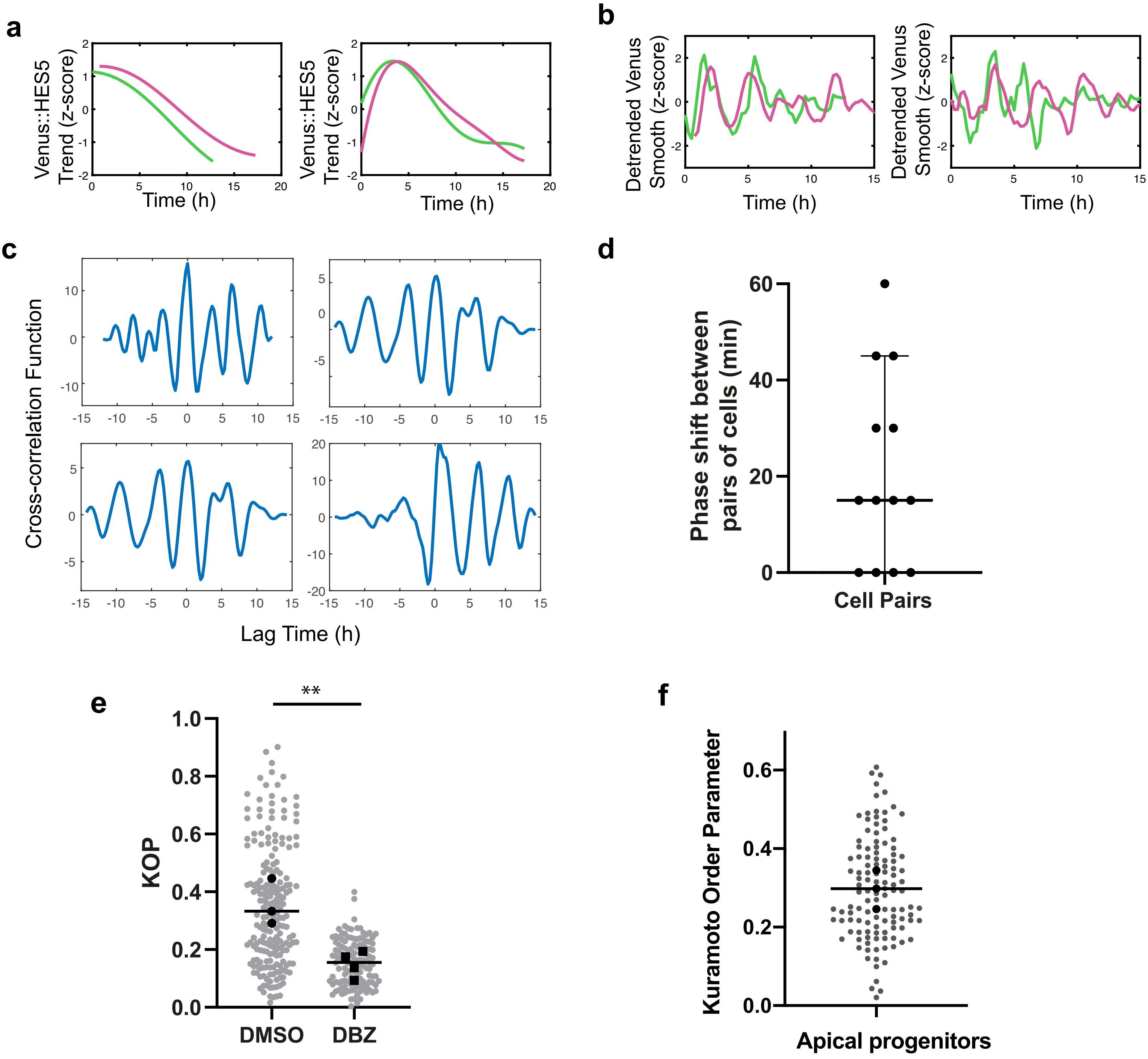

### Appendix Figure S5. Related to Figure 5.

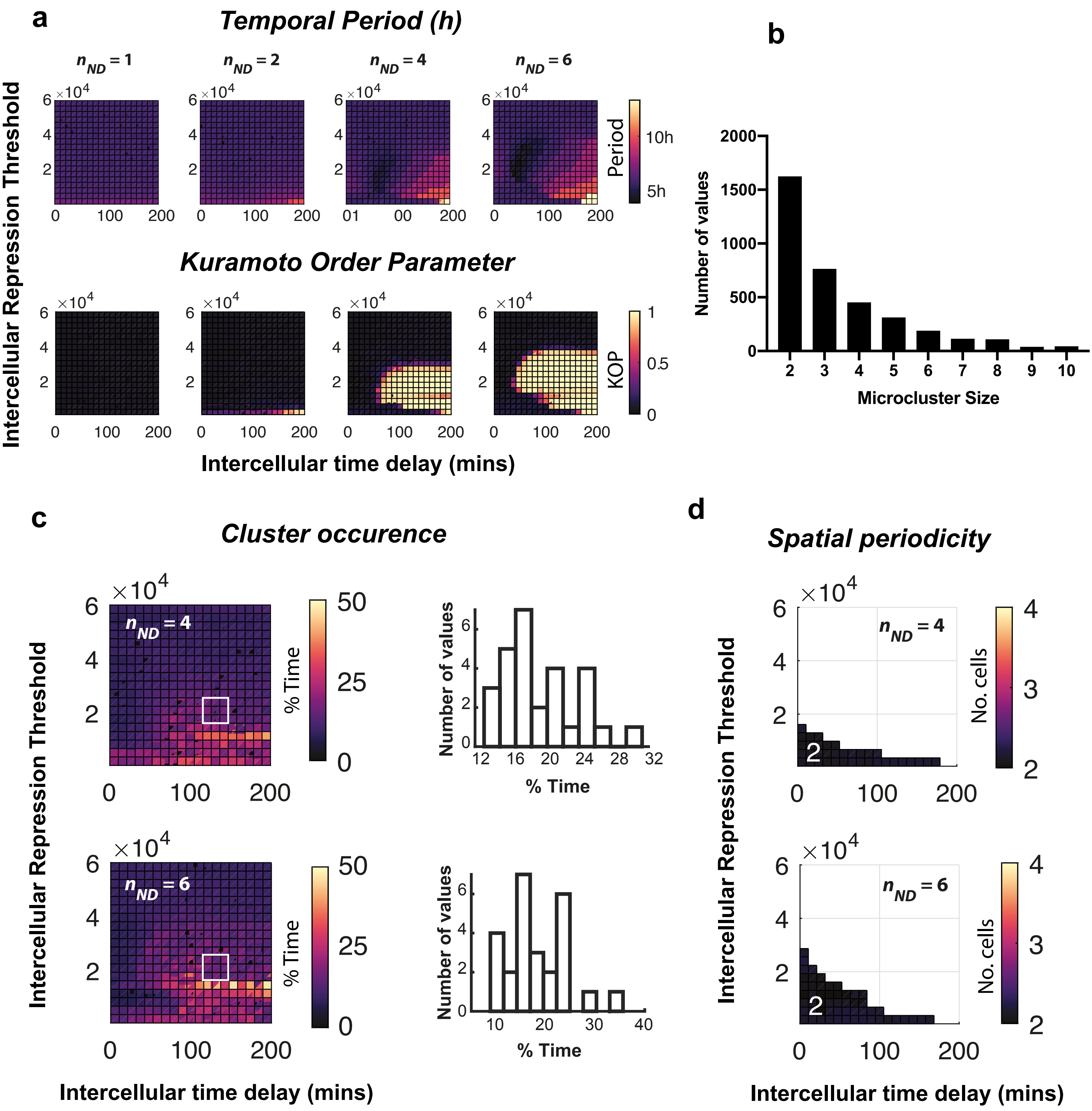

### Appendix Figure S6. Related to Figure 5.

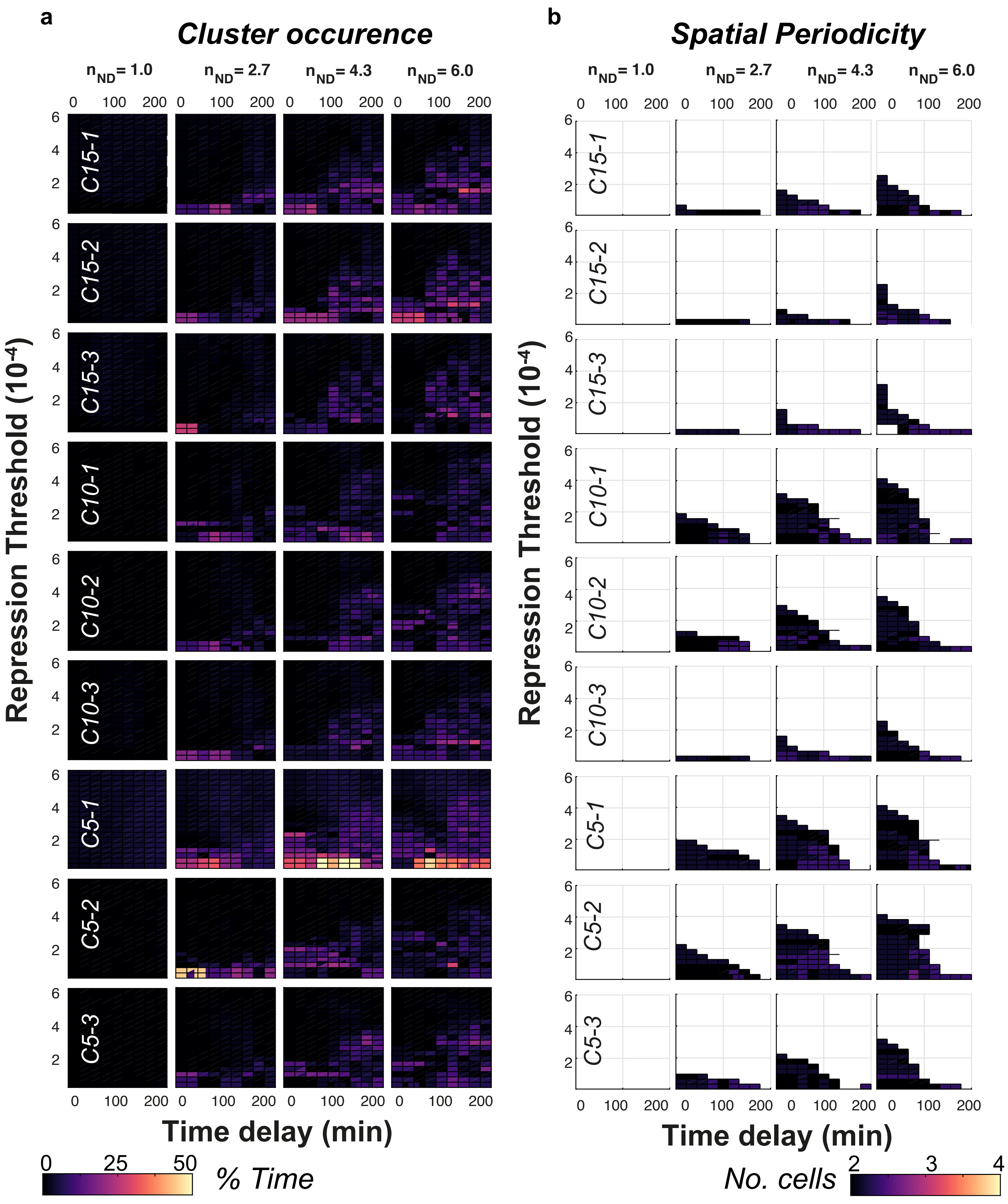

### Appendix Figure S7. Related to Figs. 7 and 8

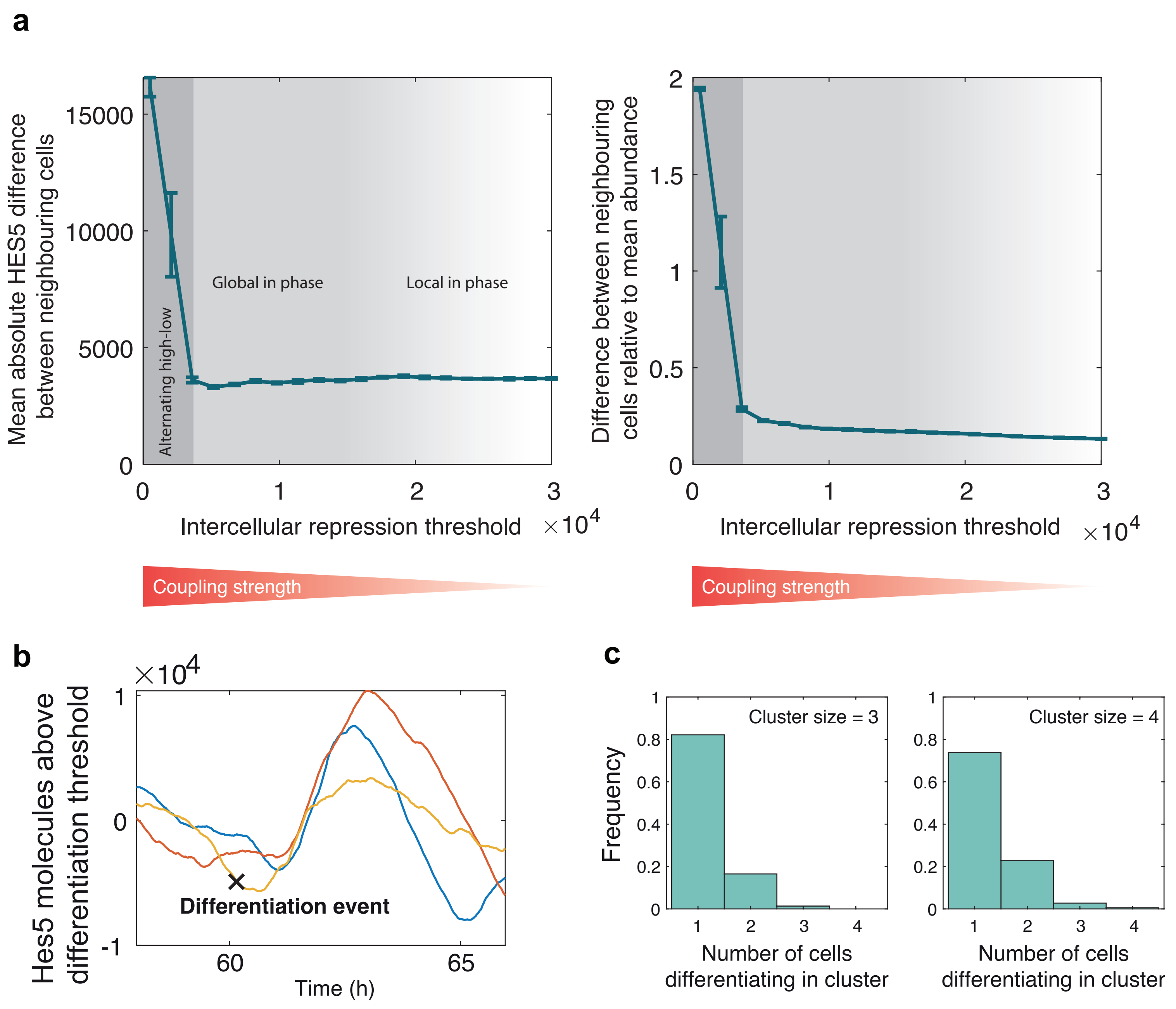

### Appendix Figure S8

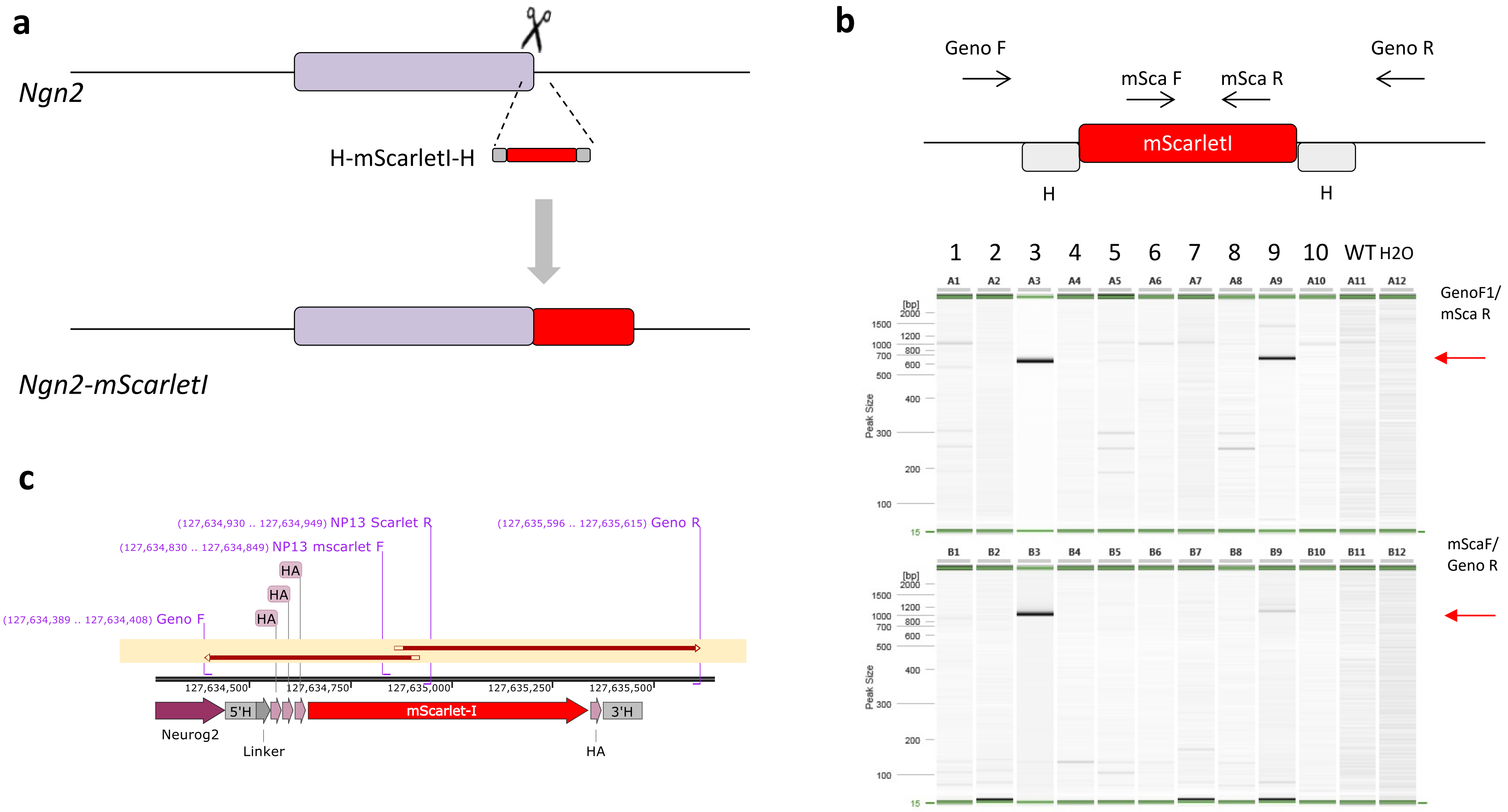

**Appendix Table S1.**
Quantification of number of kymographs per independent experiment showing spatial periodicity following an auto-correlation test with >95% statistical significance.

| Untreated Experiment | No of kymographs LHS* +RHS** | Number of kymographs at specific timepoint divided by total number of kymographs per experiment | | | | | | % Overall |
|---|---|---|---|---|---|---|---|---|
| | | 2h | 4h | 6h | 8h | 10h | 12h | |
| E 1 | 3LHS+3RHS | 6/6 | 6/6 | 6/6 | 6/6 | 6/6 | 6/6 | 100 |
| E 2 | 4LHS+4RHS | 8/8 | 8/8 | 8/8 | 8/8 | 8/8 | 8/8 | 100 |
| E 3 | 3LHS+3RHS | 6/6 | 6/6 | 6/6 | 6/6 | 6/6 | 8/8 | 100 |
| E 4 | 3LHS+3RHS | 6/6 | 6/6 | 6/6 | 6/6 | 6/6 | 6/6 | 100 |
| E 5 | 4LHS+3RHS | 7/7 | 7/7 | 7/7 | 7/7 | 7/7 | 7/7 | 100 |
| E 6 | 3LHS+3RHS | 6/6 | 6/6 | 6/6 | 6/6 | 6/6 | 6/6 | 100 |

**Appendix Table S2.**
Quantification of drift in Y-axis (Dorsal to Ventral) from single cell data corresponding to kymograph experiments.

| Condition | Average displacement in Y-axis per 12h | Accepted < 20 µm |
|---|---|---|
| Untreated 1 (E 1) | 8.41 µm | Accepted |
| Untreated 2 (E 5) | 19.86 µm | Accepted |
| Untreated 3 (E 6) | 22.805 µm | *Rejected* |
| DMSO 1 | 18.855 µm | Accepted |
| DMSO 2 | 19.84 µm | Accepted |
| DMSO 3 | 11.12 µm | Accepted |
| DBZ 1 | 13.045 µm | Accepted |
| DBZ 2 | 14.585 µm | Accepted |
| DBZ 3 | 17.205 µm | Accepted |
| DBZ 4 | 15.265 µm | Accepted |

##### Appendix Table S3.

Model parameters used.

| | | Single-cell parameters | | | | | | | Multicellular parameters | | |
|---|---|---|---|---|---|---|---|---|---|---|---|
| | | Transcription rate | Translation rate | Repression threshold | Self-repression time delay | Self-repression Hill coefficient | mRNA degradation rate | Protein degradation rate | Intercellular repression threshold | Intercellular time delay | Intercellular Hill coefficient |
| Coherence | Parameter set name | $\alpha_m$ $(min^{-1})$ | $\alpha_p$ $(min^{-1})$ | $P_{H0}$ (# of proteins) | $\tau_H$ (min) | $n_H$ | $\mu_m$ $(min^{-1})$ | $\mu_p$ $(min^{-1})$ | $P_{ND0}$ (# of proteins) | $\tau_{ND}$ (min) | $n_{ND}$ |
| 15% | **Main** | **0.8** | **26.4** | **25088** | **20** | **3.6** | $\frac{ln(2)}{30}$ | $\frac{ln(2)}{90}$ | $0-60000$ | $0-200$ | $1-6$ |
| | C15a | 8.2 | 8.6 | 17914 | 23 | 2.7 | | | | | |
| | C15b | 26.9 | 9.4 | 10902 | 20 | 2.6 | | | | | |
| | C15c | 7.3 | 26.9 | 10733 | 24 | 2.3 | | | | | |
| 10% | C10a | 1.3 | 13.7 | 47482 | 36 | 2.6 | | | | | |
| | C10b | 2.0 | 11.6 | 35319 | 19 | 2.5 | | | | | |
| | C10c | 8.0 | 13.5 | 10255 | 19 | 2.0 | | | | | |
| 5% | C5a | 0.7 | 18.5 | 54656 | 38 | 2.6 | | | | | |
| | C5b | 2.0 | 5.8 | 66657 | 28 | 2.1 | | | | | |
| | C5c | 2.5 | 8.8 | 26823 | 22 | 2.0 | | | | | |

### Appendix Figure legends

##### Appendix Figure S1. Related to Figure 2.

##### Draq5 and Venus::HES5 spatial periodicity in live spinal cord E10.5 tissue cultures.

**(a)** Auto-correlation plot of Venus::HES5 and Draq5 from the same tissue profile; black dashed lines show confidence limits for peak significance based on bootstrap approach on the intensity profile (see Methods); red triangle - significant peak, black triangle – non-significant peak; multiple significant peaks in auto-correlation shows periodicity in spatial profile. **(b)** Representative examples of Lomb Scargle periodogram averaged over multiple slices in the same experiment; line indicates statistical significance with p value 0.0001. **(c)** Pearson correlation coefficients between tracked AP-position time series of pairs of cells plotted against mean 3D distance between them; x-values indicate the right edge of the data bin for each box plot shown; data consists of multiple timeseries of positional information acquired from approx. 100 single cells tracked in E10.5 spinal cord in n=3 experiments. **(d)** Mean peak to peak distances per experiment in apical, medium and basal regions (10, 30 and 60 μm from ventricle respectively); bars indicate mean and SD per experiment; datasets consisted of at least 3 z-sections and both left and right sides of ventricle analysed in n=6 experiments.

**Appendix Figure S2. Related to Figure 3.**

**Venus::HES5 spatio-temporal behaviour in apical region of E10.5 spinal cord tissue.**

**(a)** Kymographs of Venus::HES5 expression profiles observed along ventral-dorsal direction in a 15μm wide apical region; panels indicate 5 example slice cultures in addition to example in **Figure 3b**. **(a)** Kymograph obtained from data shown in **Figure 3b** after a 2μm Gaussian blur filter has been applied (see Methods). **(c)** Representative examples showing clustering of Venus::HES5 z-scored intensities (additional to **Figure 3c**) collected from 20um regions; these diagrams indicate that expression in every region is dynamic corresponding to High-to-Low, Low-to-High or Re-occuring cluster activity; one diagram per independent experiment. **(d)** Persistence time computed in regions showing a cluster in the initial 2h interval (see Appendix, zero-crossing method) showing (left panel) time intervals spent above the mean (High) and below the mean (Low) in kymograph data; and (right panel) the ratio of time in High vs Low in the same region; dots represent individual regions from 5 independent experiments and multiple z-stacks. **(e)** Persistence High to Low ratio in the same region observed in areas of the tissue showing a cluster in the initial 2h interval; uses data also included in **(d).**

**Appendix Figure S3. Related to Figure 3.**

**Venus::HES5 spatio-temporal expression and phase mapping in E10.5 apical region.**

**(a)** Spatio-temporal plots showing Venus::HES5 intensities detrended with a polynomial order 4 observed along ventral-dorsal direction; signals obtained by averaging kymograph data over 2h time intervals; panels represent 4 example slice cultures in addition to example in **Figure EV2k-left**. **(b)** Spatial phase map reconstruction using Hilbert (see Methods) obtained from detrended Venus::HES5 data shown in **(a)**; panels correspond to data in **(a)** and are additional to the one in **Figure EV2k-right.**

**Appendix Figure S4. Related to Figure 3.**

**Co-ordination in ultradian dynamics observed in Venus::HES5 expressing progenitors tracked in E10.5 spinal cord tissue ex-vivo slice cultures.**

**(a,b)** Gaussian Process fitting of single cell timeseries shown in **Figure 3i,k** using methods described in Manning *et al.* 2019; Phillips *et al.* 2016 showing **(a)** z-score of Venus::HES5 trends; and **(e)** z-score of detrended Venus::HES5 over time. **(c)** Representative examples of cross-correlation analysis (see Methods) observed between pairs of neighbouring single cell detrended Venus::HES5 timeseries in spinal cord tissue. **(d)** Phase-shift distribution obtained from detrended Venus::HES5 in neighbouring single cells pairs from tissue; dots represent pairs of neighbouring cells, lines indicate median with interquartile range; median is 15 min which corresponds to the sampling rate. **(a)** Comparison of phase synchronisation measured as population Kuramoto Order Parameter (KOP, see Methods) values measured in single cell progenitors from tissue treated with DMSO (control) and Notch inhibitor DBZ; gray dots indicate individual timepoints between 0 to 10-12h with black dots indicating average per slice; bar represents mean of (DMSO, n=3) and (DBZ, n=4) slices; unpaired 2-tailed t test with p=0.0068**. **(b)** Quantification of phase synchronisation measured as population KOP in sparsely tracked apical progenitor cells in ex-vivo E10.5 spinal cord slice cultures; gray dots indicate values at individual timepoints between 0 to 12-14h with black dots indicating average per slice; bar represents mean of 3 slices.

**Appendix Figure S5. Related to Figure 5.**

**Multi-parameter exploration of synthetic multicellular HES5 model.**

**(a)** Explorations of the inter-cellular parameters determining the repressive interactions between cells via a Hill function (**Figure 5b**): intercellular time delay (x-axes), repression threshold (y-axes), and Hill coefficient (plotted as separate maps in each column); pseudo-color indicates (top panels) average temporal period of HES5 oscillations (in hours) and (bottom panels) average KOP values from 5 independent simulations per parameter combination; statistics were collected from synthetic HES5 timeseries using a 26 by 10 grid size per simulation with length of 100 hours and generated using the stochastic model with randomised initial conditions. Single cell HES5 model parameter set listed in **Appendix Table S3-Main**. **(b)** Microcluster diameter measurements obtained from synthetic weakly coupled cells corresponding to repression threshold values between 20,000 to 24,000 (see Methods); dataset represents the frequency distribution of microclusters detected in 250 synthetic data kymographs of 80 hours length; **(c)** Inter-cellular parameter exploration of frequency of cluster detection (fraction of time points in a simulation where clusters are detected, see Methods); panels correspond to Hill coefficient values of 4 (top) and 6 (bottom); values observed within a weak coupling range are presented as frequency plots to the right. **(d)** Inter-cellular parameter exploration of spatial periodicity for Hill coefficient values 4 (top) and 6 (bottom); spatial periods (as cell number) found significant using a Fisher G significance test (see Methods). Statistics in **(c,d)** represent average of 5 independent simulations collected from a grid of 26 by 10 synthetic HES5 timeseries per simulation with length of 100 hours and generated using the stochastic model with randomised initial conditions.

**Appendix Figure S6. Related to Figure 5.**

**Extensive search over multiple single-cell parameter sets shows no additional model behaviour at lower coherence values.**

Multi-parametric exploration of HES5 inter-cellular and intra-cellular dynamics used to investigate microcluster occurrence **(a)** and spatial periodicity **(b)**. Panels represent maps of intercellular time delay (x-axes), repression threshold (y-axes), and Hill coefficient (plotted as separate maps in each column). Panels arrayed horizontally correspond to distinct HES5 auto-repression parameter sets marked as C15, C10 and C5 with 3 repeats (e.g. C15-1 to 3) corresponding to low coherence (15% to 5%) dynamics where low coherence indicates reduced oscillatory activity in the population. Parameter sets used are presented in **Appendix. Table S3**. Pseudo-color indicates **(a)** frequency of cluster detection (fraction of time points in a simulation where clusters are detected, see Methods) and **(b)** spatial period (as cell number) found significant using a Fisher G significance test (see Methods); statistics were collected from synthetic HES5 timeseries using a 26 by 10 grid size per simulation with length of 100 hours and generated using the stochastic model with randomised initial conditions.

**Appendix Figure S7. Related to Figures 7 and 8.**

**Exploration of cell:cell HES5 abundance differences and differentiation within a cluster of locally in-phase cells using the multicellular synthetic HES5 model.**

(**a**) Analysis of the multicellular model shows that protein abundance differences between pairs of neighbouring cells (immediate neighbours) increases sharply at higher coupling strength/lower intercellular repression threshold; (Left panel) absolute concentration differences between pairs of neighbours and (right panel) concentration differences relative to the mean population Hes5 expression. Synthetic data was generated from 26 by 10 grid of cells run for 100h and for each repression threshold 10 repeats were run (SD shown by error bars) . **(b)** Example time traces plotted from a cluster of 3 locally-synchronised cells (KOP>0.9 over 8h window) with an example differentiation event highlighted, which occurs only in one cell out of the three (details of how a differentiating cell is defined is given in **Appendix Methods: Rate of differentiation from synthetic data**). **(c)** Frequency of differentiating cell numbers in a cluster size of 3 (left panel) and cluster size of 4 (right panel). The most likely number of differentiating cells was found to be 1, with decreasing probability for increasing numbers of differentiating cells (within a time frame of 8h). Simulation run to generate frequency plots used a grid size of 26 by 20, run for 300h.

**Appendix Figure S8. Related to Figure 8.**

**Generation of Neurog2::mScarlet-I knock-in mouse**

**(a)** Targeting design for CRISPR mediated knock in. Scissors indicate CRISPR target site over STOP codon of *Ngn2*, and homology flanked mScarletI long ssDNA donor below. Homology direct repair (HDR) results in the integration of mScarletI tag in frame with final exon of *Ngn2*. (**b)** Genotyping by PCR strategy, primer locations are indicated on the schematic. Images are PCR reactions run on Qiaxcel with red arrows indicating correct HDR product size. **(c)** Sanger sequence alignment of pup 3 from two overlapping sequencing reactions (forward and reverse).

#### Appendix Supplementary Methods 1-Neurog2 knock-in mouse

**Generation of Neurog2::mScarlet-I mice**

We used the EASI-CRISPR strategy (Quadros et al, 2017) to generate a C-terminally tagged Ngn2-mScarletI mouse. Two sgRNA targeting the STOP codon of the Ngn2 gene were selected using the Sanger WTSI website (http://www.sanger.ac.uk/htgt/wge/; Hodgkins et al, 2015) that adhered to our criteria for off target predictions (guides with mismatch (MM) of 0, 1 or 2 for elsewhere in the genome were discounted, and MM3 were tolerated if predicted off targets were not exonic). sgRNA sequences (GGGACTGTATCTAGAGCTGC-*GGG* and GCAGCTCTAGATACAGTCCC-*TGG*) were purchased as crRNA oligos, which were annealed with tracrRNA (both oligos supplied by Integrated DNA Technologies) in sterile, RNase free injection buffer (TrisHCl 1mM, pH 7.5, EDTA 0.1mM) by combining 2.5 $\mu$g crRNA with 5 $\mu$g tracrRNA, heating to 95$^o$C, and slowly cooling room temperature. To generate the long single strand DNA donor repair template (**Appendix Figure S8a)**, a homology flanked flexible linker-3xHA-mScarletI-HA DNA sequence was cloned and used as a template in an initial PCR reaction with primers Ngn2_lssDNA_F catgctcccggccgccatgggaattcgagctcgtggactactggcagcccccacctccggagaagcatcgtta and Ngn2 lssDNA R ctgctctgtgaagtggagtccggggcgtggagggtgaggggcgggagaaggatggaagagggccgggtagaggacgag agagggagacccgcagctttaagcgtaatctggaacatcgtatgggt. This amplicon appended the universal pGEM_dual_Bio_F primer sequence (catgctcccggccgccatg) at the 5' end, and the lssDNA comprising 5'H(76nt)-linker-3xHA-ScarletI-HA-3'H(96nt) was purified as described (Bennett et al, 2020).

For embryo microinjection the annealed sgRNA were complexed with EnGen Cas9 protein (New England Biolabs) at room temperature for 10 minutes, before addition of long ssDNA donor (final concentrations; each sgRNA 20 ng/μl, Cas9 protein 40 ng/μl, lssDNA 10 ng/μl). The injection mix was directly microinjected into C57BL6/J (Envigo) zygote pronuclei using standard protocols. Zygotes were cultured overnight and the resulting 2 cell embryos surgically implanted into the oviduct of day 0.5 post-coitum pseudopregnant mice. Potential founder mice were screened by PCR, using primers that flank the Homology arms (Geno F aactccacgtccccatacag, Geno R cagtcttacgaggttcccca), used in combination with internal mScarletI primers (mScarletI R gtcctcgaagttcatcacgc, mScarletI F tcccctcagttcatgtacgg) (**Appendix Figure S8b**; middle and lower panel). A single pup (#3) was PCR positive for both reactions, and Sanger sequencing (**Appendix Figure S8c**) confirmed perfect integration of the gene tag. Note pup #9 demonstrated a positive PCR result for the first reaction, and a weaker amplification product for the second reaction which we could not re-amplify with high fidelity polymerases. Pup #3 was bred forward with a WT C57BL6/J mouse, germline transmission confirmed through PCR and sequencing, and a colony established.

#### Appendix Supplementary Methods 2- Model

##### Stochastic model of auto-repression coupled between cells

We adapted a HES5-parameterised single-cell model from previous work into a Notch-Delta-coupled multicellular model (**Figure 5a**) (Manning *et al.*, 2019). The single cell network is the basic unit of the model and consists of negative feedback loop of HES5 protein onto its own mRNA expression described by stochastic delay differential equations (SDDEs) capable of producing stochastic autonomous oscillations. The SDDEs implement a Langevin approach over a Gillespie algorithm in favour of reduced computational cost and fixed time steps which enable easier addition of time delays (Gillespie, 2000).

To extend the single cell model to a multicellular one, the cells were coupled together with an interaction representative of the Notch-Delta pathway and its interaction with HES5. Instead of modelling every reaction in the chain of events that constitute the Notch-Delta pathway, we used a highly adjustable coupling function – an inhibitory Hill function – that requires only two parameters to specify its shape. This drastically reduced the potential number of parameters required to describe the Notch-Delta pathway that couples HES5 dynamics between cells while maintaining the ability to describe a wide variety of possible coupling function shapes that may represent the biological system. The Hill function in this model therefore describes how the production rate of HES5 in one cell is inhibited by high levels of HES5 expression in a neighbouring cell and has the form of

$$J(P) = \frac{1}{1 + \left(\frac{P}{P_0}\right)^n} \tag{1}$$

where $J(P)$is the mRNA production rate response of a cell to the concentration of protein $P$ in a neighbouring cell, $P_0$ is the repression threshold (also known as the effective concentration EC50) and $n$ is the Hill coefficient. $P_0$ and $n$ define the shape of the Hill function (illustrated in **Figure 5b**) and thus how a cell will respond to the levels of HES5 in its neighbouring cells. Specifically, we refer to repression threshold as the inverse of the coupling strength as this parameter defines at what concentrations of HES5 in a neighbouring cell a receiving cell becomes repressed. The Hill coefficient is also capable of changing the coupling strength but to a lesser extent as it affects the slope of the curve around the value of the repression threshold, creating a more step-like response at high Hill coefficient value and reproducing a Michaelis-Menten curve at a value of 1.

For the analysis in this paper, we chose neighbours to be taken as the closest or first rank neighbours, which in the case of hexagonal geometry implies a cell can be coupled to the dynamics of a maximum of 6 immediate neighbours (illustrated in **Figure 5c**). The incoming protein concentration that a cell experiences is defined as the average of its neighbours

$$\bar{P} = \frac{1}{N}\sum_{n=1}^{N} P_n \tag{2}$$

where $N$ is the number of neighbouring cells and $P_n$ is the protein concentration in neighbouring cell $n$. Equation (2) assumes that a cell receives equal contribution from each of its neighbouring cells, and there is no efficiency of signalling or scaling parameter in (2) as this would be redundant in addition to the intercellular repression threshold. Taking into account the number of neighbours ($\frac{1}{N}$ in (2)) rather than a static multiplication term reduces the effects of different patterning at the boundaries of the simulated tissue and was found to result in similar dynamics as when periodic boundaries were used.

Finally, to capture the dynamics of the Notch-Delta pathway more accurately, a time delay was added to the Hill function. The use of a nonlinear function to describe the multiple steps neglects the time-delays associated with the reaction, transport/diffusion, and synthesis, so an explicit time delay parameter was added. The full form of the coupling function taking into account the neighbours and time delay is given by

$$J(\bar{P}(t-\tau_{ND})) = \frac{1}{1+\left(\frac{\bar{P}(t-\tau_{ND})}{P_{ND0}}\right)^{n_{ND}}} \quad (3)$$

where the subscript $ND$ indicates intercellular parameters associated with Notch-Delta signalling and $\tau_{ND}$ is the intercellular time delay.

The full set of equations that describe the dynamics in an individual cell within the multicellular model is the single cell model from previous work (Manning *et al.*, 2019) but with the introduction of (3) as a multiplicative term that affects the production rate of mRNA

$$\frac{dM(t)}{dt} = -\mu_m M(t) + \alpha_m G(P(t-\tau_H))J(\bar{P}(t-\tau_{ND})) + \sqrt{\mu_m M(t) + \alpha_m G(P(t-\tau_H))J(\bar{P}(t-\tau_{ND}))}\xi_m(t) \quad (4)$$

$$\frac{dP(t)}{dt} = -\mu_p(t)P(t) + \alpha_p M(t) + \sqrt{\mu_p(t)P(t) + \alpha_p M(t)}\xi_p(t) \quad (5)$$

Where $M$ is HES5 mRNA concentration, $\mu_{m,p}$ are the degradation rate of HES5 mRNA and protein respectively, $\alpha_{m,p}$ are the transcription and translation rates of HES5, $\xi_{m,p}$ are Gaussian white noise, and $G(P(t-\tau_H))$ is an inhibitory Hill function that describes Hes5 autorepression

$$G(P(t-\tau_H)) = \frac{1}{1+\left(\frac{P(t-\tau_H)}{P_{H0}}\right)^{n_H}} \quad (6)$$

where $P_{H0}$ is the repression threshold for HES5 acting on its own promoter and $n_H$ is the corresponding Hill coefficient. The multiplicative nature of the two Hill functions in (4) is based on previous literature (Chikayama, 2014; Lewis, 2003)

In (4) and (5), there is a deterministic part which represents the overall increase or decrease in protein or mRNA to be expected at given concentrations and then a stochastic part that accounts for random binding or non-binding of transcription and translation machinery in the cell. The stochastic part is scaled by the square root of the number of events that can happen in a given time and so the stochastic part becomes increasingly significant at low protein/mRNA numbers. The Euler–Maruyama method was used to numerically calculate solutions to the SDDEs in (4) and (5) using a time step size of 1 minute.

Parameter values for simulations of the model included in **Figure 5** and **Appendix Figures S5** are given in **Appendix Table S3** marked as **Main**. We also investigated several other parameter combinations (**Appendix Table S3** C15/10/5-1 to 3 and **Appendix Figure S6**) previously identified to reproduce HES5 tissue data by Bayesian inference (Manning *et al.*, 2019).

##### Synthetic kymograph and spatial data generation

The 15μm strips used for spatial analysis in the experimental data (**Figure 2a**) were found to correspond to a width of 1-2 nuclei. Therefore, to obtain similar data from the model, spatial signals were generated by averaging 2 adjacent columns of cells. This single column of averaged data was used for generating each time point in the kymographs (**Figure 5g,h**) and also to test cluster occurrence and spatial periodicity (**Appendix Fig S5c,d** and **Appendix Fig S6a,b**).

##### Parameter space explorations and parameter selection

Parameter space 2D maps were used to visualise regions of different model behaviour (**Figure 5**, **Appendix Figure S54** and **Appendix Figure S6**). These parameter spaces have intercellular time delay plotted along x-axis and repression threshold value along the y-axis and at each point an output value of the model is plotted (e.g. temporal period). For each grid point in parameter space, the average output from 5 simulations with random initial conditions is shown. Parameter selection **(Figure 5f)** was used to identify suitable ranges for intercellular time delay and repression threshold by comparing statistics from synthetic data against experimental statistics obtained from tissue data whereby synthetic data values found within 2.4 standard deviations away from the experimental mean were accepted. Parameter selection was performed separately for temporal period and Kuramoto order parameter (see **Phase reconstruction**). Temporal period was computed from synthetic timeseries of HES5 as inverse of dominant frequency peak from power spectrum reconstruction by FFT. Temporal period used for parameter selection was average period per simulation where single cell period estimates above 10h were excluded as non-oscillatory.

##### Rate of differentiation from synthetic data

We explored how absolute sensing of HES5 by downstream targets might affect the rate of differentiation at different intercellular repression thresholds (coupling strengths) by assuming a simple linear differentiation condition where lower HES5 levels increase the probability of a cell committing to eventual differentiation (**Figure 6a**). This assumption is based on the fact that HES5 represses downstream pro-neural targets which promote a cell to head towards a differentiated state.

Because the level of expression changes as the coupling strength is varied, no set absolute threshold is assumed, instead a differentiation threshold is determined from the mean population expression in each simulation. Above this threshold, a cell has a zero chance of differentiation, whereas below the threshold has a linearly increasing probability of differentiation the further it drops below the threshold. A probability is calculated at each time point with no memory of past HES5 dynamics and is calculated as

$$P_{diff} = \begin{cases} 0, & P(t) > D_{thresh} \\ R\left(\dfrac{D_{thresh} - P(t)}{D_{thresh}}\right), & P(t) < D_{thresh} \end{cases} \quad (7)$$

where $R$ is the differentiation rate (with units of $\Delta t^{-1}$ as probability of differentiation is calculated on every time step), and $D_{thresh}$ is the differentiation threshold.

Cells are simply marked as having differentiated in this setup, and the dynamics of an individual cell remain the same after a differentiation event, which can be thought of an asymmetric division, where the differentiating cell leaves the population and it's space in the grid is filled with a non-differentiating cell. Differentiation rate is calculated by taking the total number of differentiation events in a simulation and dividing by the total time. Single cell parameters used are shown in **Appendix Table S3**, and the multicellular parameters used were $n_{ND} = 4$, $\tau_{ND} = 150$ minutes, and range of repression thresholds were explored $500 < P_{ND0} < 30000$.

**Synthetic microcluster occurrence and diameter calculation**

To detect presence of clusters of similar expressing cells in the synthetic data, spatial kymograph data was first smoothed using a polynomial fit. At each time point, the maximal peak in the polynomial was identified, followed by identification of the nearest trough below 80% the value of the peak which is consistent with a 1.2x spatial fold change in amplitude. The presence of a peak and trough satisfying these conditions is indicative of a cluster at a specific timepoint. The frequency of cluster occurrence was calculated as the number of time points in a simulation a cluster was identified.

The microcluster diameter was calculated from spatial data by searching for groups of cells that exhibit local in-phase characteristics. We performed phase reconstruction followed by pairwise phase synchronisation measurements by KOP (see Phase reconstruction) and counted the number of neighbouring cells in phase by filtering for KOPi,j >0.9 where I, j indicate neighbouring positions in the grid. In no-coupling conditions we estimated < 4% occurrence for clusters with diameter of 2 cells and <0.5% for diameter of 3. Thus microcluster diameters obtained under coupling conditions contain only a minute amount of false positives and diameters larger than 3 are not observed under no coupling.

#### References

Bennett H, Aguilar-Martinez E, Adamson, A.D. (2020). CRISPR-mediated knock-in in the mouse embryo using long single stranded DNA donors synthesised by biotinylated PCR. Methods. In Press. https://doi.org/10.1016/j.ymeth.2020.10.012

Eilers, P. H., & Goeman, J. J. (2004). Enhancing scatterplots with smoothed densities. Bioinformatics, 20(5), 623-628.

Gillespie, D. T. (2000). The chemical Langevin equation. The Journal of Chemical Physics, 113(1), 297-306.

Hodgkins A, Farne A, Perera S, Grego T, Parry-Smith DJ, Skarnes WC, Iyer V (2015) WGE: a CRISPR database for genome engineering. Bioinformatics 31: 3078 – 3080

Manning CS, Biga V, Boyd J, Kursawe J, Ymisson B, Spiller DG, Sanderson CM, Galla T, Rattray M, Papalopulu N (2019) Quantitative single-cell live imaging links HES5 dynamics with cell-state and fate in murine neurogenesis. Nature communications

Phillips NE, Manning CS, Pettini T, Biga V, Marinopoulou E, Stanley P, Boyd J, Bagnall J, Paszek P, Spiller DG et al (2016) Stochasticity in the miR-9/Hes1 oscillatory network can account for clonal heterogeneity in the timing of differentiation. Elife 5

Quadros, R.M., Miura, H., Harms, D.W. *et al. Easi*-CRISPR: a robust method for one-step generation of mice carrying conditional and insertion alleles using long ssDNA donors and CRISPR ribonucleoproteins. *Genome Biol* **18,** 92 (2017). https://doi.org/10.1186/s13059-017-1220-4
